# Quantifying the dynamics of viral recombination during free virus and cell-to-cell transmission in HIV-1 infection

**DOI:** 10.1101/2020.08.28.272351

**Authors:** Jesse Kreger, Josephine Garcia, Hongtao Zhang, Natalia L. Komarova, Dominik Wodarz, David N. Levy

## Abstract

Recombination has been shown to contribute to HIV-1 evolution *in vivo*, but the underlying dynamics are extremely complex, depending on the nature of the fitness landscapes and of epistatic interactions. A less well-studied determinant of recombinant evolution is the mode of virus transmission in the cell population. HIV-1 can spread by free virus transmission, resulting largely in singly infected cells, and also by direct cell-to-cell transmission, resulting in the simultaneous infection of cells with multiple viruses. We investigate the contribution of these two transmission pathways to recombinant evolution, by applying mathematical models to *in vitro* experimental data on the growth of fluorescent reporter viruses under static conditions (where both transmission pathways operate), and under gentle shaking conditions, where cell-to-cell transmission is largely inhibited. The parameterized mathematical models are then used to extrapolate the viral evolutionary dynamics beyond the experimental settings. Assuming a fixed basic reproductive ratio of the virus (independent of transmission pathway), we find that recombinant evolution is fastest if virus spread is driven only by cell-to-cell transmission, and slows down if both transmission pathways operate. Recombinant evolution is slowest if all virus spread occurs through free virus transmission. This is due to cell-to-cell transmission (i) increasing infection multiplicity, (ii) promoting the co-transmission of different virus strains from cell to cell, and (iii) increasing the rate at which point mutations are generated as a result of more reverse transcription events. This work further resulted in the estimation of various parameters that characterize these evolutionary processes. For example, we estimate that during cell-to-cell transmission, an average of 3 viruses successfully integrated into the target cell, which can significantly raise the infection multiplicity compared to free virus transmission. In general, our study points towards the importance of infection multiplicity and cell-to-cell transmission for HIV-evolution.

## Introduction

Human immunodeficiency virus-1 (HIV-1) infection eventually results in the development of AIDS, typically after several years. Viral evolution is thought to be a major contributor to disease progression, and poses important challenges to antiviral treatments and to the development of protective vaccines [1-3]. HIV-1 is characterized by a relatively high mutation rate (3×10^−5^ per base pair per generation) [4], which, together with the fast turnover of the virus population [5-7] contributes to the large evolutionary potential of the virus. Besides mutations, however, HIV-1 can also undergo recombination because the virions are diploid [8, 9]. If a cell is infected with two different virus strains and two different genomes are packaged into the offspring virus, recombination can occur between these two strains when the virus infects a new cell and undergoes reverse transcription. Recombination can accelerate the generation of 2-hit mutant viruses (virus strains with point mutations at two relevant locations) from two different 1-hit mutants (virus strains with a single relevant point mutation), which is likely faster than the generation of 2-hit mutants by point mutations alone. These processes can be especially important for the evolution of viral variants that simultaneously escape multiple immune cell clones or drugs.

While recombination has been shown to significantly contribute to viral evolution in vivo [9], mathematical models have demonstrated that the dynamics of recombination can be extremely complex [10-15]. Recombination can not only help the generation of multi-hit mutants, it can also break existing mutant combinations apart. The net effect of recombination is difficult to predict and depends on underlying assumptions about fitness landscapes, the nature and magnitude of epistatic interactions, and the relative balance of free virus and direct cell-to-cell transmission. Efforts have been made to quantify some of those processes, such as the fitness landscapes and the nature of epistasis in the evolution of drug resistance [16].

Here, we seek to quantify in detail how the different virus transmission pathways impact the evolution of recombinants. The nature of virus transmission during viral spread through the cell population is likely crucial for the rate of recombinant evolution, through variations in infection multiplicity [17-19]. Free virus transmission typically results in the infection of cells containing one virus, while direct cell-to-cell transmission through virological synapses typically results in the multiple infection of cells, and additionally is likely to result in the co-transmission of different virus strains from one cell to the next [18, 19]. Therefore, cell-to-cell (or synaptic) transmission is likely to be beneficial for the rate at which viral recombinants emerge.

Here, we combine mathematical models with *in vitro* experiments that utilize fluorescent reporter viruses [8] to test this hypothesis, to quantify the dynamical processes that lead to recombinant generation, and to estimate underlying parameters. Viruses can carry genes for cyan fluorescent protein (eCFP) and yellow fluorescent protein (eYFP). Recombination between these two variants results in viruses characterized by green fluorescence in target cells [8]. Additionally, recombination of the green fluorescent virus with a non-glowing virus can break the recombinant apart, resulting in yellow and cyan fluorescent viruses. The viral transmission pathway can be modulated by placing the culture on a gentle rocking platform [20, 21], which disrupts synaptic transmission. The frequency of infected cells displaying cyan, yellow and/or green is measured by flow cytometry over several days after infection. Mathematical models are applied to these experimental data to quantitatively characterize the dynamics.

## Materials and Methods

### Experimental setup

2×10^6^ CEM-SS cells were infected with 25µL NLENY1-IRES (YFP) or NLENC1-IRES (CFP), representing 19 virions/cell, by spinoculation at 1200g for 2 hours at 37°C as described previously [22]. Cells were cultured in Gibco Advanced RPMI-1640 with 5% FBS plus penicillin and streptomycin and 50 μM β-mercaptoethanol. The next day these cells were washed and YFP and CFP infected cells were mixed with uninfected CEM-SS cells.

Three independent experiments were performed at the following cell numbers in 3 ml of culture medium in T-75 flasks:

| Expt | Subset | YFP+CFP<br>$\times 10^6$ | Uninfected<br>$\times 10^6$ |
| --- | --- | --- | --- |
| 1, 2, 3 | A | 0.75 | 2 |
| 1, 2, 3 | B | 0.3 | 2 |
| 1, 2 | C | 0.075 | 2 |

These cultures were established in duplicate, one for the stationary condition and one for the rocking condition. The stationary set of cultures was placed on a shelf of a TC incubator at a 10° angle to concentrate the cells at one end. The rocking set was placed on a rocking platform in the incubator set at 12 tilts per minute. On each subsequent day 1/8-1/4 of the culture was collected for flow cytometry for YFP, CFP and GFP as previously described [8]. New culture medium was added to maintain a consistent volume in each culture. Details about the fluorescent reporter viruses used in this study can be found in [8].

### Mathematical models

We consider an ordinary differential equation (ODE) model of virus dynamics that tracks the populations of uninfected cells, as well as different types of infected cell populations. It incorporates both free virus and direct cell-to-cell transmission, and includes recombination processes. A basic schematic of the model is shown in Figure 1. Due to the complexity of the equations, they are displayed in the Supplementary Materials, Section 1. Basic model assumptions and their relation to the experimental data are summarized in the Results section. The different models are fit to the experimental data with standard methods, and the model that is most powerful at explaining the data is selected with the F-test for nested models. Details of the data fitting procedures are provided in the Supplementary Materials, Section 2.

**Figure 1.**
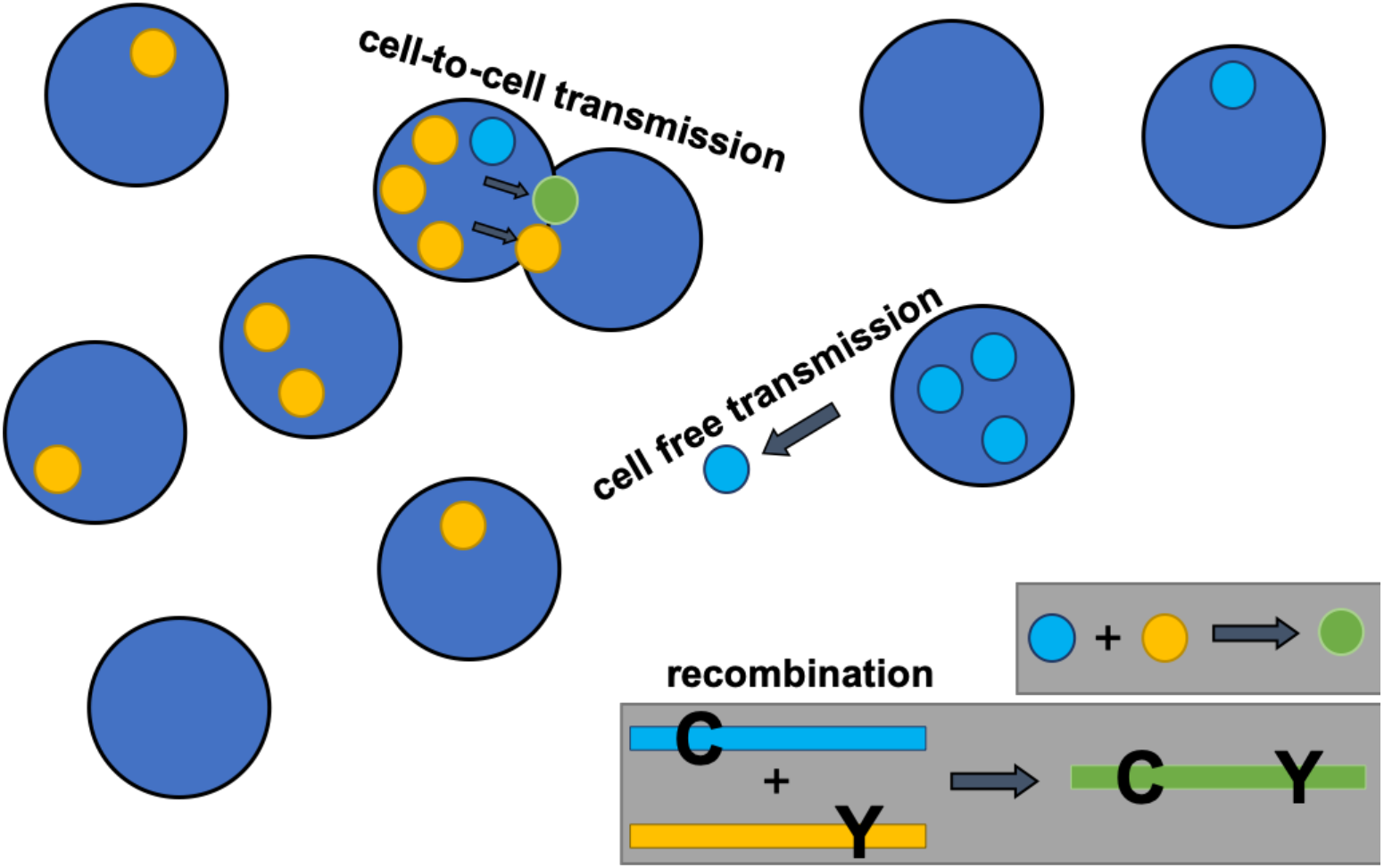
Basic model schematic. Cell free and synaptic cell-to-cell transmission are shown. In the ordinary differential equation model, both processes are non-spatial. An example of a recombination event is also shown.

## Results

### Basic dynamics of *in vitro* virus growth

The experimental system and the mathematical models are described in Materials and Methods, with additional details given in the Supplementary Materials. Based on previous work by others and us [20, 21], *in vitro* virus growth experiments were performed under two conditions: (i) under static conditions in which both free virus and synaptic transmission operate, and (ii) under shaking conditions where cultures were placed on a gentle rocking platform, which prevents most synaptic transmission events from taking place. For each experimental condition, eight experiments were performed that differed in viral inoculum size. Time-series were obtained for each condition, where we tracked cells infected with two types of single-hit mutant: those labeled with cyan fluorescent protein, denoted as C, and those labeled with yellow fluorescent protein, denoted as Y. In addition, we followed the population of cells infected with double-hit mutants labeled with green fluorescent protein, denoted as G, as well as populations of cells that contained combinations of virus types, such as YC, YG, CG, and YCG. Examples of this can be seen in Figures 2 (shaking condition) and 3 (static condition).

**Figure 2.**
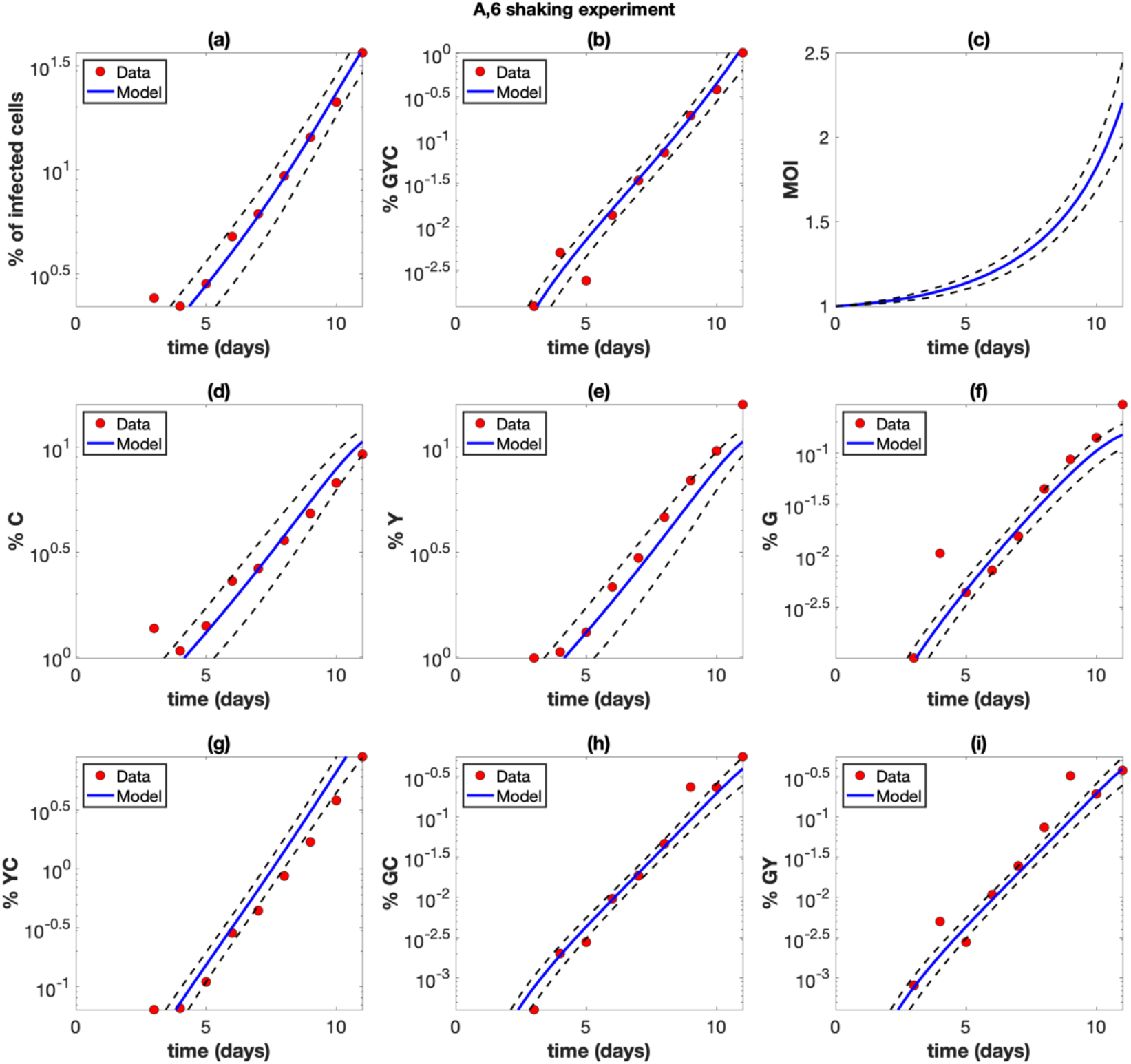
Example of a shaking experiment (experiment A6). The experimental data (red circles) are presented with best fit curves from the model (blue lines). The mathematical model is described in the Materials and Methods section with details in the Supplementary Materials, and the fitting procedure is described in the Supplementary Materials. Best fit parameters are included in Table 2 in the Supplementary Materials. The horizontal axis for all panels represents time (days). (a) The overall percentage of infected cells. (b) The percentage of cells infected with at least one copy of G, Y, and C. (c) The average multiplicity of infection (MOI) over all infected cells. (d) The percentage of cells infected with at least one active copy of C. (d) The percentage of cells infected with at least one active copy of Y. (f) The percentage of cells infected with at least one active copy of G. (g) The percentage of cells infected with at least one copy of Y and C. (h) The percentage of cells infected with at least one copy of G and C. (i) The percentage of cells infected with at least one copy of G and Y. The dashed black lines represent pointwise 95% prediction confidence bands.

As expected, exponential growth was observed for the total population of infected cells. This growth was faster under static compared to shaking conditions, resulting in higher virus levels for static conditions during the time frame of the experiment (Figure 3).

**Figure 3.**
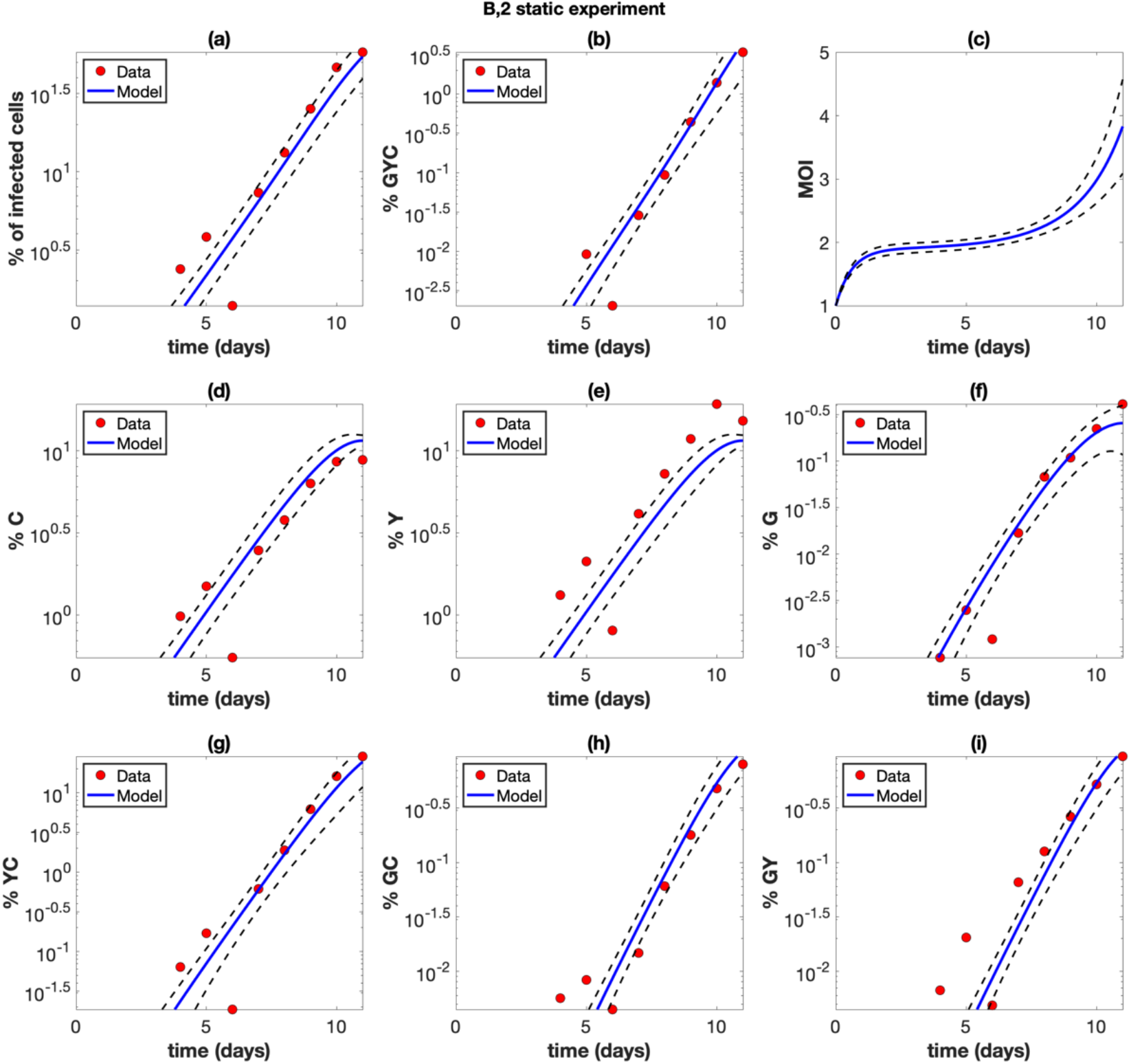
Example of a static, non-shaking experiment (experiment B2). The experimental data (red circles) are presented with best fit curves from the model (blue lines). Best fit parameters are included in Table 2 in the Supplementary Materials. The horizontal axis for all panels represents time (days). (a) The overall percentage of infected cells. (b) The percentage of cells infected with at least one copy of G, Y, and C. (c) The average multiplicity of infection (MOI) over all infected cells. (d) The percentage of cells infected with at least one active copy of C. (d) The percentage of cells infected with at least one active copy of Y. (f) The percentage of cells infected with at least one active copy of G. (g) The percentage of cells infected with at least one copy of Y and C. (h) The percentage of cells infected with at least one copy of G and C. (i) The percentage of cells infected with at least one copy of G and Y. The dashed black lines represent pointwise 95% prediction confidence bands.

### The mathematical model and data fitting

We used an extension of basic virus dynamics models [23, 24] that includes the multiple infection of cells as well as the occurrence of both free virus and synaptic transmission, based on our previous work [21, 25]. Due to the need to track reporter viruses that glow in multiple colors, the equations for the model are rather complicated and are described in detail in the Supplementary Materials, Section 1. Here, basic assumptions are summarized, and also shown schematically in Figure 1. Uninfected target cells are assumed to proliferate with a rate *r* and die with a rate *d*. Infected cells are generated through free virus transmission with a rate β, and through synaptic transmission with a rate γ. Free virus transmission results in the infection of the target cells with one virus. Synaptic transmission is assumed to lead to the simultaneous infection of the target cell with S copies of the virus, where the value of S is estimated from parameter fitting procedures. Infected cells are assumed to die with a rate *a*. Cells can be infected with (i) single-mutant viruses that carry cyan fluorescent protein, C, or yellow fluorescent protein, Y; (ii) the recombinant virus strain that shows green fluorescence (G); and (iii) the non-glowing virus strain that does not show any fluorescence, W. Hence, the model tracks cell populations that are infected with any combination of these virus strains, at defined multiplicities (i.e. cells can contain *i* copies of C, *j* copies of Y, *k* copies of G, and *l* copies of W viruses). The fitness of all virus types is assumed to be identical. During the infection process, the model assumes that recombination can occur with a certain probability if the infecting virus carries two genetically different genomes. All possible recombination events are described in the Supplementary Materials. The model is given by a set of ordinary differential equations and is hence non-spatial.

Besides these basic processes, an accurate model fit to all the data required the incorporation of two more processes into the model.

1. It has been demonstrated that infection can result in the generation of a latently infected cell, but that super-infection / multiple infection can result in the activation of the latent virus. The reason is that superinfection results in TAT complementation and thus activation of the latent genome in the cell [26]. For example, if a cell becomes infected with Y virus, it can become latent and the infected cell will not glow. Subsequent superinfection with C virus, however, will result in an infected cell that glows in both colors. As a consequence, the observed (glowing) population of singly infected cells is smaller than the actual population. This applies mostly to free virus transmission because it occurs mostly in singly-infected cells. Because we assume synaptic transmission to result in the infection of cells with S viruses (S>1), this effect can be ignored in this context. Hence, we assumed that upon infection of a cell with free virus, there is a probability ε that the infection becomes latent and that the virus consequently does not glow. Upon superinfection, however, we assume that the latent virus becomes activated and glows. Without this addition, the model under-predicts the double color YC cell numbers, and the model with latency can predict the YC population more accurately, as determined statistically by the F-test for nested models (Figure S1).
2. We found that for the shaking experiments, the model constructed so far consistently underpredicted multiply infected cells that contained the recombinant virus, i.e. the triple color GYC cells and double color cells GY and GC, although the number of cells infected with only GFP viruses (G) cells could be predicted more accurately (Figure S2). We hypothesized that although shaking results in the mixing of cells, the perfect mixing dynamics assumed by our ordinary differential equation model might be responsible for this discrepancy. In particular, we propose that when e.g. a YC-infected cell releases virus particles, they are likely to re-enter the same cell and generate a G virus upon recombination, thus explaining the higher than expected number of GYC cells. The reason is that as viruses are released from a cell, this cell is unlikely to immediately move away from its present location, but rather remains in the current vicinity for a while, which makes re-infection a likely event. Similarly, if a cell is infected with GW (where “W” stands for non-glowing virus) and only glows green, recombination between G and W can give rise to a C or a Y virus. Re-infection of the same cell with these viruses can then yield GCW and GYW cells (which glow in two colors). We first tested this idea with an agent-based model that tracked the spatial location of cells; preliminary explorations indicated that such a model could account for the data more accurately, but due to the complexity of the model and the corresponding data fitting procedures, we decided not to proceed with the agent-based model. Instead, we modified the ODE model to capture the assumption of a higher frequency of self-infection phenomenologically (see Supplementary Materials, Section 2). This model, while still compatible with a straightforward fitting procedure, could more accurately describe the GYC, GY, and GC data than the simpler model without this assumption (Figure S2), and was determined to be a more powerful model (despite containing an extra “self-infection” parameter) by the F-test for nested models.

These results indicate that the straightforward perfect mixing ODE models that are typically used to describe virus dynamics are not accurate descriptions of these *in vitro* dynamics at the level of detail presented here (this is discussed further below).

For data fitting, we simultaneously fit the model to the corresponding shaking and static experiments and to all infected subpopulations that were experimentally quantified. The detailed methodology is available in Section 2 of the Supplementary Materials. Some parameters were fixed according to information in literature, and these values are shown in Table 1 in the Supplementary Materials. The rest of the parameter values were estimated by model fitting, and the best fit parameters are given in Table 2 in the Supplementary Materials. The model fits are shown for select experiments in Figures 2 and 3, and for all experiments in the Supplementary Materials (Figures S13– S28). We note that the data are typically more noisy for the shaking compared to the static experiments, due to stochastic effects, especially if the experiments start with a low initial percentage of infected cells.

In what follows, we describe different parameter estimates and their implications for infection dynamics.

### Quantifying the relative contribution of free virus and synaptic transmission to virus spread

On the most basic level, we estimated the growth rate of the virus in the shaking and static conditions, and hence estimated the rate of free virus transmission and the rate of synaptic transmission. Previous work showed that shaking can increase the rate of free virus spread compared to static cultures due to mixing the virus more efficiently among the cells, by a factor of approximately f=1.33 [21]. Therefore, we corrected for this when calculating the rate of free virus spread from the shaking experiments (full details of the methodology are given in the Supplementary Materials). According to our calculations, the rate of free virus spread was on average 1.80±0.31 fold higher than the rate of synaptic transmission (see Section 3 in the Supplementary Materials). In our previous study, these rates were more equal, with the rate of free virus transmission being on average 1.1±0.1 fold higher than the rate of synaptic transmission [21]. This is further elaborated on in the Discussion section.

### Further parameter estimates

The model fitting to the experimental data allowed us to also estimate a variety of other parameters that characterize this system. (i) The probability for the virus to become latent upon single infection was rather consistent across the different experiments, with an average of ε=0.41± 0.084. In other words, about 40% of all infection events of uninfected cells (via free virus transmission) are estimated to result in viral latency in this system. (ii) We estimated the recombination probability to be ρ = 0.30±0.14. This is relatively close to the maximally possible recombination probability of ρ = 0.5 in this model (see Supplementary Materials). (iii) An important question concerns the number of viruses that are transferred from one cell to another through virological synapses during cell-to-cell transmission. Experimental data indicate that virus transfer through virological synapses is a very efficient process, with tens to hundreds of viruses transferred [17, 27]. Not all of these viruses, however, are likely to successfully integrate into the genome of the target cell. In our experiments, we estimated that on average S=3.0±0.35 viruses successfully infect a target cell per synaptic transmission event. (iv) The parameterized model further allowed us to estimate the average infection multiplicities in the experiments. As expected, the estimated multiplicities increased as the virus population grew. For the shaking experiments multiplicities ranged between 1 and 2.2 during the time frame of the experiments (Figure 2c). For the static experiments, the average infection multiplicity ranged from 1-4 during the phase of virus growth (Figure 3c).

### Comparing the effect of viral transmission pathways on recombinant generation: beyond the experimental setup

The experimental setup discussed above represents an artificial system to parameterize an evolutionary model of viral recombination. Here we use the parameterized model to simulate the dynamics in more general terms and to go beyond the experimental setup in the following ways.

1. Shaking conditions resulted in largely free virus transmission, while static conditions allowed both free virus and synaptic transmission to proceed. Using the estimated parameters in computer simulations, we further predict the dynamics for a scenario where only synaptic transmission takes place, which was not feasible experimentally.
2. The inherent differences in viral growth rates in the shaking and static experiments make a direct comparison of recombinant evolution under the different transmission pathways difficult to interpret. Hence, we adjusted the infection rates such that they are the same for free virus transmission only, synaptic transmission only, and a combination of the two.
3. The experimental setup is an artificially constructed system that was used to measure key parameters connected to recombinant evolution. We can use the parameterized equations to move beyond this particular system and describe a more complete evolutionary picture. That is, we can start with a “wild-type” virus (which was not part of the experimental setup) and model the generation of single-hit mutants by point mutation (which would technically correspond to the Y and C viruses, even though these reporter viruses are not generated by single mutation events in the experiments), and the subsequent evolution of double mutants through recombination (which would correspond to G viruses).

In the first set of simulations, we started with an equal number of single-mutant viruses, as done in the experiments, and ignored point mutations. We started the simulations with 10^9^ uninfected cells, and in contrast to the experiments, we did not allow for exponential expansion of the uninfected cell population, because lack of extensive cell expansion corresponds better to the conditions in which virus grows during acute HIV infection *in vivo*. We let the total number of infected cells grow until a population size of 4×10^8^ cells was reached. This was done assuming that only synaptic transmission (not possible in experiments), only free virus transmission, or both types of transmission occur. As a first step, we used unadjusted transmission parameters that were identical to the ones estimated from the experimental data. The number of cells that contain the recombinant virus was quantified as a function of time (Figure 4a). In these simulations, recombinants rise the fastest if both transmission pathways operate, compared to only a single active transmission pathway. This is largely due to both transmission types resulting in the fastest overall rate of viral transmission.

**Figure 4.**
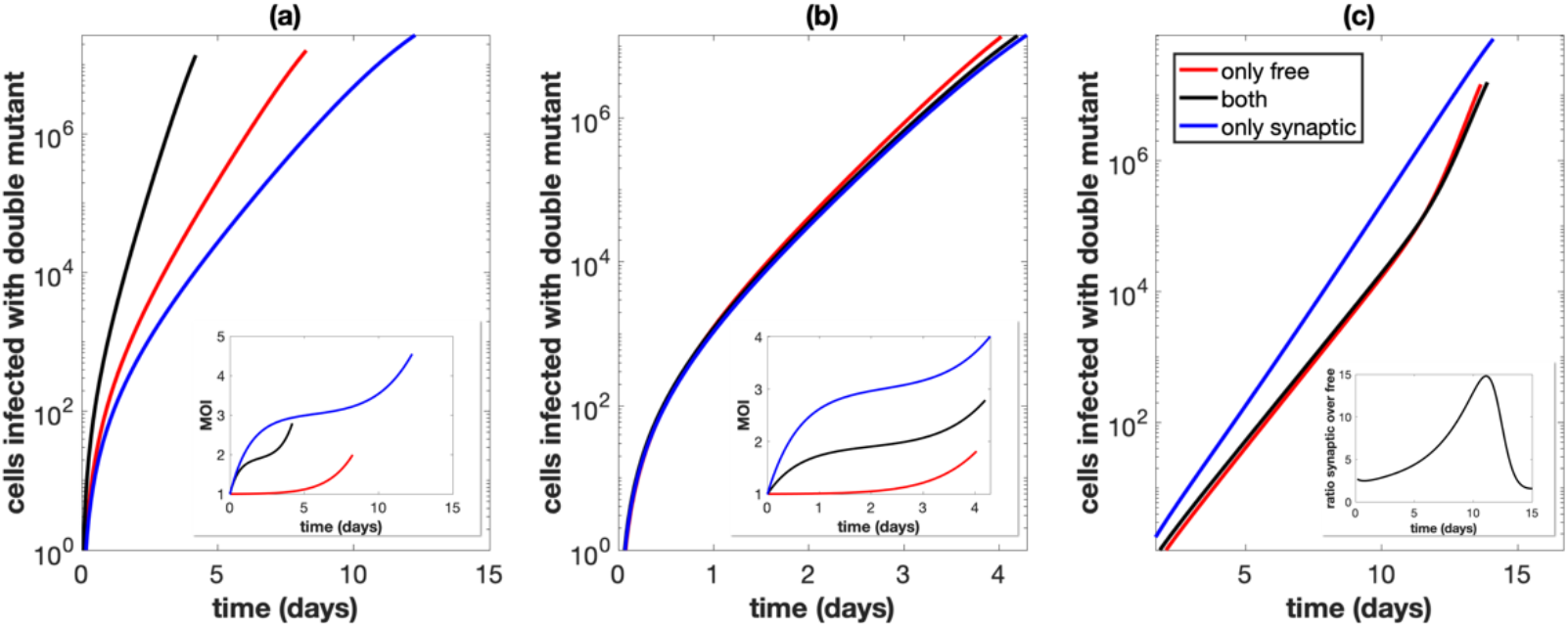
Total number of cells infected with at least one active copy of double mutant virus plotted against time. The multiplicity of infection (MOI) is included in the inset. We assume we have 10^9^ initial cells, and run the simulation until the number of infected cells reaches 40% of this initial amount, while also assuming that uninfected cells do not divide. Parameters are the best fit parameters from static experiment B2, which are listed in Table 2 in the Supplementary Materials. The combination of both transmission pathways is represented by the black lines. For the only free virus transmission case (red lines), synaptic transmission is turned off. For the only synaptic transmission case (blue lines), free virus transmission and reinfection are turned off. (a) Parameters are exactly as in the experiment. (b) The overall growth rate is the same across the three cases. (c) The overall growth rate is the same across the three cases and the simulation starts with only a single infected cell, which is coinfected with both single mutant strains.

To compare the effect of the transmission pathways themselves on recombinant generation (independent of the differences in viral transmission rates), we corrected for the differences in viral transmission rates associated with the different pathways, such that the basic reproductive ratio of the virus 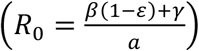was identical for free virus only, synaptic only, and for a combination of both transmission pathways (Figure 4b). When starting from the experimentally implemented initial conditions (infected cells that contain either one or the other virus type), we observe that the transmission pathways do not make a significant difference for the rate at which recombinants evolve. The reason is that we start with an infected cell population that contains either one or the other virus. Recombination requires the two virus types to come together in the same cell, which is likely to happen only at larger virus loads both for free virus and for synaptic transmission, explaining the lack of a difference in these simulations. Further, we note that at high virus loads, the number of recombinants in simulations with only free virus transmission slightly overtakes those in simulations with synaptic transmission; at this stage in the dynamics, the superinfection of latently infected cells becomes a common event in free virus transmission, which elevates the rate of productive free virus infection due to TAT complementation, and results in a higher effective reproductive number for the free-virus pathway compared to the synaptic pathway.

A different picture is observed if we start with a low number of coinfected cells that contain both types of single mutant virus (Figure 4c). Now, recombinant evolution is significantly faster for simulations with only synaptic transmission than for simulations that include free virus transmission. The reason is that synaptic transmission allows the repeated co-transmission of genetically different virus strains, which facilitates recombination processes in the growing virus population, highlighting the importance of this mechanism. Free virus transmission, in contrast, contributes to the separation of the two single mutant virus strains into separate cells, which slows downs recombinant generation.

Next, we consider a more complete evolutionary scenario. We start with a wild-type virus only, and introduce point mutations into the model. The wild-type virus can thus give rise to two different kind of single-mutant viruses, and double mutant viruses can be generated either by recombination between the single mutants, or by additional point mutations in those single mutants (Figure 5). We start with basic simulations (unadjusted parameters) and consider the number of cells that contain two different single mutant viruses over time, since these cells form the basis for recombination. Despite the fact that the estimated rate of synaptic transmission is slower than the estimated rate of free virus transmission, in our experimental system, the number of cells containing both single mutants rises sharply at low viral load in simulations that that take into account only synaptic transmission, compared to simulations that only take into account free virus transmission (Figure 5a). The reason is that (i) more reverse transcription events occur with synaptic transmission, thus generating more single mutant viruses (Figure 5a inset), and (ii) once the two different single mutant viruses have come together in the same cell, synaptic transmission enables their repeated co-transfer to target cells, explaining the sharp rise. As viral load grows to higher levels in the simulations, the number of cells coinfected with both single mutants under free virus transmission overtakes those under synaptic transmission (Figure 5a) because at high viral loads, the rate of coinfection becomes relatively high even for free virus transmission, and the estimated rate of free virus transmission is faster than the rate of synaptic transmission.

**Figure 5.**
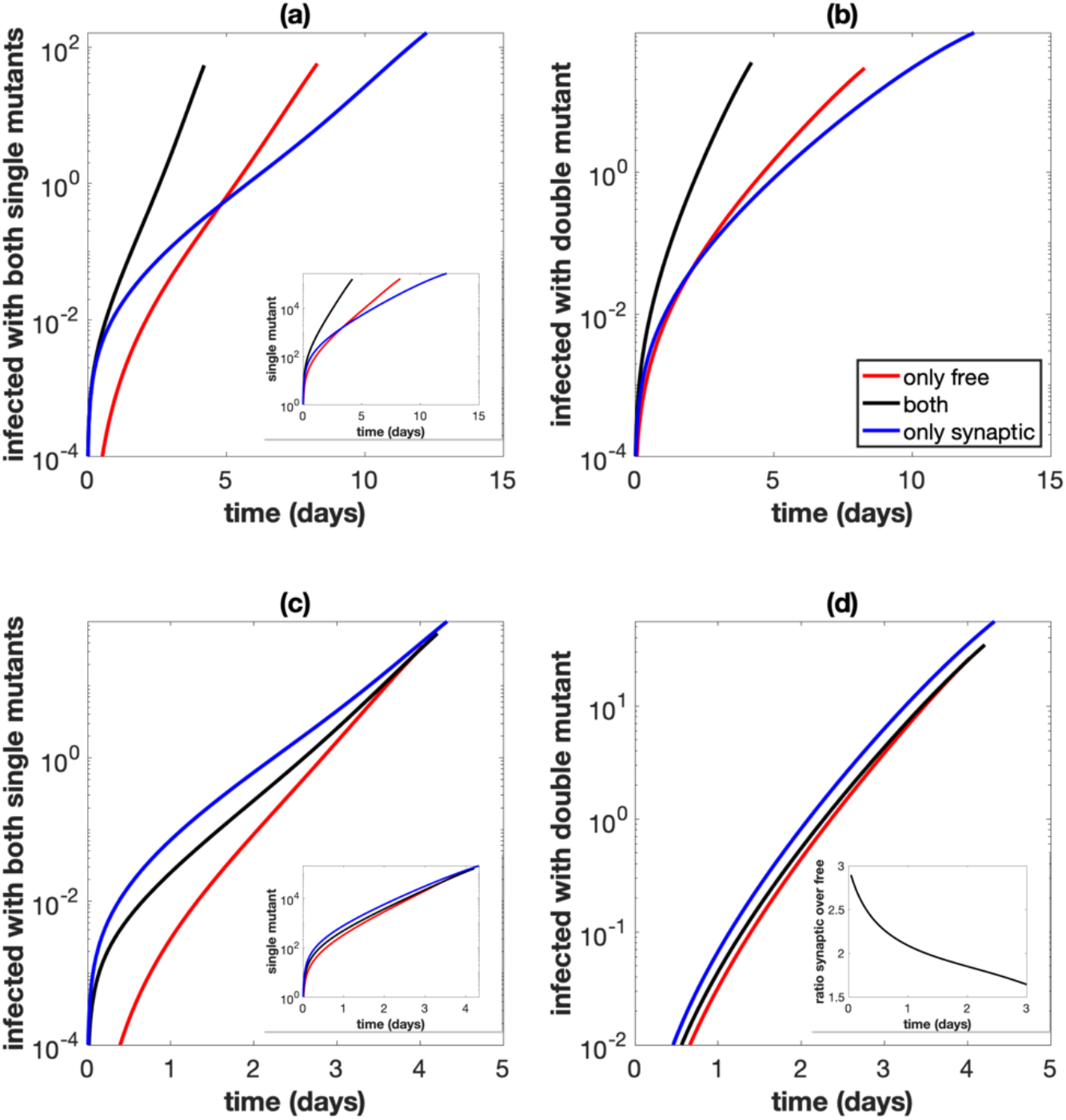
Same as in Figure 4, but the simulations start with a small equal amount of cells singly infected with the wild type, and mutations are included. (a) Total number of cells infected with at least one copy of both single mutant strains. The number of cells infected with at least one copy of one of the single mutants is included in the inset. Parameters are exactly as in the experiment. (b) Total number of cells infected with at least one active copy of double mutant virus plotted against time. Parameters are exactly as in the experiment. (c) Same as panel (a), but the overall growth rate is the same across the three cases. (d) Same as panel (b), but the overall growth rate is the same across the three cases. The ratio of the number of cells infected with the double mutant for only synaptic transmission versus only free virus transmission is included in the inset.

Looking at the number of double mutants (Figure 5b), we observe similar patterns. Figure 5c and d repeat these plots with adjusted parameters, such that the basic reproductive ratio of the virus is the same, independent of the transmission pathways that are assumed to occur. We see that the synaptic pathway contributes most to double mutant evolution, due to the repeated co-transmission of the two single-mutant strains. This is most pronounced when considering the number of cells that are coinfected with both single mutants (Figure 5c). The effect of synaptic transmission on the evolution of double mutants is qualitatively the same, but less pronounced because apart from recombination (Figure 5d), mutation processes also occur in these simulations. Therefore, although synaptic transmission significantly enhances the chances for recombination to occur (due to the co-transmission of the different single mutants), the mutation processes in the much more abundant singly infected cell population mask this effect to an extent.

## Discussion and Conclusion

In this study, we used mathematical models in combination with experimental data to determine the contribution of free virus transmission and direct cell-to-cell transmission to the evolution of recombinant viruses. This was possible through the use of experimental techniques to separate the two viral transmission pathways, and the use of fluorescent reporter viruses that allowed us to track recombinant evolution. Fitting of mathematical models to the experimental data allowed the estimation of important parameters that characterize the recombination process, and the models were further used to run evolutionary simulations to go beyond the experimental system and to quantify how the two transmission pathways influence the number of recombinants generated under a more complete set of evolutionary processes.

We found that direct cell-to-cell transmission through virological synapses promotes the evolution of recombinants, due to the following mechanisms: (i) synaptic transmission increases the infection multiplicity of cells, which is the basis for recombinant generation; (ii) synaptic transmission promotes the repeated co-transmission of two different virus strains from cell to cell, which increases the chances to eventually generate a recombinant virus; (iii) synaptic transmission increases point mutation generation because the simultaneous transfer of multiple viruses to the target cell increases the number of reverse transcription events, during which point mutations are mostly likely to occur. We estimated cell-to-cell transmission to result in the successful integration of about 3 viruses per synapse in the target cell. Experiments have shown that tens to hundreds of virus particles can be simultaneously transmitted per synapse [17, 27], but it is likely that only a subset of these result in successful integration.

We note that while for a fixed infection rate, purely synaptic transmission results in fastest recombinant evolution, free virus transmission can yield other advantages for the virus that might be equally important for its success, such as more efficient mixing of viruses among cells, which can promote virus spread and dissemination. The rate of recombinant evolution studied here is only one component that determines the success of the virus, and the presence of the two viral transmission pathways has to be interpreted in this light.

Our analysis also repeated some of our previous work [21] where we estimated the relative contribution of free virus and synaptic transmission to virus growth. In this study, it was estimated that the two transmission pathways contributed more or less equally to virus spread, where the rate of free virus transmission was approximately 1.1 fold faster than the rate of synaptic transmission. A subsequent study repeated our work and came to the same conclusion [28]. In the present study, we estimated the rate of free virus transmission to be approximately 1.8 fold faster than the rate of synaptic transmission, which is a larger difference. The reason for this discrepancy is that in the present study, we took into account the generation of latently infected cells upon free virus transmission, while the previous work did not. During synaptic transmission, latency is not established in our model because this is very unlikely due to the phenomenon of TAT complementation in multiply infected cells [26]. If we ignore free virus infection events that result in latency and re-calculate the rate of free virus transmission (see Section 3 in the Supplementary Materials), it is only about 1.06-fold faster than the rate of synaptic transmission, i.e. the two transmission pathways contribute about equally to virus spread, as in previous studies [21, 28]. Note that while the rate of productive free virus infection is the measure that is of immediate importance for the expansion of the virus population, the latently infected cells that are being generated can become relevant with a time delay if the virus spontaneously activates or becomes activated through TAT complementation upon superinfection. Hence, it is useful to consider both the total rate of free virus infection and the rate of productive infection.

Other work indicates that the relative importance of the two transmission pathways can depend on the microenvironment. For instance, in [29], HIV-1 spread in suspension was driven completely by free virus transmission, whereas virus spread in 3D collagen was driven by synaptic transmission, with approximately a 22% (36%) contribution of free virus transmission in loose (dense) collagen. This also brings up the effect of spatially restricted virus spread for the evolutionary dynamics explored here. The dynamics of virus growth under spatially restricted direct cell-to-cell transmission has been explored in the context of Hepatitis C virus infection [30] and more generally [31]. In the context of HIV infection, evidence for spatially clustered virus growth has been provided from experiments with HIV-infected humanized mice [18], and the consequences of spatially restricted synaptic transmission for the evolution through recombination has been studied recently with mathematical models [15]. It is unlikely that extensive spatial restrictions apply to the experimental virus growth cultures investigated here, in which the virus population grew exponentially rather than according to growth laws that are more typical for infected cell clusters. An extension of the currently described evolutionary dynamics to a spatial setting, however, will be important for future work.

An interesting finding in our study was that at the resolution of the data presented here, standard ordinary differential equations of virus dynamics failed to adequately describe several multiply infected cell subpopulations. We found that incorporating the assumption of limited mixing and the consequent re-infection of cells by their own offspring virus resulted in better model fits to the data. While this can be relevant to our *in vitro* system with transformed cell lines as infection targets, the relevance for virus replication *in vivo* might be less due to different infected cell life-spans and reverse transcription kinetics *in vivo* compared to the *in vitro* setting. Spatially restricted virus spread to new target cells (rather than self-infection), however, could have a similar effect since several viruses from the source cell likely get passed on to the same target cell due to the limited number of cells in the spatial neighborhood.

Our models and data analysis provide an important link between direct cell-to-cell transmission, heightened infection multiplicity, and an increased rate of recombinant evolution. While this was established within the framework of *in vitro* experimentation, these notions are likely also relevant *in vivo*. Synaptic transmission has been documented *in vivo* and infection multiplicities of cells in the tissue from HIV-infected patients have been shown to be between 3-4 or even higher on average [32]. Other studies reported the average infection multiplicities in the blood and tissue of HIV-infected patients to be closer to one [33, 34], although the restriction of the analysis to cells that express the CD4 receptor could have artificially lowered the estimate of infection multiplicities. The CD4 receptor becomes eventually down-regulated following the infection of the cell with the first virus. Ignoring such cells in the analysis could miss those cells with the highest infection multiplicities. The spatial modeling approaches discussed above might provide additional insights for these evolutionary dynamics *in vivo*. Future studies should further examine how these processes affect viral evolution under the assumption that mutants differ in fitness, and that the coinfection of cells with virus strains of different fitness can result in complementary and inhibitory interactions. The modeling framework and parameter estimates provided here form a basis for future investigation.

