## Supplementary Materials for "Quantifying the dynamics of viral recombination during free virus and cell-to-cell transmission in HIV-1 infection"

**Jesse Kreger[a], Josephine Garcia[c], Hongtao Zhang[c], Natalia L. Komarova[a], Dominik Wodarz[b,a] and David N. Levy[c]**

a) Department of Mathematics, University of California Irvine, Irvine CA 92697

b) Department of Population Health and Disease Prevention Program in Public Health Susan and Henry Samueli College of Health Sciences, University of California, Irvine CA 92697

c) Department of Basic Science, New York University College of Dentistry, New York NY 10010

---

#### Contents

**1 Mathematical model** **1**

**2 Fitting procedures and best fit parameters** **4**

**3 Relative contribution of free virus and synaptic transmission** **10**

**4 Role of synaptic transmission in evolution** **13**

**5 Further theoretical considerations** **16**

**6 Further justification of parameter estimates** **18**

**7 Best fit parameters and plots** **23**

**8 Comparing the effect of viral transmission pathways on recombinant dynamics for all best fit parameter values** **23**

---

#### 1 Mathematical model

Mathematical models have been used previously to describe the relative contribution of free virus and synaptic transmission to HIV-1 spread [1]. Here, we expand this model to better quantify multiple components of infection. In order to study the dynamics of recombinant viruses, this model can be expanded to include multiple infection and infection with multiple strains of virus. Here, based on the experimental setup, we consider single mutant strain C (equipped with glowing cyan fluorescent protein), single mutant strain Y (equipped with glowing yellow fluorescent protein), double mutant strain G (recombinant strain that includes both mutations and glows green), and double mutant strain W (non-glowing recombinant strain that includes neither mutation). We denote $x_{i,j,k,l}(t)$ as the number of cells infected with $i$ copies of strain C, $j$ copies of strain Y, $k$ copies of strain G, and $l$ copies of strain W. Further, we denote $\mathcal{N} = i + j + k + l$ as the total number of viruses that the cell is infected with. As $N$ is the maximum multiplicity of infection, $0 \leq \mathcal{N} \leq N$ must be satisfied.

Upon an infection event (either free virus or cell-to-cell), the infected cell produces a virus package that contains two copies of viral RNA. We assume that if recombination happens (with probability $0 \leq \rho \leq \frac{1}{2}$) that a distinct virus strain is created if possible (which is equivalent to not making this assumption and allowing $0 \leq \rho \leq 1$). Figure 1 in the main text shows a possible recombination event between strains Y and C to create recombinant strain G. We denote $P(C|i,j,k,l)$ to be the probability for a cell infected with $i$ copies of strain C, $j$ copies of strain Y, $k$ copies of strain G, and $l$ copies of strain W to infect with a copy of strain C upon an infection event. This can happen in the absence of recombination if at least one of the viral copies is of the C strain, and then that strain is chosen for infection. If one viral copy of G and one viral copy of W are chosen, and recombination occurs, then a C or Y virus is created. We assume that viral copies are chosen randomly based on their density within the cell. Therefore, we have that

$$P(C|i,j,k,l) = \frac{1}{\mathcal{N}^2}\Big(i(i+k+l) + (1-\rho)ij + \rho kl\Big). \tag{1}$$

Similarly, we have that

$$P(Y|i,j,k,l) = \frac{1}{\mathcal{N}^2}\Big(j(j+k+l) + (1-\rho)ij + \rho kl\Big), \tag{2}$$

$$P(G|i,j,k,l) = \frac{1}{\mathcal{N}^2}\Big(k(k+i+j) + (1-\rho)kl + \rho ij\Big), \tag{3}$$

$$P(W|i,j,k,l) = \frac{1}{\mathcal{N}^2}\Big(l(l+i+j) + (1-\rho)kl + \rho ij\Big). \tag{4}$$

Therefore, we have that the total intensity to infect with strain $\alpha$ is given by

$$B_\alpha = \sum_{0<\mathcal{N}\leq N} P(X|i,j,k,l,)x_{i,j,k,l} \quad \text{for } \alpha = C,\ Y,\ G,\ W, \tag{5}$$

where again $\mathcal{N}$ is the total number of copies of virus within the cell, so here we are sum over all infected subpopulations.

We also define $Z$ and $X$ as the total number of cells and total number of infected cells respectively, so we have that

$$Z = \sum_{0\leq\mathcal{N}\leq N} x_{i,j,k,l}, \tag{6}$$

$$X = \sum_{0<\mathcal{N}\leq N} x_{i,j,k,l}. \tag{7}$$

To include synaptic transmission, we first define $S \geq 1$ to be the constant number of viral copies transferred per synaptic infection event. Therefore, for each synaptic infection event, we must choose $S$ viral copies from the infected cell to pass on to the target cell. Assume that the infecting cell is infected with $i$ copies of strain C, $j$ copies of strain Y, $k$ copies of strain G, and $l$ copies of strain W. Synaptic transmission can result in infection by $\hat{i}$ copies of strain C, $\hat{j}$ copies of strain Y, $\hat{k}$ copies of strain G, and $\hat{l}$ copies of strain W, as long as $\hat{i}+\hat{j}+\hat{k}+\hat{l} = S$ is satisfied. The likelihood for each of these synaptic infection events is based on the probability to pick each strain for infection ($P(\alpha|i,j,k,l)$ for $\alpha$ = C, Y, W, and G). For each transmission event, we multiply the probability to transmit the respective number of each strain $P(C|i,j,k,l)^{\hat{i}}P(Y|i,j,k,l)^{\hat{j}}P(G|i,j,k,l)^{\hat{k}}P(W|i,j,k,l)^{\hat{l}}$ by the number of ways to transmit that number of each strain, which is the multinomial coefficient $\binom{S}{\hat{i},\hat{j},\hat{k},\hat{l}} = \frac{S!}{\hat{i}!\hat{j}!\hat{k}!\hat{l}!}$. Therefore, for $\hat{i}+\hat{j}+\hat{k}+\hat{l} = S$, we define the synaptic infection intensity as

$$G_{\hat{i},\hat{j},\hat{k},\hat{l}} = \frac{S!}{\hat{i}!\hat{j}!\hat{k}!\hat{l}!}\sum_{0<\mathcal{N}\leq N} P(C|i,j,k,l)^{\hat{i}}P(Y|i,j,k,l)^{\hat{j}}P(G|i,j,k,l)^{\hat{k}}P(W|i,j,k,l)^{\hat{l}}x_{i,j,k,l}. \tag{8}$$

Then the differential equation model is given by

$$\dot{x}_{0,0,0,0} = rx_{0,0,0,0} - \frac{\beta+\gamma}{Z} x_{0,0,0,0} X, \tag{9}$$

$$\begin{aligned} \dot{x}_{i,j,k,l} &= \frac{\beta}{Z}\left(x_{i-1,j,k,l}B_C + x_{i,j-1,k,l}B_Y + x_{i,j,k-1,l}B_G + x_{i,j,k,l-1}B_W - x_{i,j,k,l}X\right) \\ &+ \frac{\gamma}{Z}\left(\sum_{\hat{i}+\hat{j}+\hat{k}+\hat{l}=S} x_{i-\hat{i},j-\hat{j},k-\hat{k},l-\hat{l}} G_{\hat{i},\hat{j},\hat{k},\hat{l}} - x_{i,j,k,l}X\right) - ax_{i,j,k,l}, \end{aligned} \tag{10}$$

with the convention that any negative index is 0 and with the appropriate adjustments for maximum multiplicity of infection $N$. Here $r$ is the division rate of uninfected cells, $a$ is the death rate of infected cells, $\beta$ is the rate of free virus transmission, and $\gamma$ is the rate of synaptic transmission. Parameter descriptions (for all models) are given in Table S1. Note that the less detailed model used in [1] can be obtained by summing up the equations for all infected subpopulations. Since these are ordinary differential equations, the only difference between free virus and synaptic transmission is that multiple copies of virus can be transmitted during synaptic infection, as both processes are non-spatial.

**Latent infection.** It turns out (see Section 2) that the basic model described above is not sufficient to describe experimental data. As the next level of complexity, we also include latent infection in the description. We assume that upon infection, a cell that is only infected with a single copy of virus may not actively produce new virus. Once this cell is infected by additional copies of virus, it activates and starts glowing according to its viral contents. We denote silent cells with a single copy of a C or Y virus as $w_{1,0,0,0}$ and $w_{0,1,0,0}$ and silent cells with a single copy of a G or W virus as $w_{0,0,1,0}$ and $w_{0,0,0,1}$ respectively. We assume that latently infected cells grow according to the same law as the uninfected cells. The probability for an uninfected cell to become infected and not actively produce virus (become latent) is denoted by $\varepsilon$. Therefore, in the presence of latently infected cells, the total number of cells is given by

$$Z = \sum_{0 \le \mathcal{N} \le N} x_{i,j,k,l} + w_{1,0,0,0} + w_{0,1,0,0} + w_{0,0,1,0} + w_{0,0,0,1}. \tag{11}$$

The model with latent infection is then given by

$$\dot{x}_{0,0,0,0} = r x_{0,0,0,0} - \frac{\beta+\gamma}{Z} x_{0,0,0,0} X, \tag{12}$$

$$\dot{x}_{1,0,0,0} = \frac{\beta}{Z}\left(x_{0,0,0,0} B_C (1-\varepsilon) - x_{1,0,0,0} X\right) - \frac{\gamma}{Z}\left(x_{1,0,0,0} X\right) - a x_{1,0,0,0}, \tag{13}$$

$$\dot{w}_{1,0,0,0} = \frac{\beta}{Z}\left(x_{0,0,0,0} B_C\, \varepsilon - w_{1,0,0,0} X\right) - \frac{\gamma}{Z}\left(w_{1,0,0,0} X\right) + r w_{1,0,0,0}, \tag{14}$$

$$\dot{x}_{0,1,0,0} = \frac{\beta}{Z}\left(x_{0,0,0,0} B_Y (1-\varepsilon) - x_{0,1,0,0} X\right) - \frac{\gamma}{Z}\left(x_{0,1,0,0} X\right) - a x_{0,1,0,0}, \tag{15}$$

$$\dot{w}_{0,1,0,0} = \frac{\beta}{Z}\left(x_{0,0,0,0} B_Y\, \varepsilon - w_{0,1,0,0} X\right) - \frac{\gamma}{Z}\left(x_{0,1,0,0} X\right) + r w_{0,1,0,0}, \tag{16}$$

$$\ldots$$

$$\dot{x}_{2,0,0,0} = \frac{\beta}{Z}\left((x_{1,0,0,0} + w_{1,0,0,0}) B_C - x_{2,0,0,0} X\right) - \frac{\gamma}{Z}\left(x_{2,0,0,0} X\right) - a x_{2,0,0,0}, \tag{17}$$

$$\dot{x}_{0,2,0,0} = \frac{\beta}{Z}\left((x_{0,1,0,0} + w_{0,1,0,0}) B_Y - x_{0,2,0,0} X\right) - \frac{\gamma}{Z}\left(x_{0,2,0,0} X\right) - a x_{0,2,0,0}, \tag{18}$$

$$\dot{x}_{1,1,0,0} = \frac{\beta}{Z}\left((x_{1,0,0,0} + w_{1,0,0,0}) B_Y + (x_{0,1,0,0} + w_{0,1,0,0}) B_C - x_{1,1,0,0} X\right) - \frac{\gamma}{Z}\left(x_{1,1,0,0} X\right) - a x_{1,1,0,0}, \tag{19}$$

$$\ldots$$

$$\begin{aligned} \dot{x}_{i,j,k,l} &= \frac{\beta}{Z}\left(x_{i-1,j,k,l} B_C + x_{i,j-1,k,l} B_Y + x_{i,j,k-1,l} B_G + x_{i,j,k,l-1} B_W - x_{i,j,k,l} X\right) \\ &+ \frac{\gamma}{Z}\left(\sum_{\hat{i}+\hat{j}+\hat{k}+\hat{l}=S} x_{i-\hat{i},j-\hat{j},k-\hat{k},l-\hat{l}} G_{\hat{i},\hat{j},\hat{k},\hat{l}} - x_{i,j,k,l} X\right) - a x_{i,j,k,l}. \end{aligned} \tag{20}$$

again with the convention that any negative index is 0, with the appropriate adjustments for maximum multiplicity of infection $N$, and here also assuming $S \geq 3$ for simplicity of writing out the equations. For $S \leq 2$, subpopulations with $0 < \mathcal{N} \leq 2$ can also gain cells via infection by cell-to-cell transmission (with rate $\gamma$).

**A phenomenological reinfection term.** It turns out (see Section 2) that an additional layer of complexity is needed to describe the experimental data. In the experiments, spatial effects of infecting nearest neighbors are non-negligible. In order to incorporate these effects phenomenologically, we introduce "reinfection" terms in the equations. Namely, we assume that if a cell is infected with a double mutant virus, there is a rate at which it leaves that subpopulation and becomes infected with single mutants C and/or Y. Assuming that $k + l > 0$, we have that the model with the additional $\xi$ terms is given by

$$\begin{aligned} \dot{x}_{i,j,k,l} &= \frac{\beta}{Z}\left(x_{i-1,j,k,l} B_C + x_{i,j-1,k,l} B_Y + x_{i,j,k-1,l} B_G + x_{i,j,k,l-1} B_W - x_{i,j,k,l} X\right) \\ &+ \frac{\gamma}{Z}\left(\sum_{\hat{i}+\hat{j}+\hat{k}+\hat{l}=S} x_{i-\hat{i},j-\hat{j},k-\hat{k},l-\hat{l}} G_{\hat{i},\hat{j},\hat{k},\hat{l}} - x_{i,j,k,l} X\right) \\ &+ \xi\left(x_{i-1,j,k,l}/2 + x_{i,j-1,k,l}/2 - x_{i,j,k,l}\right) - a x_{i,j,k,l}, \end{aligned} \tag{21}$$

with the appropriate adjustments for latent infection (cells leave the latently infected with G and latently infected with W populations with rate $\xi$ and enter the appropriate coinfected subpopulation).

#### 2 Fitting procedures and best fit parameters

Based on previously estimated division and death rates for this cell type, we set $r = 0.924$ days$^{-1}$ and $a = 0.2$ days$^{-1}$ [1]. The death rate of infected cells is assumed lower for this in vitro cell

| Notation | Description | Units | Status |
|---|---|---|---|
| $r$ | division rate of uninfected cells | days$^{-1}$ | fixed $r = 0.924$ |
| $a$ | death rate of infected cells | days$^{-1}$ | fixed $a = 0.2$ |
| $\beta$ | rate of free virus transmission | days$^{-1}$ | fitted |
| $\gamma$ | rate of synaptic cell-to-cell transmission | days$^{-1}$ | fitted |
| $\xi$ | rate of recombinant reinfection | days$^{-1}$ | fitted |
| $S$ | number of viruses transmitted per synaptic infection event | NA | fitted |
| $N$ | maximum multiplicity of infection | NA | fixed $N = 20$ |
| $\varepsilon$ | latent infection probability | NA | fitted |
| $\rho$ | recombination probability | NA | fitted |
| $\eta$ | initial percentage of infected cells | NA | fitted |
| $P(\alpha \vert i, j, k, l)$ | probability of infection with strain $\alpha$ from $x_{i,j,k,l}$ cell | NA | NA |
| $Z$ | total number of cells | NA | NA |
| $X$ | total number of actively infected cells | NA | NA |
| $f$ | shaking factor | NA | fixed $f = 1.33$ |
| $\mu$ | mutation rate | NA | fixed $\mu = 3 \times 10^{-5}$ |

Table S1: Description of model parameters. Units are included if applicable.

line compared to estimates obtained from HIV-infected patients in vivo, and is consistent with previously published decline rates of HIV-infected cells in vitro [3]. We denote by $\mathcal{C}$ the percentage of cells infected with only strain C, by $\mathcal{YC}$ the percentage of cells infected with strain Y and strain C (but no other strains), and similarly with other strains, using calligraphic notation for the percentages. We denote $\mathcal{O}$ as the overall percentage of infected cells.

It is important to note that the double mutant recombinant strain W does not glow, and thus cannot be quantified experimentally. Therefore, we must make the appropriate adjustments to all glowing populations when comparing the experimental data to the mathematical model. For instance, experimental data for $\mathcal{C}$ correspond to mathematical model population $\mathcal{C} + \mathcal{CW}$ (as $\mathcal{CW}$ glows as if it were infected only with C) and experimental data for $\mathcal{O}$ correspond to mathematical population $\mathcal{O} - \mathcal{W}$. These adjustments are included in all steps below, where in terms of the mathematical model we redefine for instance $\mathcal{C} \coloneqq \mathcal{C} + \mathcal{CW}$, and similarly for all other populations.

We further note that (especially for static experiments), the population of infected cells can reach values very close to 100%. For experiments in which this occurs, we ignore all data points where the overall percentage of infected cells starts to decline, as the dynamics are no longer in growth phase (because there are no uninfected cells left to infect).

Based on the experimental setup, we assume that initially there is an equal percentage of cells singly infected with single mutant strain C and singly infected with single mutant strain Y, and that all other cells are uninfected. Since an equal amount of C and Y are initially infected, C and Y are symmetric from a mathematical perspective (as are G and W). Therefore, for the purposes of finding the best fits, we take averages of symmetric populations and use the following six populations for each experiment: $\mathcal{O}$, $(\mathcal{C} + \mathcal{Y})/2$, $\mathcal{YC}$, $\mathcal{G}$, $(\mathcal{GC} + \mathcal{GY})/2$, and $\mathcal{GYC}$.

While the experimentally measured sizes of these populations differ by orders of magnitude, we nonetheless would still like all of them to contribute to determining the parameters. To this end, we normalize the data so that the final data point is always 1 (by simply multiplying the time series for each population by 1 divided by the final data point in that particular time series). Finally, since the populations grow over orders of magnitude during the time-course of the experiments, we fit the natural log of the data (and thus ignore any 0 data points); this results in a more balanced fit where both early and late values matter.

All fits were completed using the *lsqcurvefit* function in Matlab, which calculates the parameters that minimize the sum of the squared difference between the experimental data and mathematical model over all experimental data points. Solutions of the differential equations were completed using *ode45*, which implements the Runge-Kutta method. We used random initial parameter guesses and repeated the procedure multiple times to ensure that global minima were obtained.

We began with the shaking experiment, since this simpler setting includes only free virus transmission. As the first step, we used the $\mathcal{O}$, $(\mathcal{C}+\mathcal{Y})/2$, and $\mathcal{YC}$ populations to fit the rate of free virus transmission, $\beta$, and the initial percent of infected cells, $\eta$, using the basic model (without latent infection) given by Equations (9-10). However, we found that it was necessary to include latent infection (model given by Equations (12-20)) because latent infection was needed for achieving a good fit for the coinfected $\mathcal{YC}$ cells. As illustrated in Figure S1, it is not possible to get a good fit for the single mutant populations ($\mathcal{Y}$, $\mathcal{C}$, and $\mathcal{YC}$) without latent infection (here for simplicity we ignored the small recombinant populations, set $\rho = 0$). We can see that the $\mathcal{YC}$ population is underestimated, and the $(\mathcal{C}+\mathcal{Y})/2$ and $\mathcal{O}$ populations are overestimated. Therefore, the basic model without latent infection and reinfection given by Equations (9-10) is not used in any steps of the data fitting.

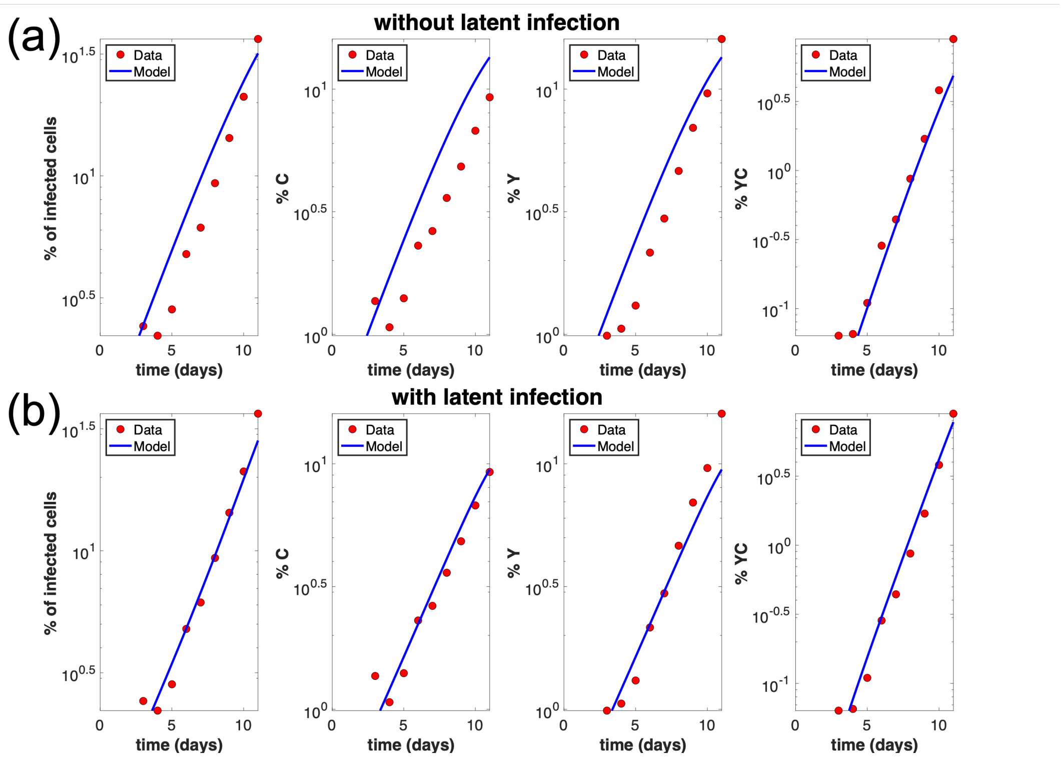


Figure S1: Comparison of best fits for shaking experiment A6 in the presence and absence of latent infection. Including latent infection results in a much better fit. The horizontal axis represents time (days) and the vertical axis represents the percentage of cells in the given category. (a) Best fit without latent infection. Here we use the mathematical model without latent infection given by Equations (9-10), and we see that the percent of cells coinfected with both Y and C is an underestimate, and the percent of cells infected with only Y (and only C) is an overestimate. (b) Best fit with latent infection. Here we use the mathematical model with latent infection given by Equations (12-20). Parameters for this fit are given in Table S3. Here get an F-statistic of $\mathcal{F} = 16.4$ and p-value of 0.007.

We can use the F-test for nested model to demonstrate the necessity of the additional level of complexity by including latent infection (and thus an additional parameter $\varepsilon$). For the experiment shown in Figure S1, we get an F-statistic of $\mathcal{F} = 16.4$, where

$$\mathcal{F} = \frac{\frac{R_1 - R_2}{p_2 - p_1}}{\frac{R_2}{n - p_2}} \tag{22}$$

with $n$ the number of data points, $p_i$ the number of parameters, and $R_i$ the sum of the squared errors in the respective models. Here the model with subscript 1 is the nested model without latent infection given by Equations (9-10) and the model with subscript 2 is the model with latent infection given by Equations (12-20). The null hypothesis is that the more complex model does not provide a significantly better fit than the reduced nested model. The F-statistic is a ratio of two quantities that are expected to be roughly equal under the null hypothesis, which produces an F-statistic of approximately 1. Here the F-statistic is 16.4 and p-value is 0.007, and so we reject the null hypothesis and use the more complex model with latent infection.

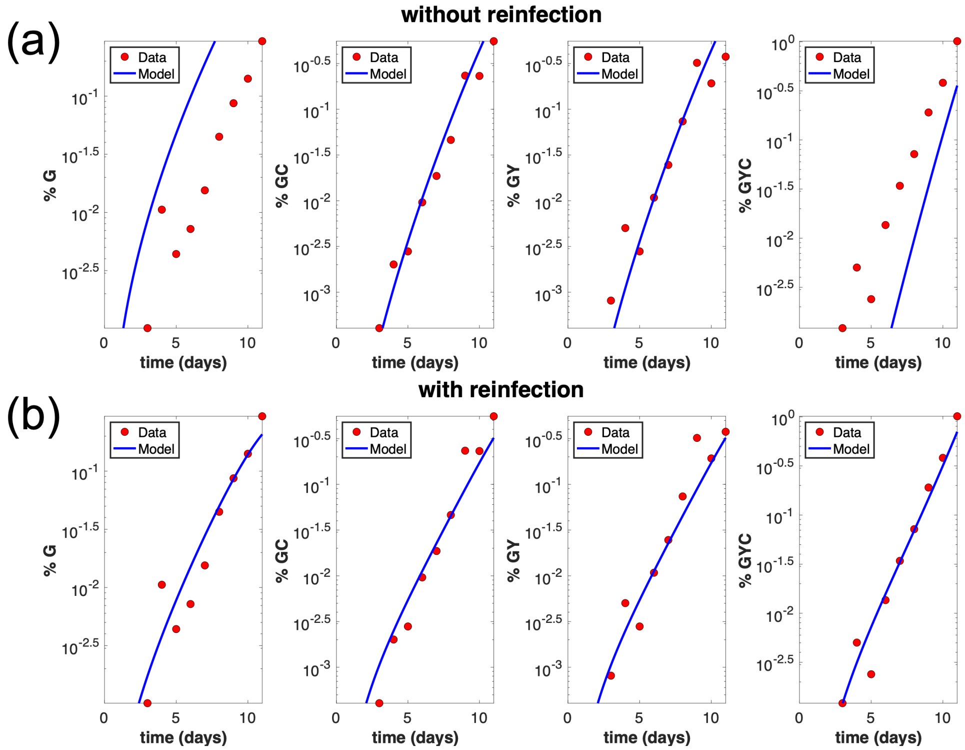


Figure S2: Comparison of best fits for shaking experiment A6 in the presence and absence of reinfection. Including reinfection results in a much better fit. (a) Best fit without reinfection, system (12-20): the percent of cells coinfected with G, Y, and C is an underestimate, and the percent of cells infected with only G is an overestimate. (b) Best fit with reinfection, system (21). Parameters for this fit are given in Table S3. F-statistic is $\mathcal{F} = 39.3$ and p-value of 0.002.

The next step involved fitting the recombinant populations of $\mathcal{G}$, $(\mathcal{GC} + \mathcal{GY})/2$, and $\mathcal{GYC}$ to estimate the recombination probability, $\rho$. However, using the model with latent infection but without reinfection, which are given by Equations (12-20), resulted in severe underestimation of $\mathcal{GYC}$ cells, see Figure S2. This is because of the tendency of newly created recombinant G viruses to reinfect the cells that they are produced from (which must contain at minimum a copy of C and a copy of Y). This spatial effect results in higher experimentally observed fractions of $\mathcal{GYC}$. Therefore, we included phenomenological reinfection terms in the ODE model. If these terms are not included, the $\mathcal{GYC}$ population will be an underestimate, and the percent of cells infected with only $\mathcal{G}$ will be an overestimate, see Figure S2. This model includes a phenomenological reinfection term $\xi$ for recombinant strains G and W, is given in Equation (21). Again, we can use the F-test for nested model to demonstrate the necessity of the additional level of complexity by including reinfection (and thus an additional parameter $\xi$). For the experiment shown in Figure S2, we get an F-statistic of $\mathcal{F} = 39.3$, where the nested model without reinfection is given by Equations (12-20) and the full model with reinfection is given by Equation (21). This gives a p-value of 0.002, and so we reject the null hypothesis and use the more complex model with reinfection.

Below we describe two fitting procedures that we implemented, and which are based on the above modeling work. The first procedure is what we called a **simultaneous fitting method**, which provides an overall lower error. The second procedure is the **sequential fitting method**, which gives similar parameter estimates (with a slightly larger overall error), but is a much faster algorithm. Additional justification for parameter estimates is given in Section 6.

**Simultaneous fitting method.** Each experimental pair includes a shaking experiment (only free virus transmission) and a static experiment (both free virus and cell-to-cell transmission). This method uses a corresponding shaking and static experiment and fits them simultaneously for all parameters using the data for $(\mathcal{C} + \mathcal{Y})/2$, $\mathcal{YC}$, $\mathcal{G}$, $(\mathcal{GC} + \mathcal{GY})/2$, and $\mathcal{GYC}$. Therefore, we have ten data time series (five from each experiment) and fit them for eight parameters ($\beta$, $\varepsilon$, $\xi$, $\eta_{shaking}$, $\eta_{static}$, $\rho$, $\gamma$, and $S$) using the most complex model with latent infection and reinfection, which is given by Equation (21). For the shaking experiment, we set $\gamma = 0$ as there is only free virus transmission (with rate $\beta$). For the corresponding static experiment, we allow $\gamma > 0$ and simultaneously fit the rate of free virus transmission $\beta/f$, where $f$ is the shaking factor. This is because shaking increases the efficiency of free virus infection on average by a factor of $f = 1.33$ [1].

This method eliminates the possibility of finding globally sub-optimal fits that exists with the sequential fitting procedure described below. However, this comes at a great computational cost, as each individual experiment (with four virus strains and maximum multiplicity of infection $N = 20$) involves $\binom{20+4}{4} = 10,626$ cell supopulations and differential equations. Fitting two experiments simultaneously for all parameters is thus computationally expensive, and can take several hours per fit.

There are eight pairs of experiments (eight shaking experiments and eight corresponding static experiments). The experiments are spilt into three subsets. Subset A contains three static experiments (A1,A2,A3) and three corresponding shaking experiments (A4,A5,A6). Here, A1 and A4 have the same initial conditions, A2 and A5 have the same initial conditions, and A3 and A6 have the same initial conditions. Similarly, subset B contains three static experiments (B1,B2,B3) and three corresponding shaking experiments (B4,B5,B6), and subset C contains two static experiments (C1,C2) and two corresponding shaking experiments (C3,C4). The best fit parameters using the simultaneous fitting procedure, along with the 95% parameter confidence intervals (based on an asymptotic normal distribution for the parameter estimate), are included in Table S2. Here for all experiments the shaking factor $f = 1.33$, maximum multiplicity of infection $N = 20$, $r = 0.924$, and $a = 0.2$.

**Sequential fitting method.** A less computationally intensive sequential fitting method is also possible. Rather than the straightforward fitting of all data for all parameters for a pair of corresponding experiments, each step of the hierarchical fitting procedure below describes which infected

| experiment | $\beta_{\text{shaking}}$ | $\varepsilon$ | $\xi$ | $\eta_{shaking}$ | $\eta_{static}$ | $\rho$ | $\gamma$ | $S$ |
|---|---|---|---|---|---|---|---|---|
| A4/A1 | $2.56 \pm 0.66$ | $0.50 \pm 0.15$ | $1.99 \pm 1.06$ | $0.0022 \pm 0.0010$ | $0.00012 \pm 0.000089$ | $0.23 \pm 0.17$ | $1.23 \pm 0.16$ | 4 |
| A5/A2 | $2.57 \pm 0.53$ | $0.48 \pm 0.13$ | $2.3 \pm 0.96$ | $0.0034 \pm 0.0012$ | $0.00061 \pm 0.0030$ | $0.18 \pm 0.10$ | $1.05 \pm 0.11$ | 3 |
| A6/A3 | $2.35 \pm 0.30$ | $0.39 \pm 0.10$ | $3.06 \pm 0.96$ | $0.0054 \pm 0.0013$ | $0.00062 \pm 0.00030$ | $0.19 \pm 0.073$ | $1.19 \pm 0.11$ | 3 |
| B4/B1 | $1.97 \pm 0.51$ | $0.39 \pm 0.17$ | $1.18 \pm 0.67$ | $0.0027 \pm 0.0015$ | $0.00047 \pm 0.00029$ | $0.39 \pm 0.31$ | $0.78 \pm 0.10$ | 3 |
| B5/B2 | $2.19 \pm 0.48$ | $0.45 \pm 0.15$ | $1.76 \pm 0.77$ | $0.0054 \pm 0.0023$ | $0.0015 \pm 0.00088$ | $0.20 \pm 0.13$ | $0.74 \pm 0.10$ | 3 |
| B6/B3 | $2.15 \pm 0.33$ | $0.43 \pm 0.12$ | $1.79 \pm 0.67$ | $0.0090 \pm 0.0032$ | $0.0027 \pm 0.0013$ | $0.20 \pm 0.095$ | $0.73 \pm 0.088$ | 3 |
| C3/C1 | $1.83 \pm 0.55$ | $0.37 \pm 0.22$ | $0.49 \pm 0.35$ | $0.0045 \pm 0.0026$ | $0.00084 \pm 0.00053$ | $0.49 \pm 0.39$ | $0.80 \pm 0.12$ | 3 |
| C4/C2 | $1.56 \pm 0.42$ | $0.23 \pm 0.24$ | $0.97 \pm 0.67$ | $0.0079 \pm 0.0044$ | $0.0014 \pm 0.00094$ | $0.50 \pm 0.40$ | $0.82 \pm 0.13$ | 3 |

Table S2: Best fit model parameters using the simultaneous fitting method (mathematical model described in Section 1 and fitting procedure described in Section 2) for each pair of experiments. For the shaking experiments, $\gamma = 0$. For the static experiments, $\beta_{\text{static}} = \beta_{\text{shaking}}/f$, where $f$ is the shaking factor. Parameter descriptions and units can be found in Table S1.

cell populations were used to fit which parameters.

We start with the shaking experiment, which only contains free virus transmission. We begin with the $\mathcal{O}$, $(\mathcal{C} + \mathcal{Y})/2$, and $\mathcal{YC}$ populations (set $\rho = \xi = 0$), to fit the rate of free virus transmission, $\beta$, the probability of latent infection, $\varepsilon$, and the initial percentage of infected cells, $\eta$ using the mathematical model with latent infection given by Equations (12-20). We then fix these three parameters ($\beta$, $\varepsilon$, and $\eta$) and use the remaining populations of $\mathcal{G}$, $(\mathcal{GC} + \mathcal{GY})/2$, and $\mathcal{GYC}$ to fit the recombination probability, $\rho$, and rate of recombinant reinfection, $\xi$, using the mathematical model with reinfection given by Equation (21). While $\rho$ and $\xi$ affect the non-recombinant populations, we assume this effect is negligible and so it is included in a separate step after the other parameters have been fixed.

For the corresponding static experiment, we fix the rate of free virus transmission to be $\beta/f$ and further we fix $\varepsilon$ and $\xi$ as what was obtained in the shaking experiment. We then use only the $\mathcal{O}$ population to fit the rate of synaptic transmission, $\gamma$, and the initial percentage of infected cells, $\eta$. We then finally use the recombinant populations of $\mathcal{G}$, $(\mathcal{GC} + \mathcal{GY})/2$, and $\mathcal{GYC}$ to fit the number of viral copies transferred per synapse, $S$. We use these populations because changing $S$ has the most significant effect on the smaller recombinant populations. Increasing $S$ increases the number of coinfection events, which increases the percentage of cells infected with $GYC$ and decreases the percentage of cells infected with $G$. For all steps in fitting the static experiment, we use the full model with reinfection.

The best fit parameters for this method are included in Table S3 and Table S4. The 95% parameter confidence intervals, calculated at the relevant step of the fitting, are also included.

| | $\beta$ | $\varepsilon$ | $\xi$ | $\eta$ | $\rho$ | corresponding static experiment |
|---|---|---|---|---|---|---|
| experiment A4 | 2.23±0.58 | 0.40±0.16 | 1.44±1.0079 | 0.0017±0.00083 | 0.31±0.14 | experiment A1 |
| experiment A5 | 1.94±0.25 | 0.28±0.10 | 2.90±1.30 | 0.0037±0.0010 | 0.35±0.10 | experiment A2 |
| experiment A6 | 1.99±0.15 | 0.28±0.063 | 2.43±1.055 | 0.0072±0.0011 | 0.28±0.077 | experiment A3 |
| experiment B4 | 1.63±0.24 | 0.22±0.12 | 1.27±0.91 | 0.0026±0.0010 | 0.36±0.17 | experiment B1 |
| experiment B5 | 1.60±0.20 | 0.21±0.11 | 1.93±1.23 | 0.0081±0.0027 | 0.34±0.14 | experiment B2 |
| experiment B6 | 1.58±0.20 | 0.20±0.12 | 1.90±0.92 | 0.017±0.0058 | 0.31±0.096 | experiment B3 |
| experiment C3 | 1.56±0.22 | 0.20±0.12 | 0.77±0.63 | 0.0034±0.0011 | 0.49±0.27 | experiment C1 |
| experiment C4 | 1.50±0.28 | 0.16±0.16 | 1.13±0.72 | 0.0069±0.0031 | 0.40±0.17 | experiment C2 |

Table S3: Best fit model parameters using the sequential fitting method for each shaking experiment. Parameter descriptions can be found in Table S1.

| | $\beta$ | $\varepsilon$ | $\xi$ | $\eta$ | $\rho$ | $\gamma$ | $S$ |
|---|---|---|---|---|---|---|---|
| experiment A1 | $2.23/f$ | 0.40 | 1.44 | 0.00021±0.00010 | 0.31 | 1.12±0.067 | 4 |
| experiment A2 | $1.94/f$ | 0.28 | 2.90 | 0.00067±0.00010 | 0.35 | 1.06±0.037 | 3 |
| experiment A3 | $1.99/f$ | 0.28 | 2.43 | 0.0018±0.00041 | 0.28 | 1.01±0.048 | 2 |
| experiment B1 | $1.63/f$ | 0.22 | 1.27 | 0.00022±0.00011 | 0.36 | 0.86±0.064 | 3 |
| experiment B2 | $1.60/f$ | 0.21 | 1.93 | 0.0013±0.0010 | 0.34 | 0.78±0.012 | 3 |
| experiment B3 | $1.58/f$ | 0.20 | 1.90 | 0.0037±0.0011 | 0.31 | 0.72±0.048 | 3 |
| experiment C1 | $1.56/f$ | 0.20 | 0.77 | 0.00036±0.00020 | 0.49 | 0.88±0.078 | 3 |
| experiment C2 | $1.50/f$ | 0.16 | 1.13 | 0.00067±0.00038 | 0.40 | 0.91±0.091 | 3 |

Table S4: Best fit model parameters using the sequential fitting method for each corresponding static experiment. Parameter descriptions can be found in Table S1.

Since the simultaneous fitting procedure results in more globally optimal fits, we use its parameters (Table S2) for all figures and calculations in both the main text and Supplementary Materials. While the parameter estimates for the less optimal sequential fitting procedure described below are not identical, the overall error is similar, with the sequential fitting method generally resulting in less error in the shaking experiment and the simultaneous fitting method generally resulting in less error in the static experiment. The best fit curves also look similar, as seen in Figures S3 and S4 which shows the comparison of the two fitting methods along with the experimental data.

### 3 Relative contribution of free virus and synaptic transmission

Using the simultaneous fitting parameters found in the previous section (Table S2), we estimate the relative contribution of free virus transmission compared to synaptic transmission in driving infection. In contrast to our previous study [1], we now include latent infection (which is described further in Section 1). In order to compare relative contributions of the synaptic and free virus transmission pathways, we note that the rate of free virus transmission, $\beta$ is on average $1.80 \pm 0.31$ fold higher than the rate of synaptic transmission (Table S5, column 5).

However, it is important to note that the presence of latent infection modifies the transmission process. In fact, the rate of productive infection (at least, relatively early in the infection process) is better characterized by $\beta(1-\varepsilon)$. Taking this into account we obtain that the rate of free virus spread is on average $1.06 \pm 0.16$ fold higher than the rate of synaptic transmission (Table S5, column 6). This is very close to the estimated ratio of 1.1 in [1].

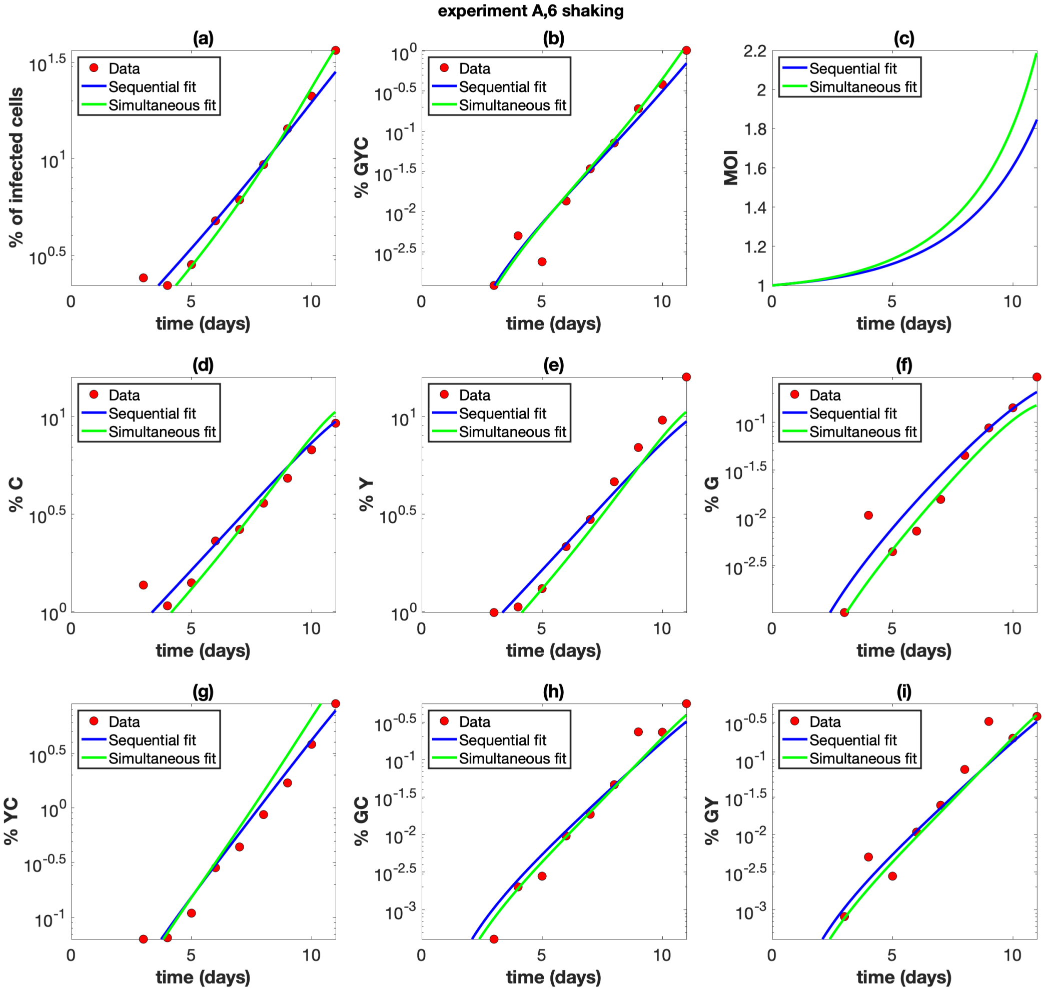


Figure S3: Comparison of the sequential fitting method (blue lines) with the simultaneous fitting method (green lines) for shaking experiment A6. The experimental data are also shown (red dots). The parameters for each fit are given in Tables S3 and S2. (a) The overall percentage of infected cells. (b) The percentage of cells infected with at least one copy of G, Y, and C. (c) The average multiplicity of infection (MOI) over all infected cells. (d) The percentage of cells infected with at least one copy of C. (e) The percentage of cells infected with at least one copy of Y. (f) The percentage of cells infected with at least one copy of G. (g) The percentage of cells infected with at least one copy of Y and C. (h) The percentage of cells infected with at least one copy of G and C. (i) The percentage of cells infected with at least one copy of G and Y.

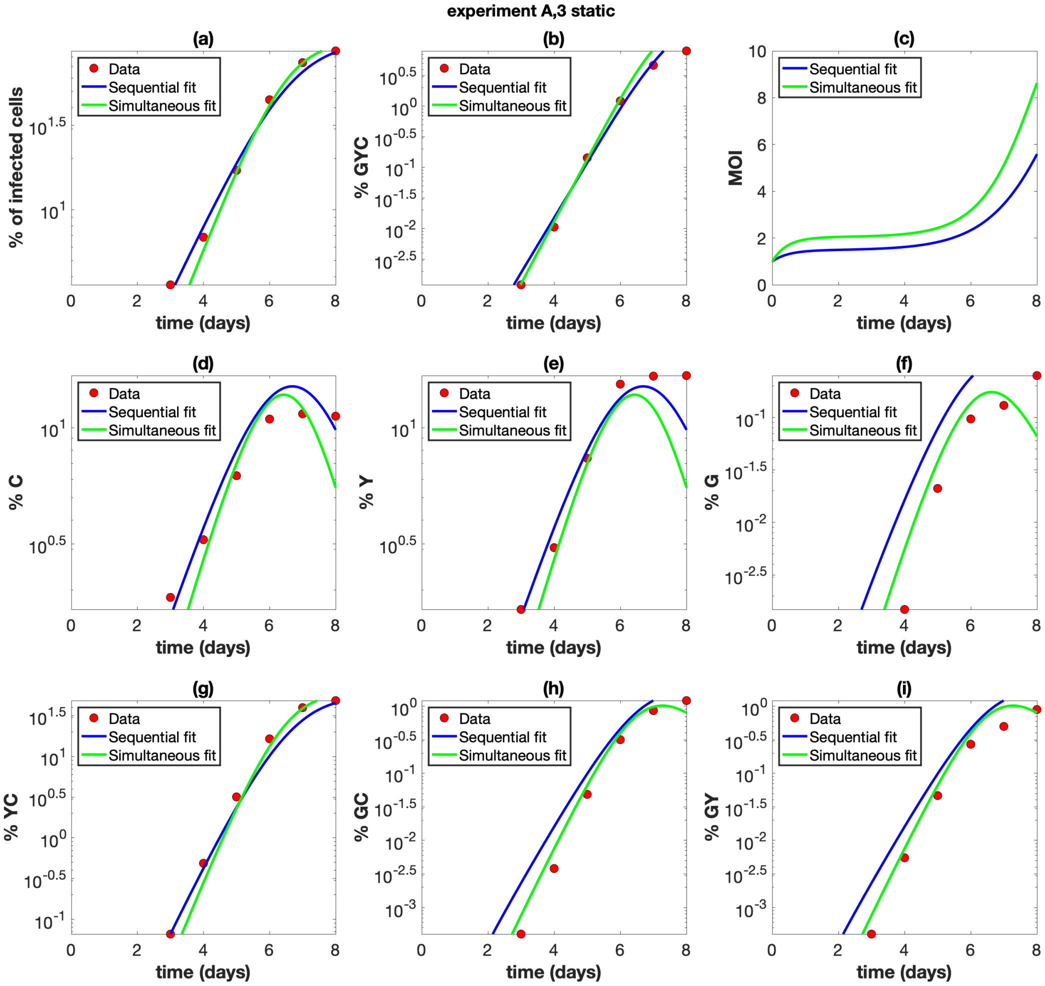


Figure S4: Comparison of the sequential fitting method (blue lines) with the simultaneous fitting method (green lines) for static experiment A3. The experimental data are also shown (red dots). The parameters for each fit are given in Tables S4 and S2. (a) The overall percentage of infected cells. (b) The percentage of cells infected with at least one copy of G, Y, and C. (c) The average multiplicity of infection (MOI) over all infected cells. (d) The percentage of cells infected with at least one copy of C. (e) The percentage of cells infected with at least one copy of Y. (f) The percentage of cells infected with at least one copy of G. (g) The percentage of cells infected with at least one copy of Y and C. (h) The percentage of cells infected with at least one copy of G and C. (i) The percentage of cells infected with at least one copy of G and Y.

| experiment | $\beta_{\text{shaking}}$ | $\beta_{\text{static}}$ | $\gamma$ | $\beta_{\text{static}}/\gamma$ | $\beta_{\text{static}}(1-\varepsilon)/\gamma$ |
|---|---|---|---|---|---|
| A4/A1 | 2.56 | $2.56/f$ | 1.23 | 1.56 | 0.78 |
| A5/A2 | 2.57 | $2.57/f$ | 1.05 | 1.84 | 0.96 |
| A6/A3 | 2.35 | $2.35/f$ | 1.19 | 1.48 | 0.91 |
| B4/B1 | 1.97 | $1.97/f$ | 0.78 | 1.90 | 1.16 |
| B5/B2 | 2.19 | $2.19/f$ | 0.74 | 2.23 | 1.22 |
| B6/B3 | 2.15 | $2.15/f$ | 0.73 | 2.21 | 1.26 |
| C3/C1 | 1.83 | $1.83/f$ | 0.80 | 1.72 | 1.08 |
| C4/C2 | 1.56 | $1.56/f$ | 0.82 | 1.43 | 1.10 |

Table S5: Comparing the contribution of free virus versus synaptic transmission for best fit model parameters using the simultaneous fitting method (Table S2) for each pair of experiments. We have $\beta_{\text{static}} = \beta_{\text{shaking}}/f$, where $f$ is the shaking factor. Parameter descriptions can be found in Table S1.

### 4 Role of synaptic transmission in evolution

The rate of generation of cells infected with at least one copy of the double mutant strain is a measure of the evolutionary potential. Using the best fit experiment parameters, we can explore the role of the different transmission pathways (free virus versus synaptic cell-to-cell versus both pathways) on the spread of different strains using the mathematical models described in Section 1. In this way, we are able to simulate beyond the many constraints of the physical experiments.

In the main text, we describe evolutionary simulations that correspond to non-expanding populations of susceptible cells. Here we focus on populations that grow exponentially, which corresponds to the experimental conditions. Two types of simulations are explored: (1) We start with a small equal amount of cells singly infected with each of the single mutant strains. (2) We start with a small amount of cells singly infected with the wild type strain (which does not include either of the glowing mutations), and include the process of mutation. In both types of simulations we assume a large initial number of cells (populations of $10^9$ are used in the figures). We run the simulation until the number of infected cells is a given fraction of the beginning number of cells, such as 40%. In order to understand the contribution of the different transmission pathways, we investigate the number of double mutant recombinant infected cells under three different conditions: i) with both free virus and synaptic transmission. Here we use the same parameters as in the static experiments. ii) with only free virus transmission. Here we use the same parameters as in the static experiments, but set $\gamma = 0$. iii) only synaptic transmission. Here we use the same parameters as in the static experiments, but set $\beta = \epsilon = \xi = 0$.

In setup (1) above, in the absence of mutations, the double mutant has to be created by recombination between the single mutant strains. This can only occur in cells that are coinfected with both single mutant strains. The double mutant strain can then spread either by more recombination in single mutant coinfected cells, or through productive infection of cells infected with the double mutant. As shown in Figure S5(a), a combination of free and synaptic transmission results in the highest number and percentage of cells infected with the double mutant. This is because synaptic transmission promotes cotransmission of both single mutants from coinfected cells, and because $S = 3$, results in more opportunities for the double mutant to be created via recombination in coinfected cells. Free virus transmission, however, promotes the spread of the double mutant once it is generated because it can result in cells that are only infected with the double mutant (where it spreads better). Furthermore, including free virus transmission in addition to synaptic transmission helps the two single mutant strains to “meet” in the same cell to begin with, as with only one of the transmission pathways the number of infected cells increases, but the multiplicity of infection is low and the percentage of infected cells actually decreases (see Section 5). Therefore, a combination of both free virus and synaptic transmission is better than either one on their own, even though they

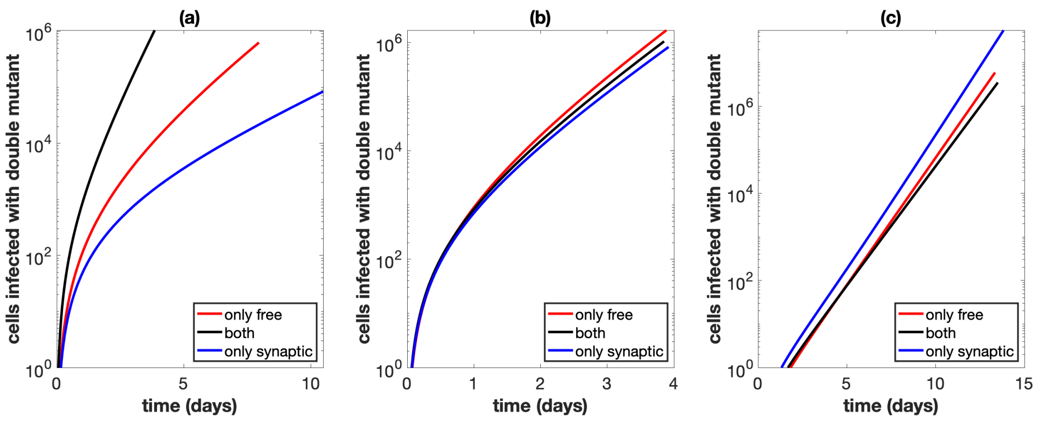


Figure S5: Total number of cells infected with at least one active copy of double mutant virus plotted against time. We assume we have $10^9$ initial cells, and run the simulation until the number of infected cells reaches 40% of this initial amount. Parameters are the best fit parameters from static experiment B2, which are listed in Table S2. The combination of both transmission pathways is represented by the black lines. For the only free virus transmission case (red lines), synaptic transmission is turned off. For the only synaptic transmission case (blue lines), free virus transmission and reinfection are turned off. (a) Parameters are exactly as in the experiment. (b) The overall growth rate is the same across the three cases. (c) The overall growth rate is the same across the three cases and the simulation starts with only a single infected cell, which is coinfected with both single mutant strains.

are both non-spatial in the context of the ODE system, and either one on their own results in a lower rate of infection which allows more time to get to the stopping threshold of $4 \times 10^8$ infected cells. For Figure S5(b) and (c), as in main text Figure 4(b) and (c), we assume that the overall growth rate is the same across the three cases. This leads to similar dynamics as discussed in the main text.

Next, we investigate the more realistic scenario (2), where we start with a small number of cells singly infected with the wild type, and include all forward and back mutations between the four strains, where for each infection event there is a $\mu$ chance to mutate at each of the two point locations. For instance, a wild type virus does not mutate with rate $(1-\mu)^2$, mutates into a single mutant strain with rate $\mu(1-\mu)$, and mutates into the double mutant strain with rate $\mu^2$, where $\mu = 3\times10^{-5}$ [4]. Here, we also assume that the self-reinfection terms apply only to the recombinant strain, since we start with the wild type and so most wild type is not generated by recombination in single mutant coinfected cells. In this setup, the single mutants have to be first created by mutation, and then the double mutant can be created either by mutation or recombination between cells coinfected with both single mutants (as direct mutation from the wild type to the double mutant is extremely rare). By turning recombination off, we find that most double mutants are created by mutation, which is promoted by synaptic infection, as for each infection event there is $S = 3$ chances for mutation to create the double mutant. As shown in Figure S6(b), when the infected cell population reaches $4\times10^8$ infected cells, synaptic transmission alone results in higher numbers of the double mutant compared to free virus transmission or a combination of both. The reason for this is two-fold: i) synaptic transmission is better to generate mutations because each transmission event results in $S = 3$ chances for mutation to occur, while free virus only results in 1, and ii) synaptic transmission only has the lowest rate of infection, which allows more time for mutants to be generated. For Figure S6(c) and (d), as in main text Figure 5(c) and (d), we assume that the overall growth rate is the same across the three cases, which again leads to similar dynamics as discussed in the main text.

Overall, it is clear that synaptic transmission plays a vital role in recombinant spread and infection evolution, because of multiple factors:

- Synaptic transmission can repeatedly cotransmit multiple mutant strains once they are to-

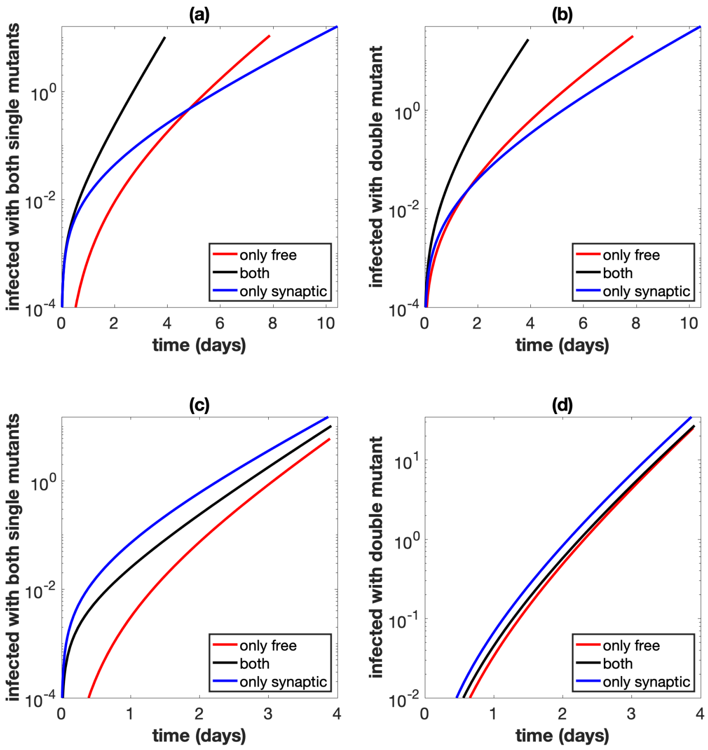


Figure S6: Evolutionary mutant dynamics. Same as Figure S5, but the simulations start with a small equal amount of cells singly infected with the wild type, and mutations are included. (a) Total number of cells infected with at least one copy of both single mutant strains. Parameters are exactly as in the experiment. (b) Total number of cells infected with at least one active copy of double mutant virus. Parameters are exactly as in the experiment. (c) Same as panel (a), but the overall growth rate is the same across the three cases. (d) Same as panel (b), but the overall growth rate is the same across the three cases.

gether in the same cell. This creates an environment where interactions between different strains can occur with cells (such as recombination).

- Synaptic transmission is beneficial to generate mutant strains, which could have evolutionary advantages. This is because more viral copies are transmitted per infection event, which gives rise to more chances for mutation.

- Similarly, transferring more viral copies per infection event gives rise to more chances for recombination, if the infecting cell is infected with distinct strains that allow for recombination.

It is also worthwhile to note that since cells have to form synapses between each other in order to initiate cell-to-cell transmission, there are likely spatial constraints to synaptic transmission (as we have found with free virus transmission through the experimental data fitting). However, all of the benefits of synaptic transmission for evolution listed above are still true and relevant in the context of spatial cell-to-cell transmission. Furthermore, the ability of synaptic transmission to repeatedly cotransmit multiple mutant strains once they are together in the same cell is amplified in the context of spatial transmission, as spatial clumps of cells all infected with mutants strains can occur [2].

Finally, Figure S7 shows an example of a shaking experiment (A6) and the corresponding static experiment (A3). In panel (e), we see that the static experiment (where both transmission pathways operate) results in higher levels of the recombinant G strain, supporting the idea that synaptic transmission plays an important role in recombinant evolution by promoting recombinant generation through repeated cotransmission of single-mutant viruses and also by increasing the overall rate of virus growth. However, in the experimental setup it is difficult to modulate the rate of infection and isolate cell-to-cell transmission (by turning off free virus transmission) and so we use the mathematical model parameterized by the experimental data in order to further study recombinant dynamics.

#### 5 Further theoretical considerations

Disregarding latent infection, we have that the basic reproductive ratio $R_0$ for the mathematical model is $\frac{\beta+\gamma}{a}$. If $R_0 > 1$ the number of infected cells will grow, and if $R_0 < 1$, then the number of infected cells will decline. However, to compare to the experiments, we are also concerned with the dynamics of the percentage of infected cells. We define the "reproductive ratio" for the percentage of infected cells, $R_{\%}$, as $R_{\%} = \frac{\beta+\gamma}{a+r}$. That is if $R_{\%} > 1$, the percentage of infected cells will increase, eventually reach 100%, and then all infected cells will eventually die. If $R_{\%} < 1$, the percentage of infected cells will decline and approach 0%.

Since the reproductive rate of uninfected cells is non-negative ($r \geq 0$), we have that $R_{\%} \leq R_0$. The infection dynamics can then be split into three cases:

- $R_{\%} \leq R_0 < 1$. Here, the infected cells will die out, and the number of uninfected cells will continue to grow toward infinity.

- $R_{\%} < 1 < R_0$. Here, the number of infected cells will increase, but the percentage of infected cells will decrease, as the number of uninfected cells will increase faster than the number of infected cells. So the number of uninfected and infected cells will approach infinity, and the percentage of infected cells will approach 0%.

- $1 < R_{\%} \leq R_0$. Here, the percentage of infected cells will increase, eventually reaching 100%, and then all cells will die.

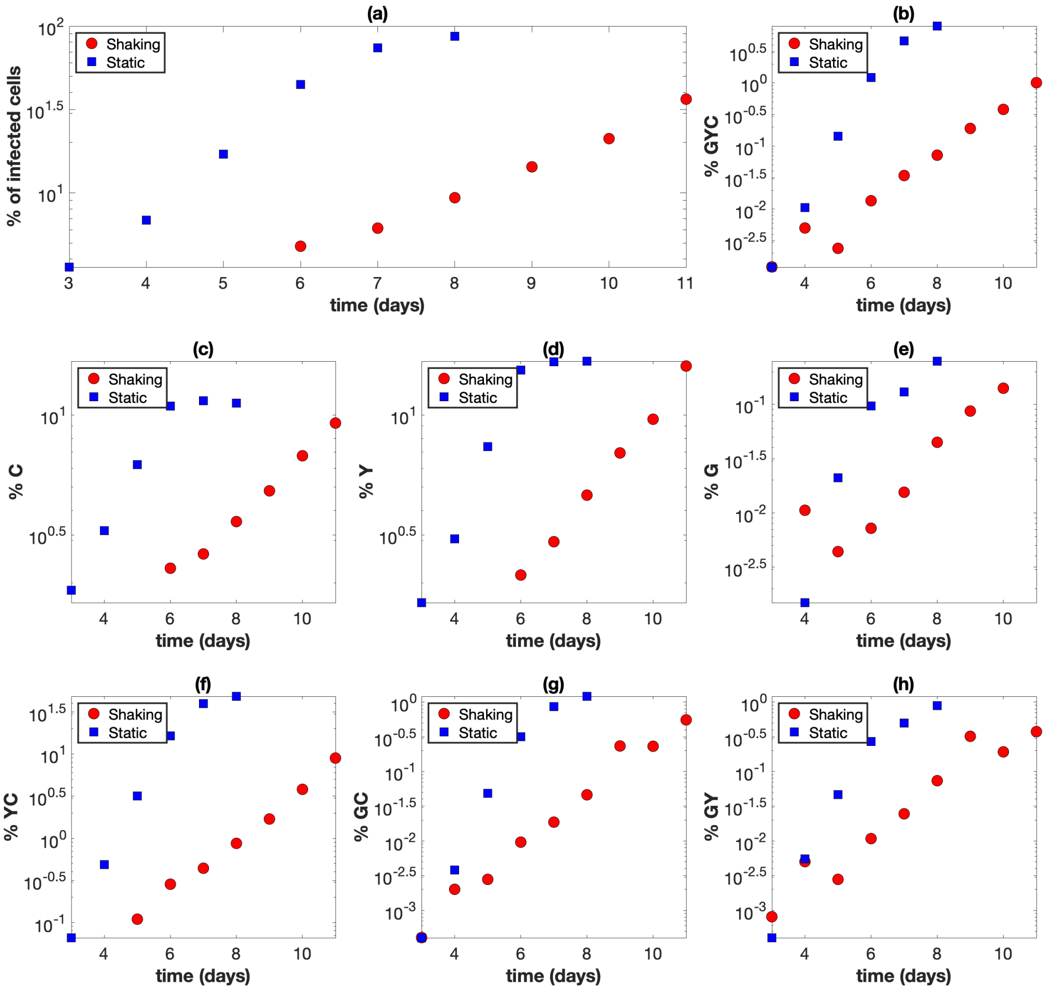


Figure S7: Shaking experiment A6 (red circles) and corresponding static experiment A3 (blue squares). The horizontal axis represents time (days). (a) The overall percentage of infected cells. (b) The percentage of cells infected with at least one copy of G, Y, and C. (c) The percentage of cells infected with at least one copy of C. (d) The percentage of cells infected with at least one copy of Y. (e) The percentage of cells infected with at least one copy of G. (f) The percentage of cells infected with at least one copy of Y and C. (g) The percentage of cells infected with at least one copy of G and C. (h) The percentage of cells infected with at least one copy of G and Y.

We can also modify the model in order introduce an infection steady state. This allows us to test model predictions with theoretically determined steady states from the differential equations. To do this, we modify the assumptions on the uninfected cells. Instead of assuming that uninfected cells reproduce with rate $r$, we assume that there is a constant influx of uninfected cells with rate $\lambda$, and that uninfected cells die with rate $d$. If we let $x$ denote the number of uninfected cells, then in the model we replace the $rx$ term with $\lambda - dx$.

Let $y$ denote the total number of infected cells. The model (grouping all infected cells together) is then

$$\dot{x} = \lambda - \frac{(\beta+\gamma)xy}{x+y} - dx \tag{23}$$

$$\dot{x} = \frac{(\beta+\gamma)xy}{x+y} - ay \tag{24}$$

The infection free steady state is given by

$$x = \frac{\lambda}{d}, \quad y = 0, \tag{25}$$

and the infection steady state is given by

$$x = \frac{\lambda}{\beta+\gamma-a+d}, \quad y = \frac{\lambda(\beta+\gamma-a)}{a(\beta+\gamma-a+d)}. \tag{26}$$

We then verify that our simulations for each of the different transmission strategies results in the correct steady state. An example of this can be seen in Figure S8.

### 6 Further justification of parameter estimates

We have performed further analysis to justify our parameter estimates and ensure the reporting of global minima. Because of the high dimension (number of parameters being fitted) it is not computationally feasible to calculate the error for a fine mesh of points within reasonable parameter ranges and pick the parameters that correspond to the lowest error. Therefore, we employed the following strategy.

We used multiple fitting algorithms and multiple random initial parameter guesses to estimate best fit parameters. We implemented the Levenberg-Marquardt Algorithm, which uses an approximate Gauss-Newton direction to approximate the minimum. Depending on the range of the initial parameter guesses, this method returns the same best fit parameters about 85% of the time, and either gets stuck at a local minimum or does not converge the other 15% of the time. For example, Figure S9 shows a plot of random initial parameter guesses for Levenberg-Marquardt Algorithm and where they converge to for two parameters $(\beta, \varepsilon)$ for experiment A6 (here we use the sequential fitting method because of the lower computational cost). The random initial guesses that converge are denoted in blue and the guesses that either do not converge or get stuck at a local minimum (with higher error) are denoted in green. The best fit parameters are denoted by the large red dot. If we zoom in on the best fit parameters (red dot), then a higher percentage of initial parameter guesses within that range will converge to it, and if we zoom out then a lower percentage of initial parameter guesses within that range will converge to the best fit parameters.

Further, we have also implemented the Trust-Region-Reflective Least Squares Algorithm, which is a robust global convergence algorithm based on trust regions (Figure S10). For this method, for $10^3$ random initial guesses within a reasonable range (i.e. $\beta \in [0, 10]$) the algorithm converged to

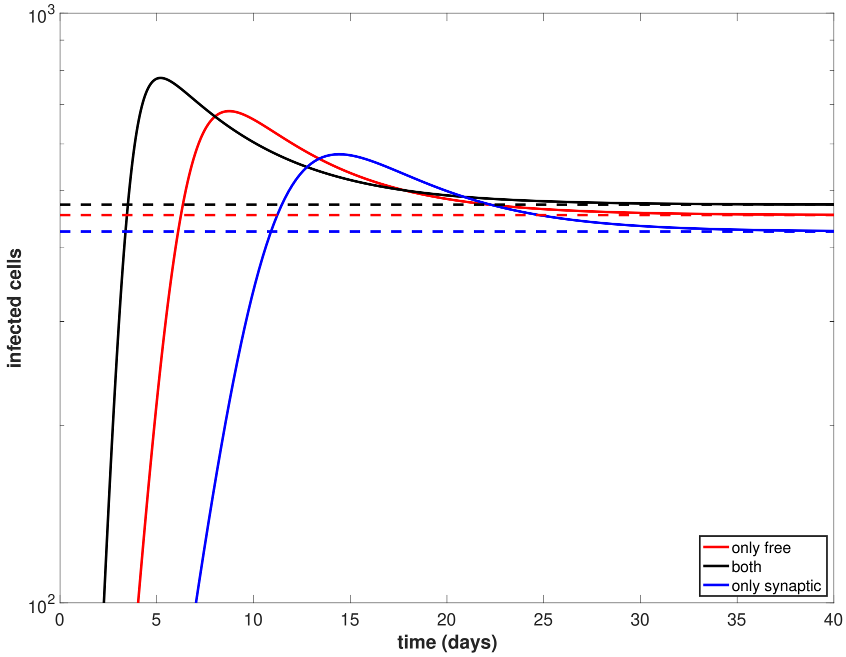


Figure S8: An example of convergence to the theoretical steady state when $rx$ is replaced by $\lambda - dx$ in the uninfeced cell equation. Parameters are as in Figures S5 and S6 (best fit parameters from static experiment B2, which are listed in Table S2), except we turn latent infection off and thus set $\varepsilon = 0$. Further, we arbitrarily set $\lambda = 100$ and $d = 0.1$. For each case, the solid line is the simulation of the model and the dashed line is the theoretical prediction of the steady state. The black lines represent both free virus and synaptic transmission, the red lines represent only free virus transmission, and the blue lines represent only synaptic transmission.

the same global minimum 100% of the time (again using the sequential fitting method because of computational cost). If we expand the range large enough (i.e. $\beta \in [0, 100]$), then it is possible for the algorithm to report different parameter estimates and get stuck at a local minimum with much higher error. This, along with reasonable confidence intervals and bands, helps to justify our parameter estimates.

Finally, we can visualize the error landscape with a two-dimension projection by fixing all but two parameters and generating a heatmap. Figures S11 and S12 show the log of the error for the simultaneous fitting of corresponding static experiment B2 and shaking experiment B5 and a range of values for specific parameter pairs $(\beta, \varepsilon)$ and $(\rho, \xi)$ while all other parameters are fixed. Here, we can calculate the error for a fine mesh of points and find the global minimum, which matches the results from the fitting algorithms. While this approach greatly reduces the dimension of the problem and is not a proof that global minima are obtained by the fitting algorithms, it provides further evidence for our parameter estimates.

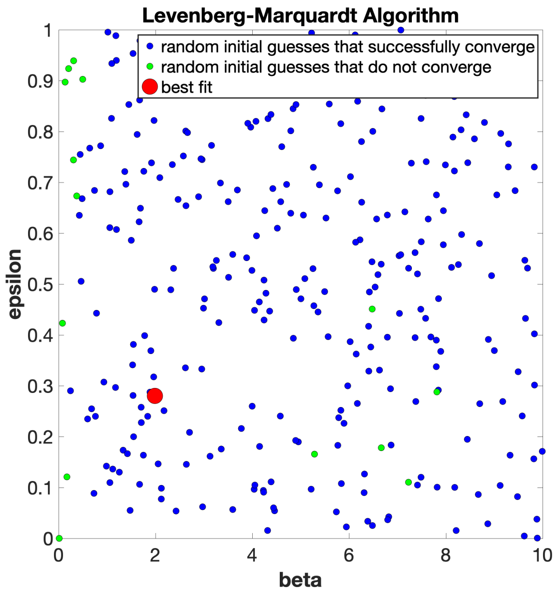


Figure S9: Plot of 300 random initial parameter guesses for the Levenberg-Marquardt Algorithm and where they converge to for two parameters $(\beta, \varepsilon)$ for experiment A6 (sequential fitting procedure, see Table S3). The random initial guesses that converge are denoted in blue. The random initial guesses that either do not converge or get stuck at a local minimum are denoted in green. The best fit parameters are denoted by the large red dot.

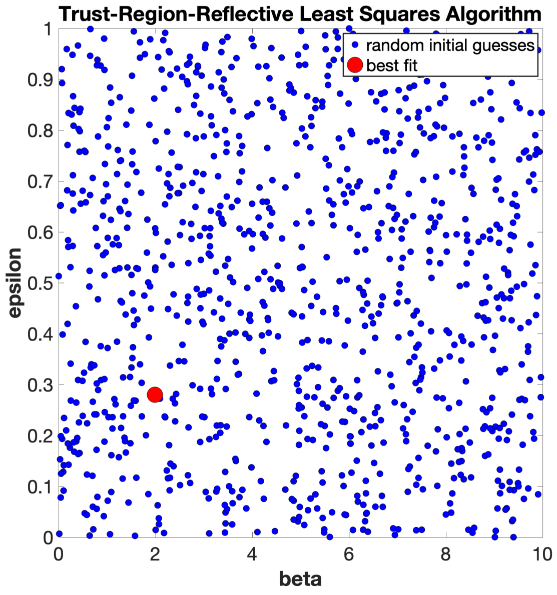


Figure S10: Plot of $10^3$ random initial parameter guesses for the Trust-Region-Reflective Least Squares Algorithm and where they converge to for two parameters $(\beta, \varepsilon)$ for experiment A6 (sequential fitting procedure, see Table S3). The random initial guesses that converge are denoted in blue. The best fit parameters are denoted by the large red dot. For the given range ($\beta \in [0, 10]$) all initial guesses converge to the best fit parameters, however it is possible to extend the range for $\beta$ such that this is no longer the case.

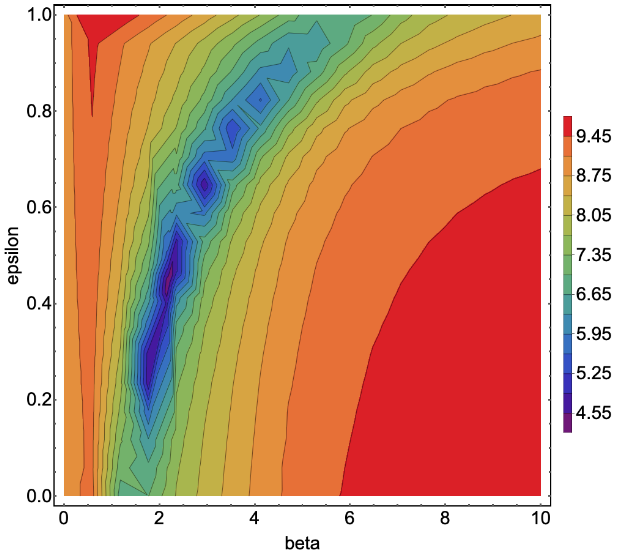


Figure S11: Heat plot of log error for the simultaneous fitting of corresponding static experiment B2 and shaking experiment B5 when all parameters other than $\beta$ and $\varepsilon$ are fixed. The horizontal axis represents $\beta$ (the rate of free virus transmission) and the vertical axis represents $\varepsilon$ (latent infection probability). The global minimum (which matches what is found by the fitting procedure, see Table S2) is at about (2.19,0.45).

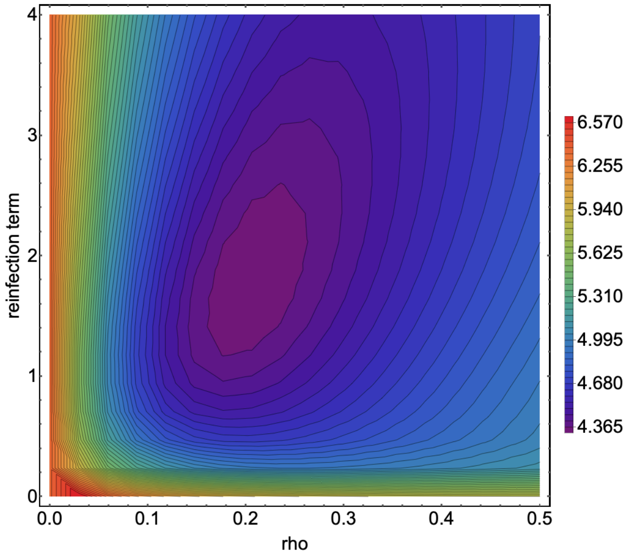


Figure S12: Heat plot of log error for the simultaneous fitting of corresponding static experiment B2 and shaking experiment B5 when all parameters other than $\rho$ and $\xi$ are fixed. The horizontal axis represents $\rho$ (probability of recombination) and the vertical axis represents $\xi$ (reinfection term). The global minimum (which matches what is found by the fitting procedure, see Table S2) is at about (0.20,1.76).

#### 7 Best fit parameters and plots

The fits (using the simultaneous fitting procedure) are given in Figures S13-S28. We present each experiment with pointwise (non-simultaneous) functional confidence bands (also obtained using the simultaneous fitting procedure). As the plots are on a log scale, negative lower bounds to the confidence interval are not shown. Each of the eight pairs of experiments is presented with the shaking experiment first, followed by the corresponding static experiment.

#### 8 Comparing the effect of viral transmission pathways on recombinant dynamics for all best fit parameter values

In this section, as in the main text, we examine the effect of the viral transmission pathways on recombinant dynamics. Therefore, we include figures that are similar to Figures 4 and 5 in the main text, for best fit parameters obtained for each of the eight pairs of experiments. Figures S29-S36 are comparable to Figure 4 in the main text, and Figure S37-S44 are comparable to Figure 5 in the main text. The parameters are obtained by the simultaneous fitting method.

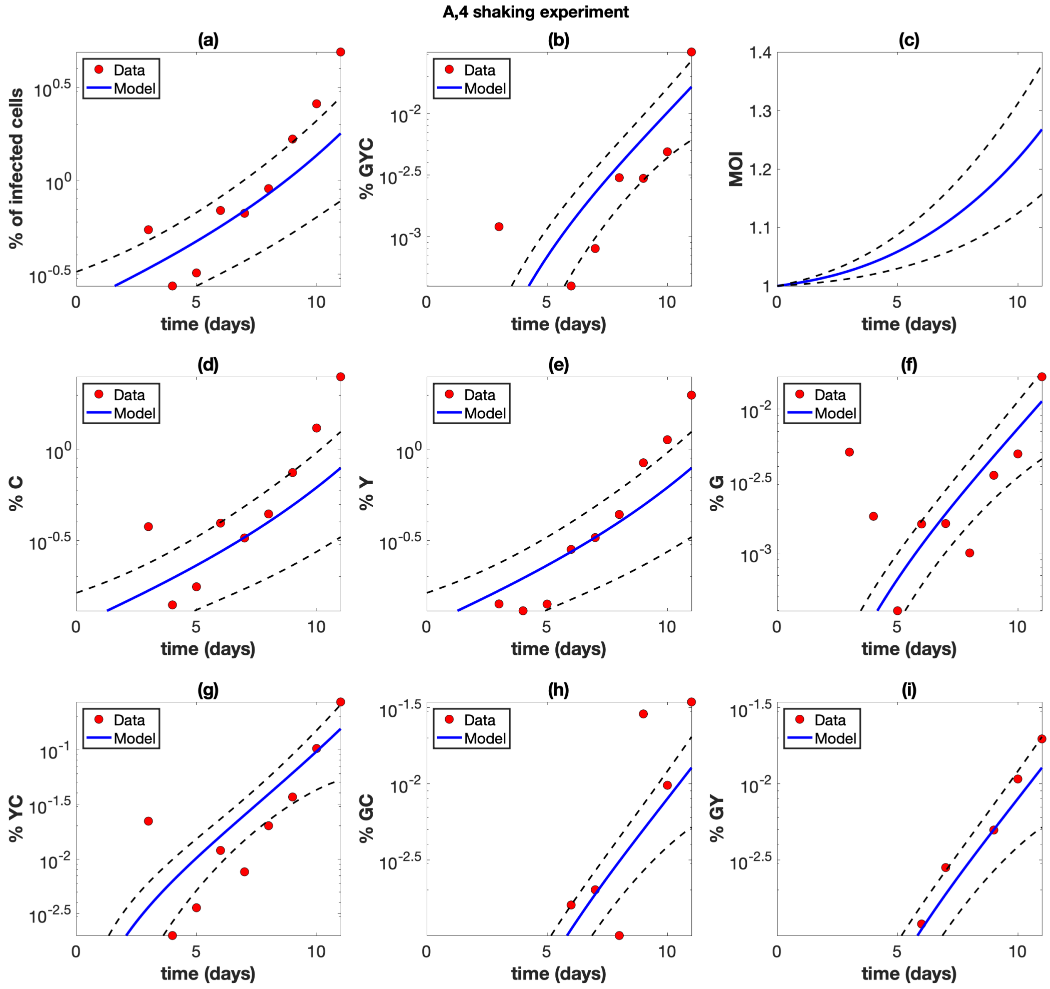


Figure S13: Shaking experiment A4. The experimental data (red circles) are presented with best fit curves from the model (blue lines). The mathematical model is described in Section 1 and the fitting procedure is described in Section 2. Best fit parameters are included in Table S2. The horizontal axis represents time (days). (a) The overall percentage of infected cells. (b) The percentage of cells infected with at least one copy of G, Y, and C. (c) The average multiplicity of infection (MOI) over all infected cells. (d) The percentage of cells infected with at least one copy of C. (e) The percentage of cells infected with at least one copy of Y. (f) The percentage of cells infected with at least one copy of G. (g) The percentage of cells infected with at least one copy of Y and C. (h) The percentage of cells infected with at least one copy of G and C. (i) The percentage of cells infected with at least one copy of G and Y. The dashed black lines represent the pointwise 95% prediction confidence bounds.

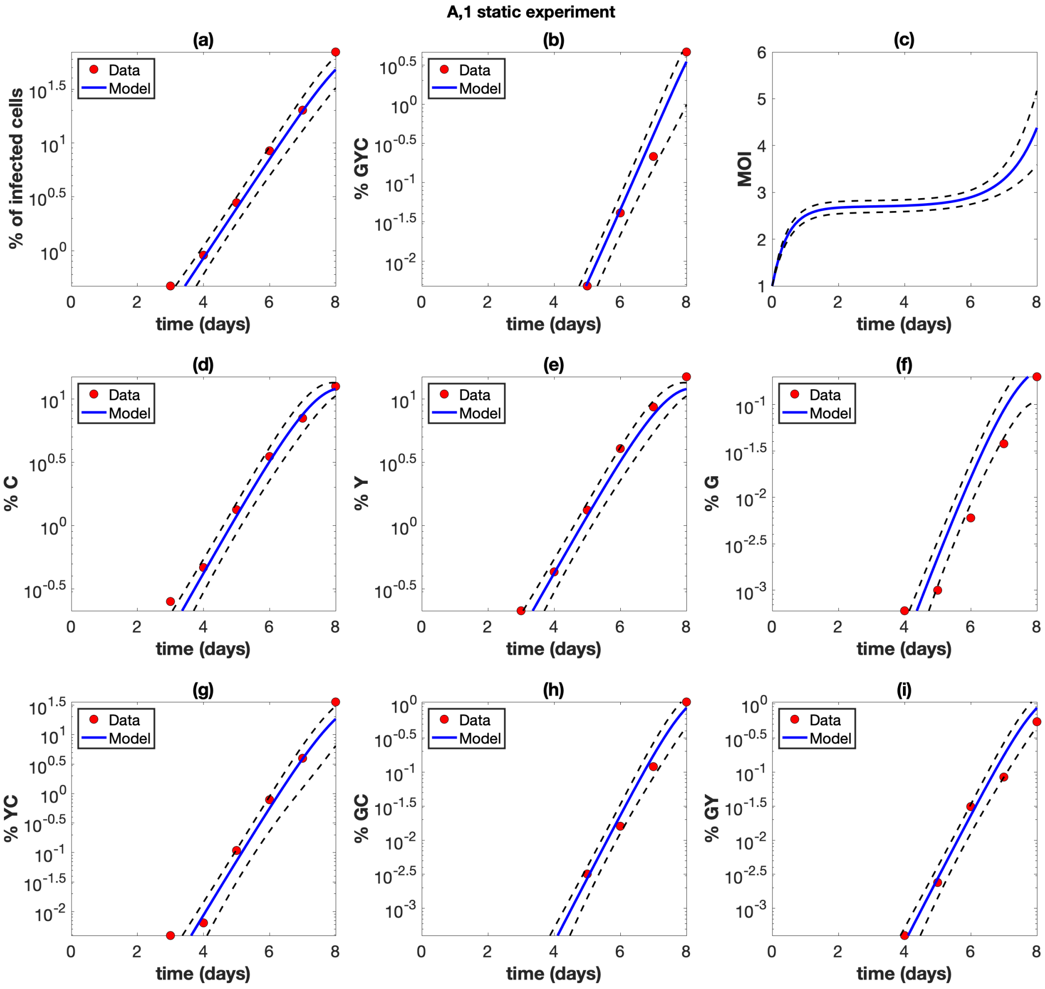


Figure S14: Static experiment A1. The experimental data (red circles) are presented with best fit curves from the model (blue lines). The mathematical model is described in Section 1 and the fitting procedure is described in Section 2. Best fit parameters are included in Table S2. The horizontal axis represents time (days). (a) The overall percentage of infected cells. (b) The percentage of cells infected with at least one copy of G, Y, and C. (c) The average multiplicity of infection (MOI) over all infected cells. (d) The percentage of cells infected with at least one copy of C. (e) The percentage of cells infected with at least one copy of Y. (f) The percentage of cells infected with at least one copy of G. (g) The percentage of cells infected with at least one copy of Y and C. (h) The percentage of cells infected with at least one copy of G and C. (i) The percentage of cells infected with at least one copy of G and Y. The dashed black lines represent the pointwise 95% prediction confidence bounds.

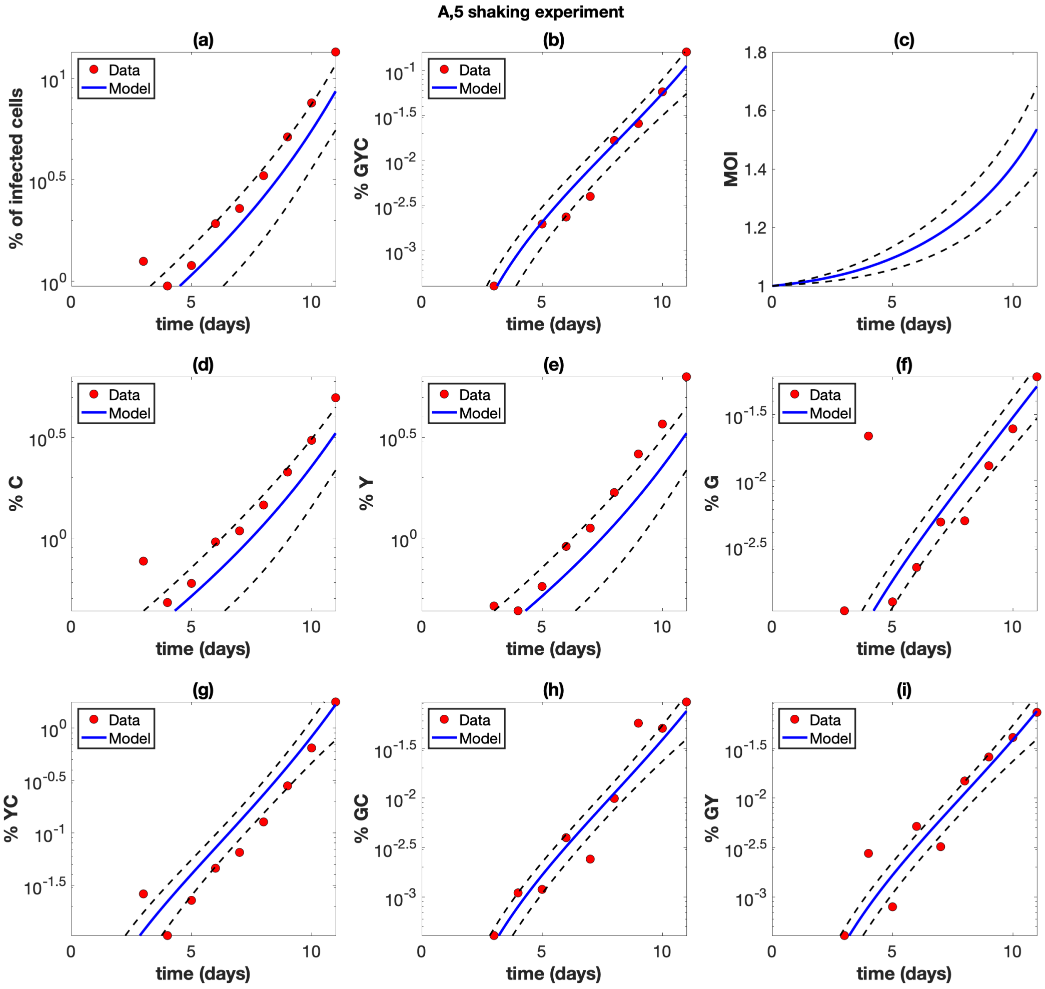


Figure S15: Shaking experiment A5. The experimental data (red circles) are presented with best fit curves from the model (blue lines). The mathematical model is described in Section 1 and the fitting procedure is described in Section 2. Best fit parameters are included in Table S2. The horizontal axis represents time (days). (a) The overall percentage of infected cells. (b) The percentage of cells infected with at least one copy of G, Y, and C. (c) The average multiplicity of infection (MOI) over all infected cells. (d) The percentage of cells infected with at least one copy of C. (e) The percentage of cells infected with at least one copy of Y. (f) The percentage of cells infected with at least one copy of G. (g) The percentage of cells infected with at least one copy of Y and C. (h) The percentage of cells infected with at least one copy of G and C. (i) The percentage of cells infected with at least one copy of G and Y. The dashed black lines represent the pointwise 95% prediction confidence bounds.

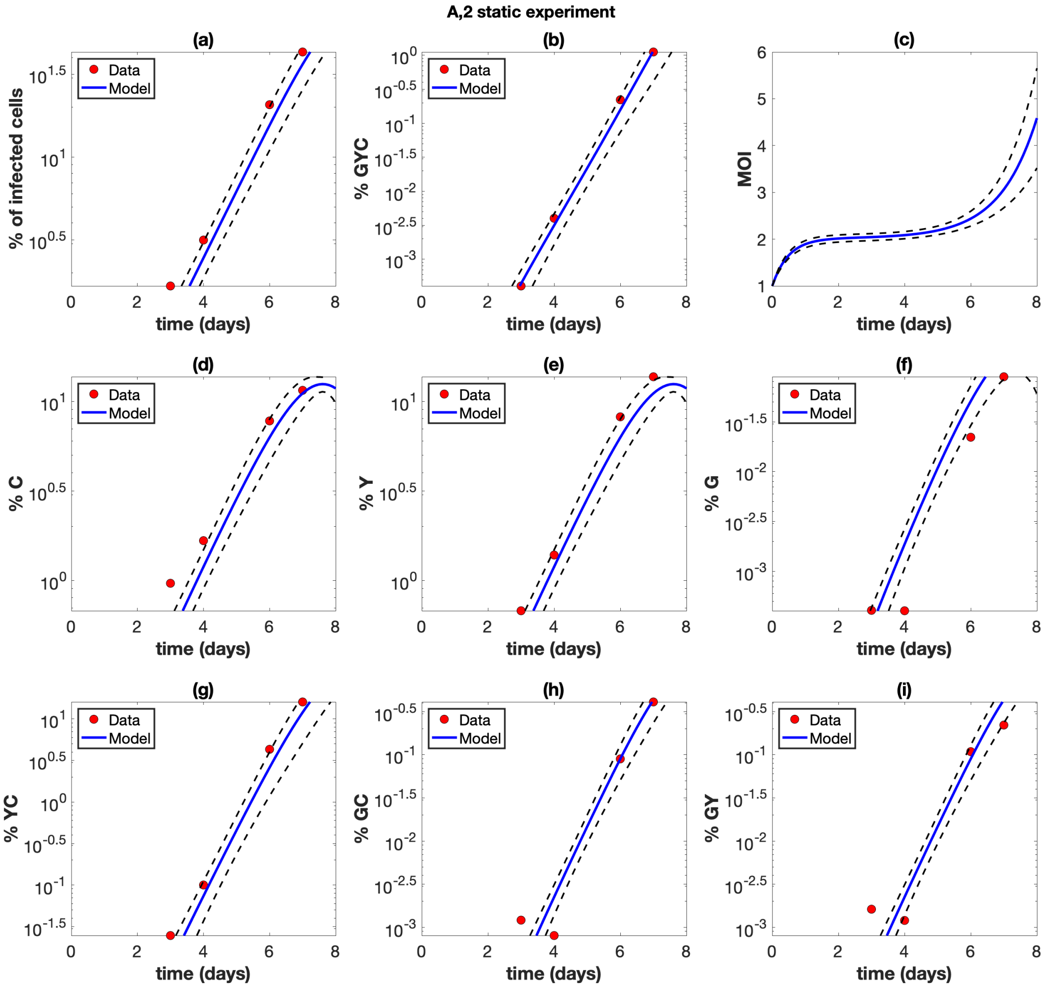


Figure S16: Static experiment A2. The experimental data (red circles) are presented with best fit curves from the model (blue lines). The mathematical model is described in Section 1 and the fitting procedure is described in Section 2. Best fit parameters are included in Table S2. The horizontal axis represents time (days). (a) The overall percentage of infected cells. (b) The percentage of cells infected with at least one copy of G, Y, and C. (c) The average multiplicity of infection (MOI) over all infected cells. (d) The percentage of cells infected with at least one copy of C. (e) The percentage of cells infected with at least one copy of Y. (f) The percentage of cells infected with at least one copy of G. (g) The percentage of cells infected with at least one copy of Y and C. (h) The percentage of cells infected with at least one copy of G and C. (i) The percentage of cells infected with at least one copy of G and Y. The dashed black lines represent the pointwise 95% prediction confidence bounds.

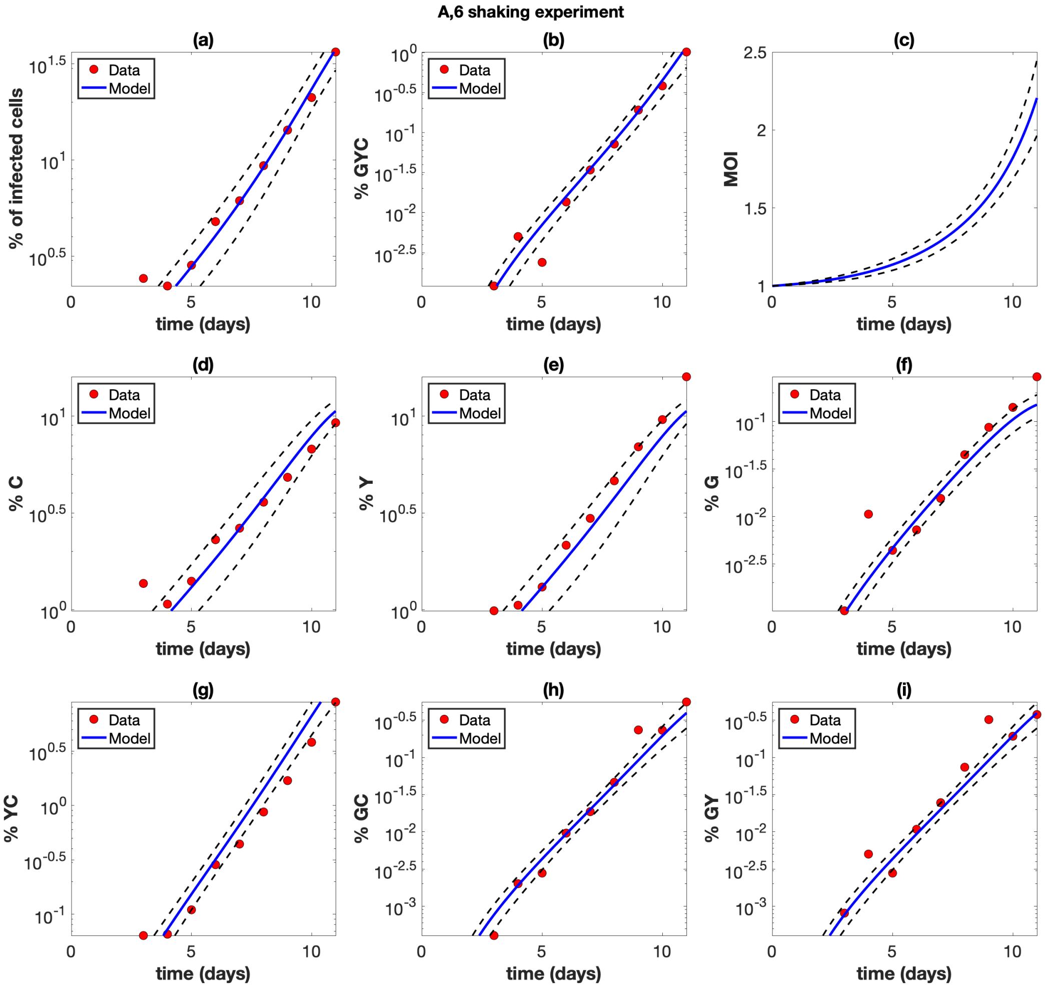


Figure S17: Shaking experiment A6. The experimental data (red circles) are presented with best fit curves from the model (blue lines). The mathematical model is described in Section 1 and the fitting procedure is described in Section 2. Best fit parameters are included in Table S2. The horizontal axis represents time (days). (a) The overall percentage of infected cells. (b) The percentage of cells infected with at least one copy of G, Y, and C. (c) The average multiplicity of infection (MOI) over all infected cells. (d) The percentage of cells infected with at least one copy of C. (e) The percentage of cells infected with at least one copy of Y. (f) The percentage of cells infected with at least one copy of G. (g) The percentage of cells infected with at least one copy of Y and C. (h) The percentage of cells infected with at least one copy of G and C. (i) The percentage of cells infected with at least one copy of G and Y. The dashed black lines represent the pointwise 95% prediction confidence bounds.

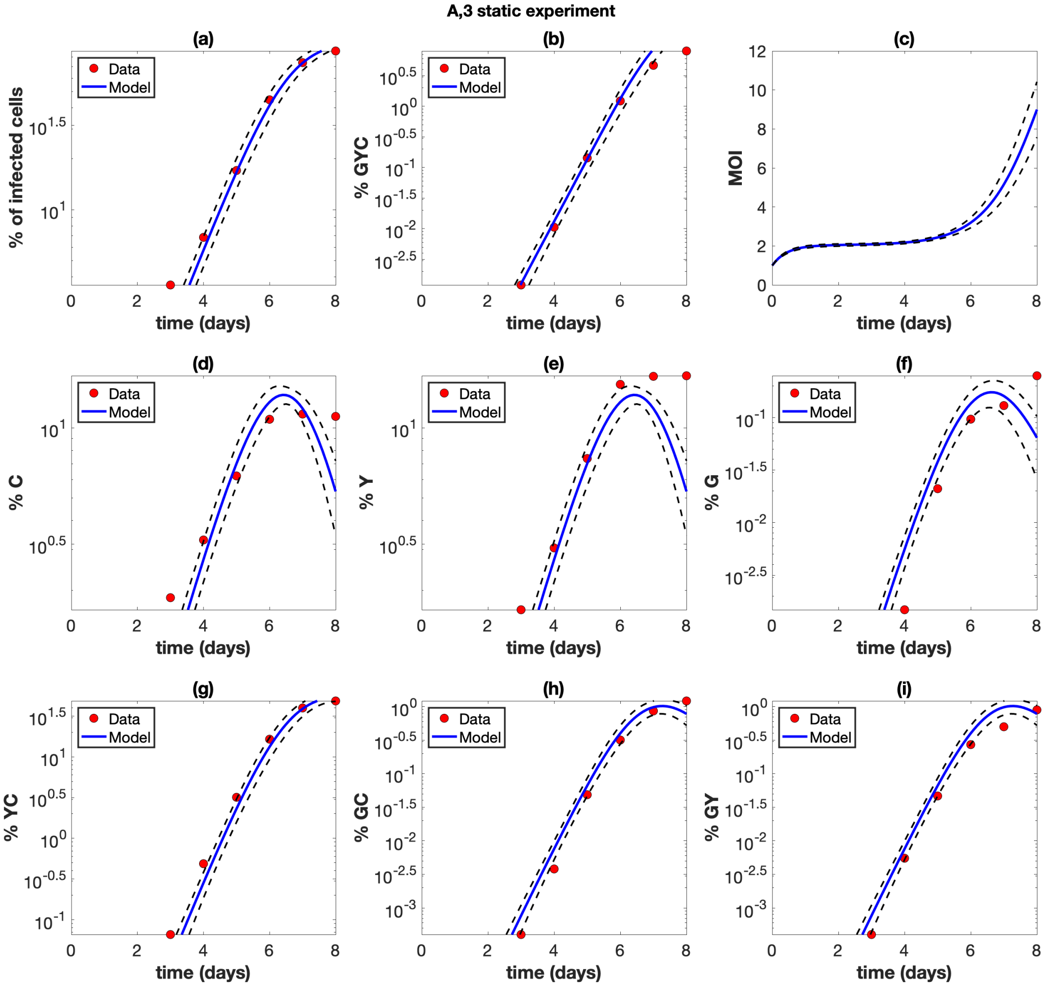


Figure S18: Static experiment A3. The experimental data (red circles) are presented with best fit curves from the model (blue lines). The mathematical model is described in Section 1 and the fitting procedure is described in Section 2. Best fit parameters are included in Table S2. The horizontal axis represents time (days). (a) The overall percentage of infected cells. (b) The percentage of cells infected with at least one copy of G, Y, and C. (c) The average multiplicity of infection (MOI) over all infected cells. (d) The percentage of cells infected with at least one copy of C. (e) The percentage of cells infected with at least one copy of Y. (f) The percentage of cells infected with at least one copy of G. (g) The percentage of cells infected with at least one copy of Y and C. (h) The percentage of cells infected with at least one copy of G and C. (i) The percentage of cells infected with at least one copy of G and Y. The dashed black lines represent the pointwise 95% prediction confidence bounds.

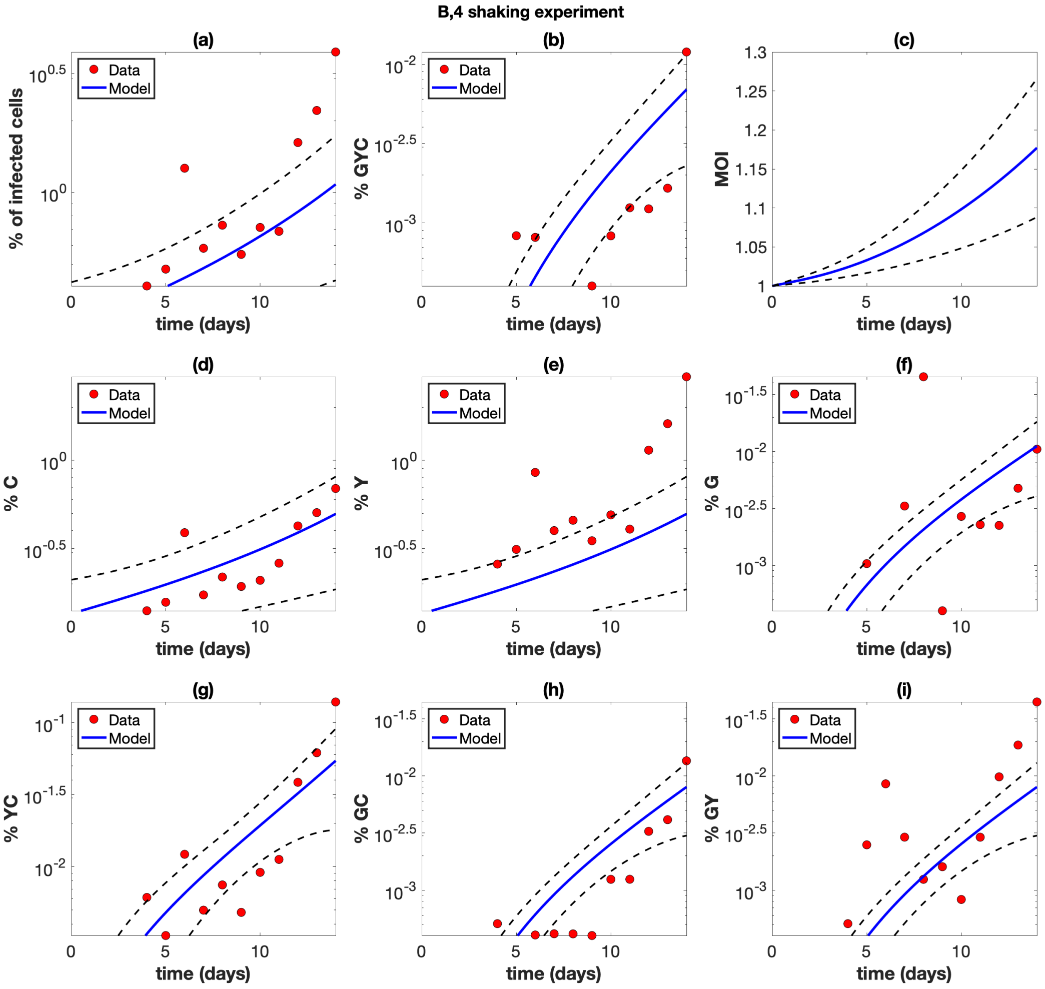


Figure S19: Shaking experiment B4. The experimental data (red circles) are presented with best fit curves from the model (blue lines). The mathematical model is described in Section 1 and the fitting procedure is described in Section 2. Best fit parameters are included in Table S2. The horizontal axis represents time (days). (a) The overall percentage of infected cells. (b) The percentage of cells infected with at least one copy of G, Y, and C. (c) The average multiplicity of infection (MOI) over all infected cells. (d) The percentage of cells infected with at least one copy of C. (e) The percentage of cells infected with at least one copy of Y. (f) The percentage of cells infected with at least one copy of G. (g) The percentage of cells infected with at least one copy of Y and C. (h) The percentage of cells infected with at least one copy of G and C. (i) The percentage of cells infected with at least one copy of G and Y. The dashed black lines represent the pointwise 95% prediction confidence bounds.

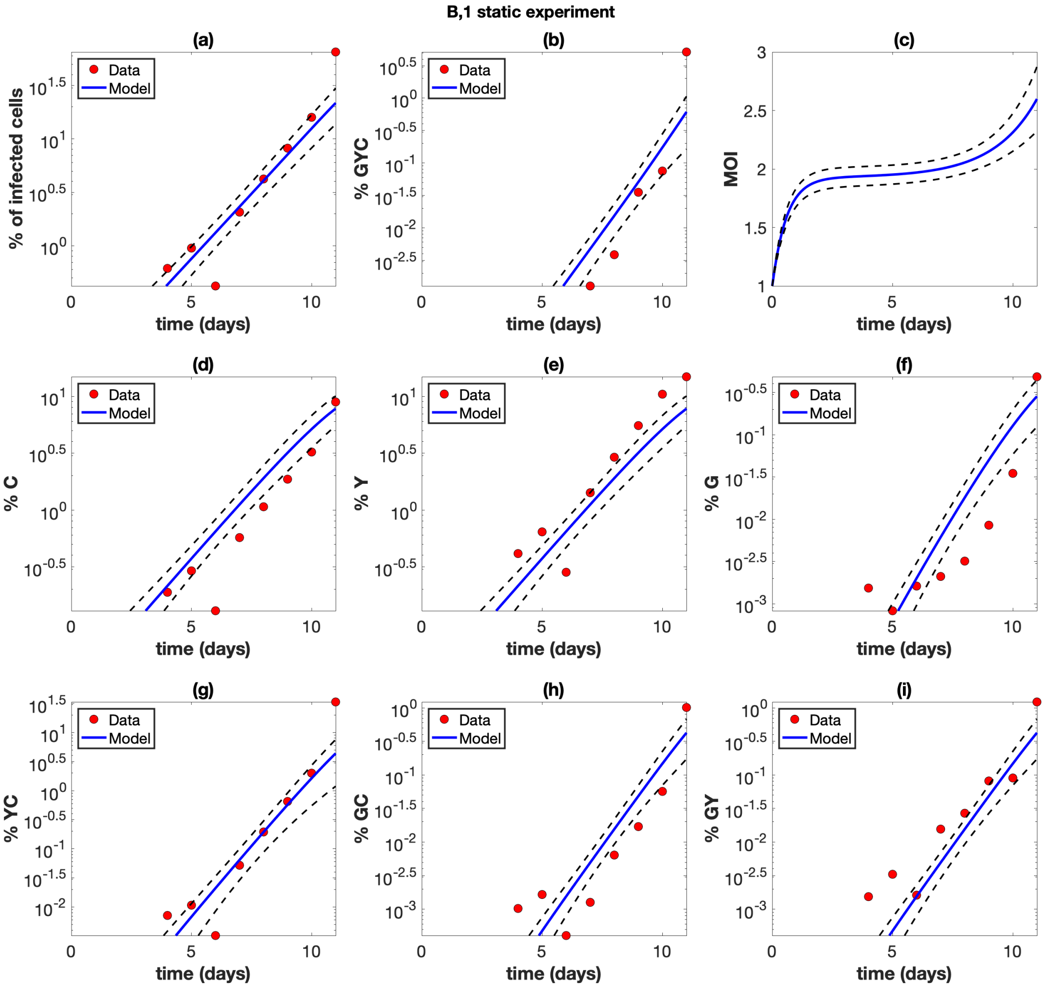


Figure S20: Static experiment B1. The experimental data (red circles) are presented with best fit curves from the model (blue lines). The mathematical model is described in Section 1 and the fitting procedure is described in Section 2. Best fit parameters are included in Table S2. The horizontal axis represents time (days). (a) The overall percentage of infected cells. (b) The percentage of cells infected with at least one copy of G, Y, and C. (c) The average multiplicity of infection (MOI) over all infected cells. (d) The percentage of cells infected with at least one copy of C. (e) The percentage of cells infected with at least one copy of Y. (f) The percentage of cells infected with at least one copy of G. (g) The percentage of cells infected with at least one copy of Y and C. (h) The percentage of cells infected with at least one copy of G and C. (i) The percentage of cells infected with at least one copy of G and Y. The dashed black lines represent the pointwise 95% prediction confidence bounds.

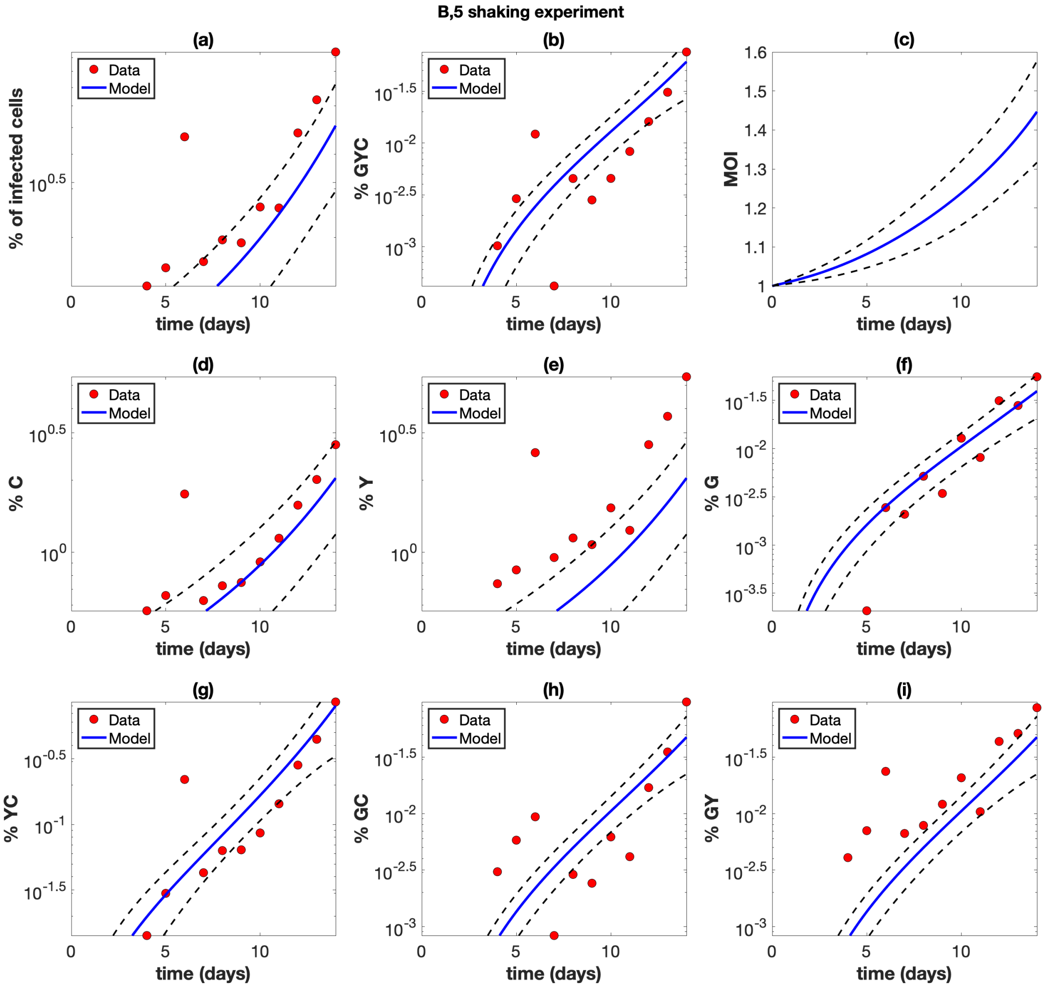


Figure S21: Shaking experiment B5. The experimental data (red circles) are presented with best fit curves from the model (blue lines). The mathematical model is described in Section 1 and the fitting procedure is described in Section 2. Best fit parameters are included in Table S2. The horizontal axis represents time (days). (a) The overall percentage of infected cells. (b) The percentage of cells infected with at least one copy of G, Y, and C. (c) The average multiplicity of infection (MOI) over all infected cells. (d) The percentage of cells infected with at least one copy of C. (e) The percentage of cells infected with at least one copy of Y. (f) The percentage of cells infected with at least one copy of G. (g) The percentage of cells infected with at least one copy of Y and C. (h) The percentage of cells infected with at least one copy of G and C. (i) The percentage of cells infected with at least one copy of G and Y. The dashed black lines represent the pointwise 95% prediction confidence bounds.

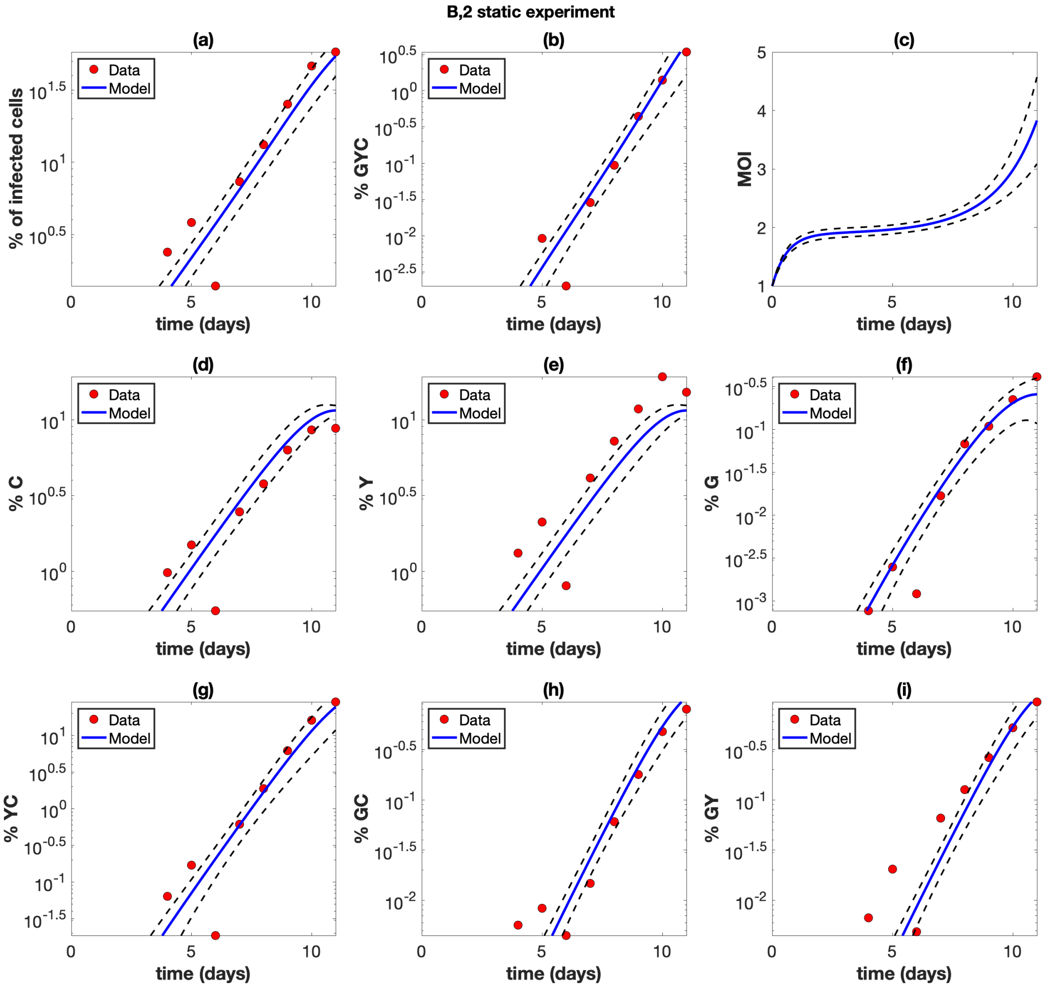


Figure S22: Static experiment B2. The experimental data (red circles) are presented with best fit curves from the model (blue lines). The mathematical model is described in Section 1 and the fitting procedure is described in Section 2. Best fit parameters are included in Table S2. The horizontal axis represents time (days). (a) The overall percentage of infected cells. (b) The percentage of cells infected with at least one copy of G, Y, and C. (c) The average multiplicity of infection (MOI) over all infected cells. (d) The percentage of cells infected with at least one copy of C. (e) The percentage of cells infected with at least one copy of Y. (f) The percentage of cells infected with at least one copy of G. (g) The percentage of cells infected with at least one copy of Y and C. (h) The percentage of cells infected with at least one copy of G and C. (i) The percentage of cells infected with at least one copy of G and Y. The dashed black lines represent the pointwise 95% prediction confidence bounds.

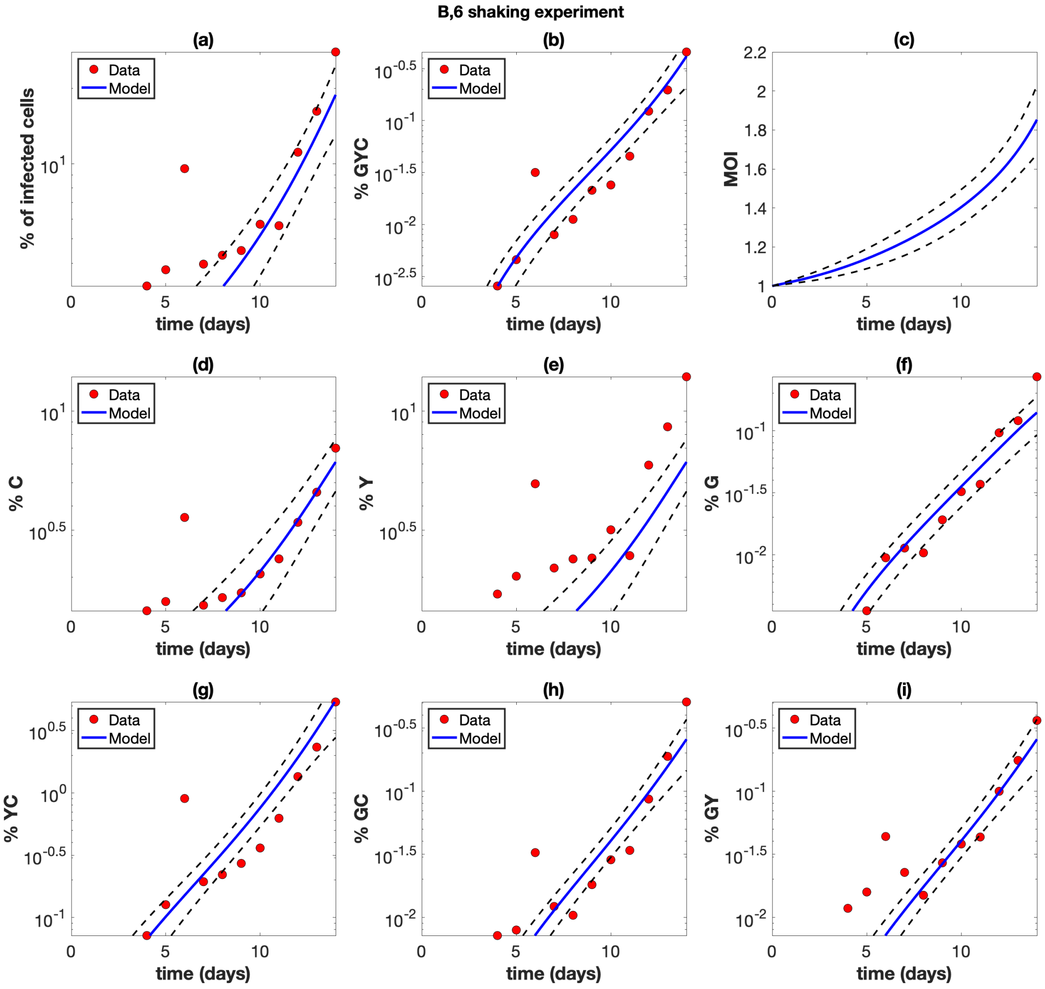


Figure S23: Shaking experiment B6. The experimental data (red circles) are presented with best fit curves from the model (blue lines). The mathematical model is described in Section 1 and the fitting procedure is described in Section 2. Best fit parameters are included in Table S2. The horizontal axis represents time (days). (a) The overall percentage of infected cells. (b) The percentage of cells infected with at least one copy of G, Y, and C. (c) The average multiplicity of infection (MOI) over all infected cells. (d) The percentage of cells infected with at least one copy of C. (e) The percentage of cells infected with at least one copy of Y. (f) The percentage of cells infected with at least one copy of G. (g) The percentage of cells infected with at least one copy of Y and C. (h) The percentage of cells infected with at least one copy of G and C. (i) The percentage of cells infected with at least one copy of G and Y. The dashed black lines represent the pointwise 95% prediction confidence bounds.

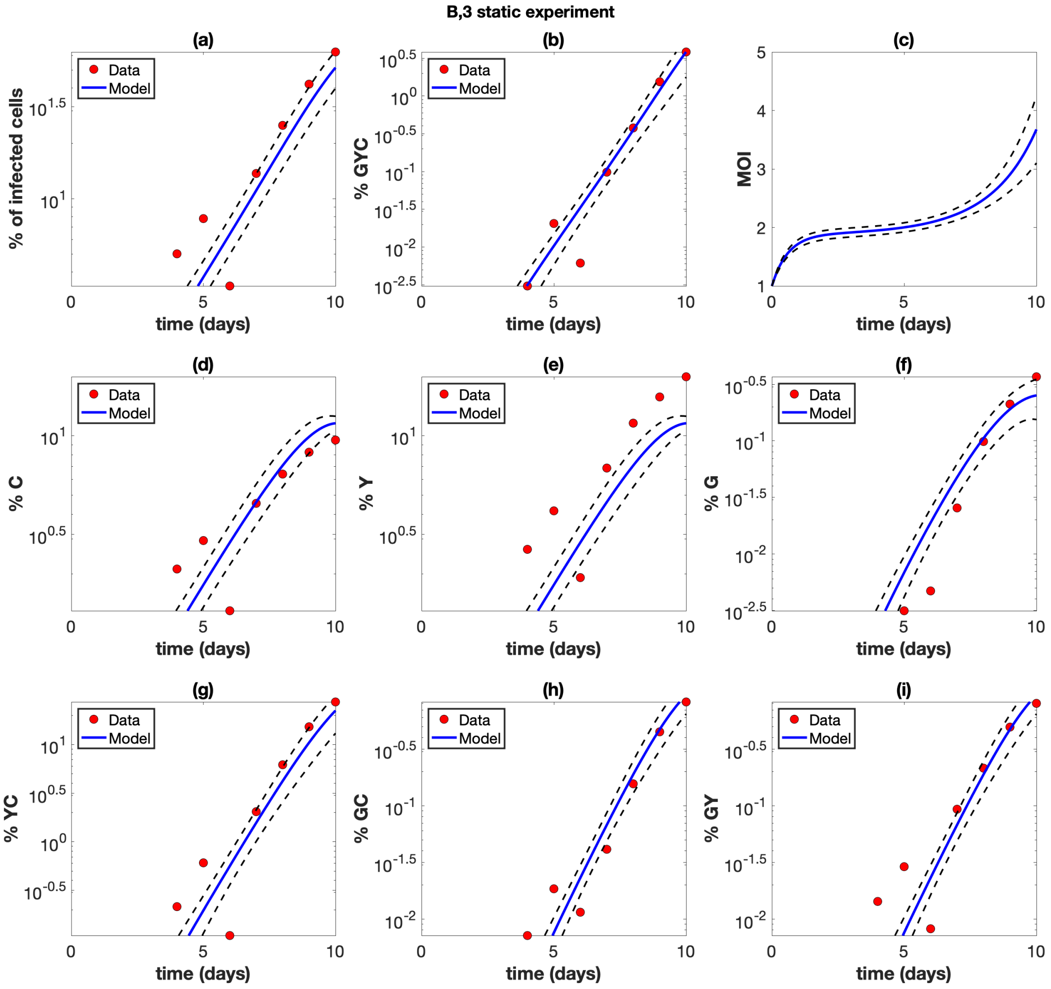


Figure S24: Static experiment B3. The experimental data (red circles) are presented with best fit curves from the model (blue lines). The mathematical model is described in Section 1 and the fitting procedure is described in Section 2. Best fit parameters are included in Table S2. The horizontal axis represents time (days). (a) The overall percentage of infected cells. (b) The percentage of cells infected with at least one copy of G, Y, and C. (c) The average multiplicity of infection (MOI) over all infected cells. (d) The percentage of cells infected with at least one copy of C. (e) The percentage of cells infected with at least one copy of Y. (f) The percentage of cells infected with at least one copy of G. (g) The percentage of cells infected with at least one copy of Y and C. (h) The percentage of cells infected with at least one copy of G and C. (i) The percentage of cells infected with at least one copy of G and Y. The dashed black lines represent the pointwise 95% prediction confidence bounds.

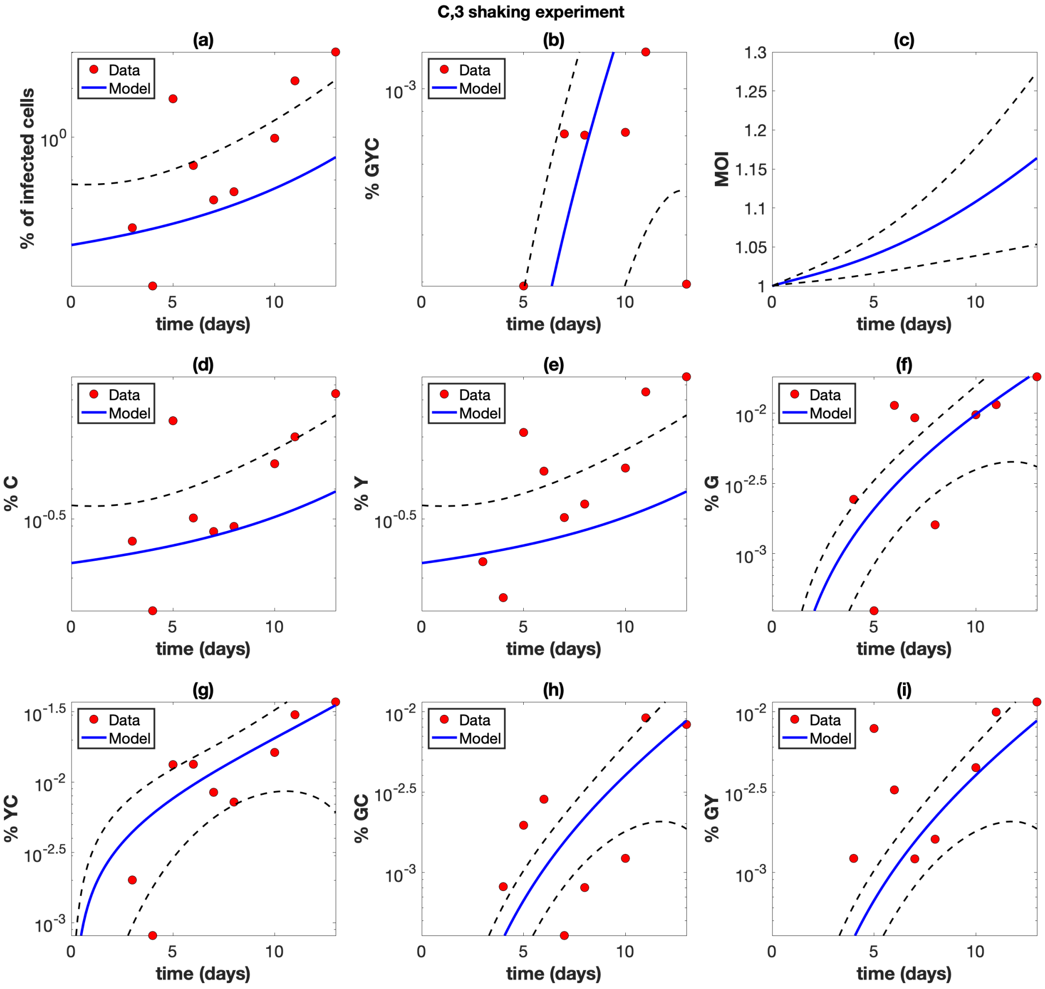


Figure S25: Shaking experiment C3. The experimental data (red circles) are presented with best fit curves from the model (blue lines). The mathematical model is described in Section 1 and the fitting procedure is described in Section 2. Best fit parameters are included in Table S2. The horizontal axis represents time (days). (a) The overall percentage of infected cells. (b) The percentage of cells infected with at least one copy of G, Y, and C. (c) The average multiplicity of infection (MOI) over all infected cells. (d) The percentage of cells infected with at least one copy of C. (e) The percentage of cells infected with at least one copy of Y. (f) The percentage of cells infected with at least one copy of G. (g) The percentage of cells infected with at least one copy of Y and C. (h) The percentage of cells infected with at least one copy of G and C. (i) The percentage of cells infected with at least one copy of G and Y. The dashed black lines represent the pointwise 95% prediction confidence bounds.

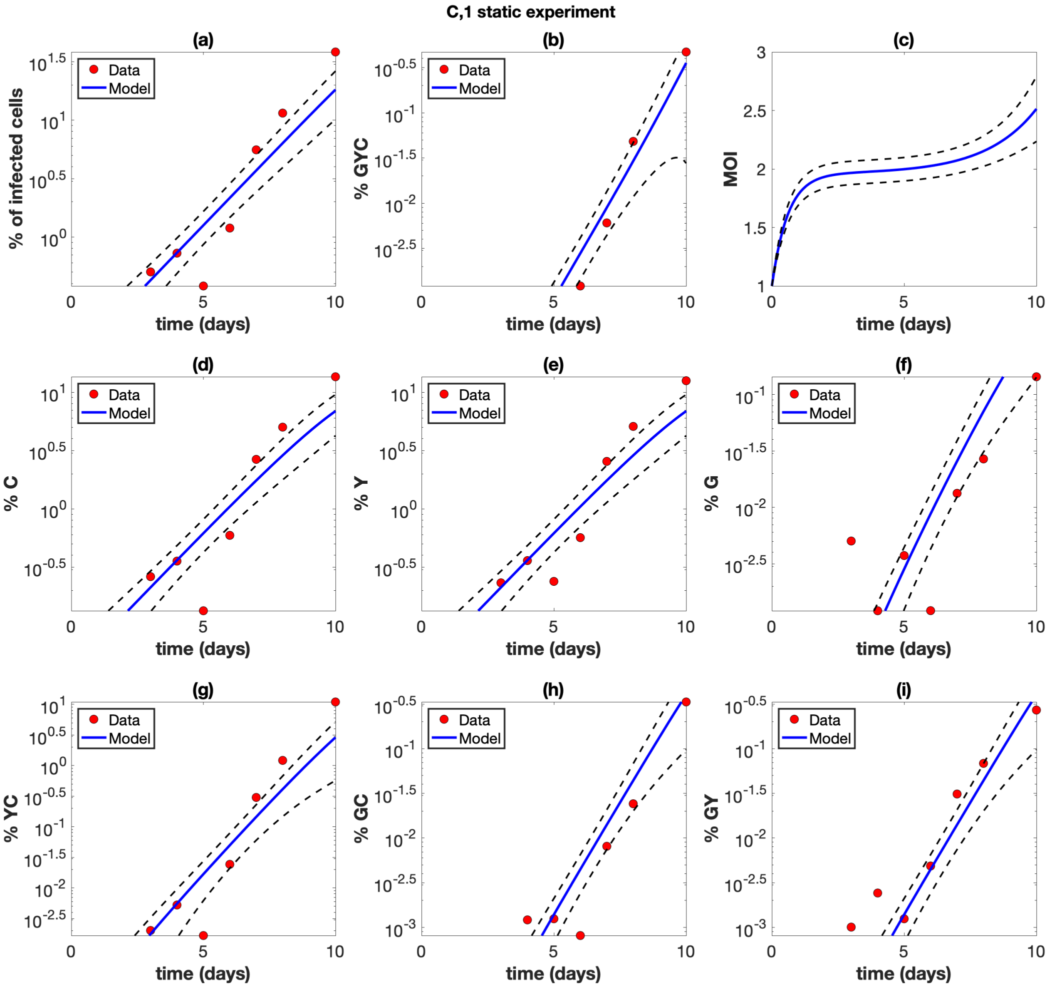


Figure S26: Static experiment C1. The experimental data (red circles) are presented with best fit curves from the model (blue lines). The mathematical model is described in Section 1 and the fitting procedure is described in Section 2. Best fit parameters are included in Table S2. The horizontal axis represents time (days). (a) The overall percentage of infected cells. (b) The percentage of cells infected with at least one copy of G, Y, and C. (c) The average multiplicity of infection (MOI) over all infected cells. (d) The percentage of cells infected with at least one copy of C. (e) The percentage of cells infected with at least one copy of Y. (f) The percentage of cells infected with at least one copy of G. (g) The percentage of cells infected with at least one copy of Y and C. (h) The percentage of cells infected with at least one copy of G and C. (i) The percentage of cells infected with at least one copy of G and Y. The dashed black lines represent the pointwise 95% prediction confidence bounds.

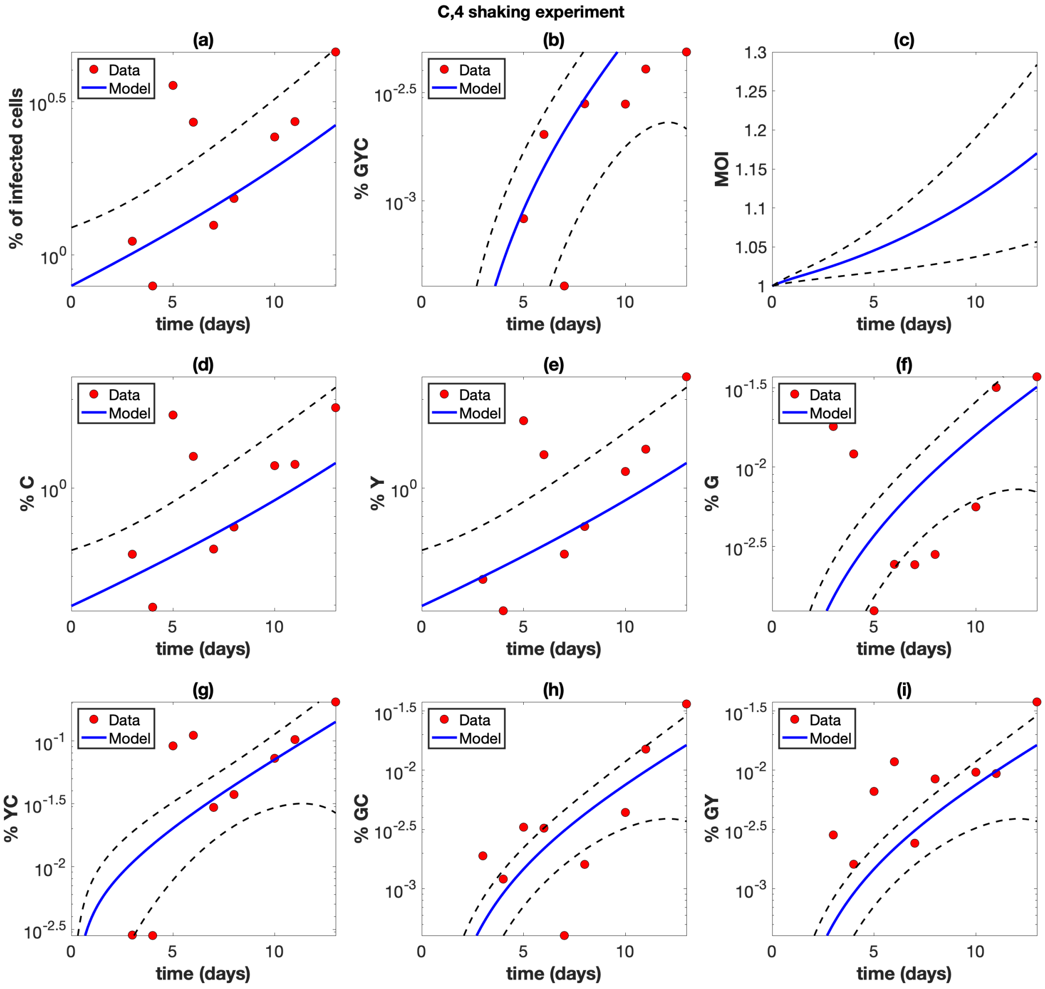


Figure S27: Shaking experiment C4. The experimental data (red circles) are presented with best fit curves from the model (blue lines). The mathematical model is described in Section 1 and the fitting procedure is described in Section 2. Best fit parameters are included in Table S2. The horizontal axis represents time (days). (a) The overall percentage of infected cells. (b) The percentage of cells infected with at least one copy of G, Y, and C. (c) The average multiplicity of infection (MOI) over all infected cells. (d) The percentage of cells infected with at least one copy of C. (e) The percentage of cells infected with at least one copy of Y. (f) The percentage of cells infected with at least one copy of G. (g) The percentage of cells infected with at least one copy of Y and C. (h) The percentage of cells infected with at least one copy of G and C. (i) The percentage of cells infected with at least one copy of G and Y. The dashed black lines represent the pointwise 95% prediction confidence bounds.

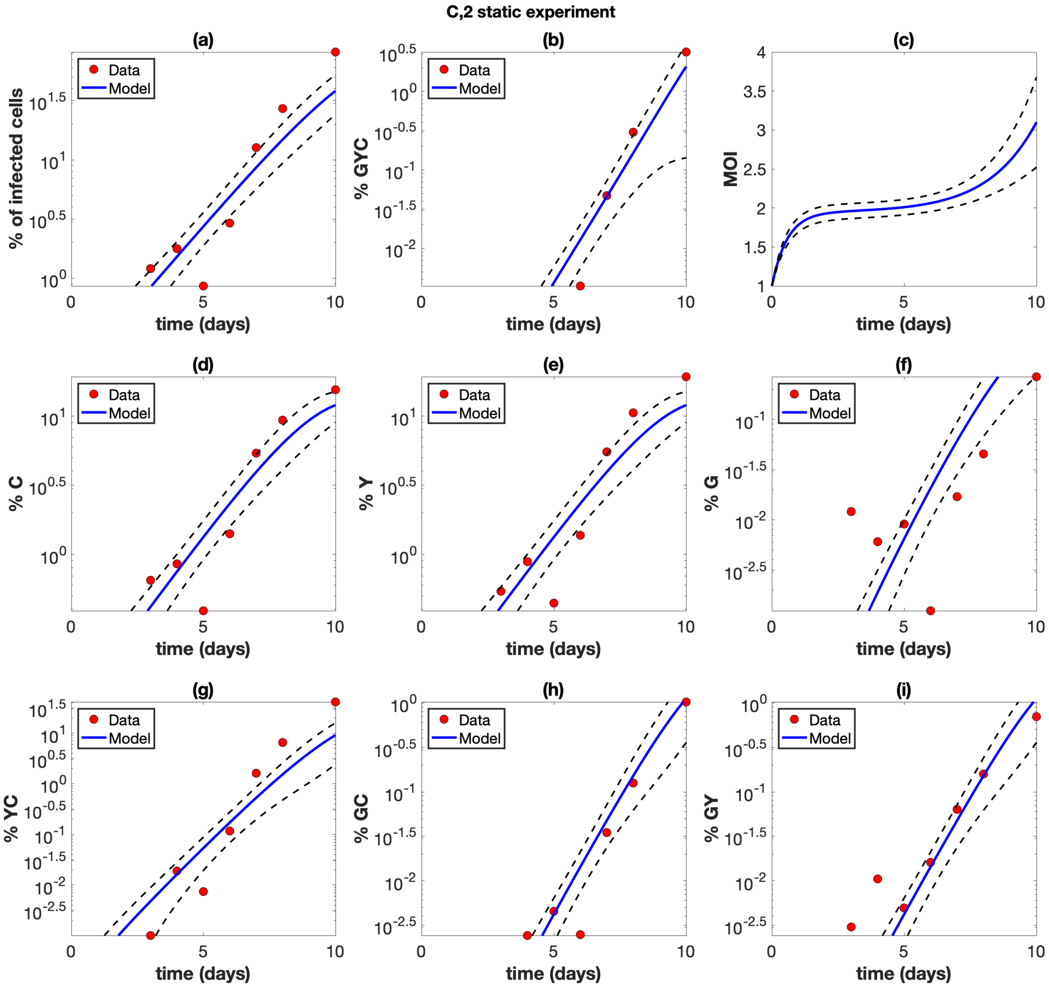


Figure S28: Static experiment C2. The experimental data (red circles) are presented with best fit curves from the model (blue lines). The mathematical model is described in Section 1 and the fitting procedure is described in Section 2. Best fit parameters are included in Table S2. The horizontal axis represents time (days). (a) The overall percentage of infected cells. (b) The percentage of cells infected with at least one copy of G, Y, and C. (c) The average multiplicity of infection (MOI) over all infected cells. (d) The percentage of cells infected with at least one copy of C. (e) The percentage of cells infected with at least one copy of Y. (f) The percentage of cells infected with at least one copy of G. (g) The percentage of cells infected with at least one copy of Y and C. (h) The percentage of cells infected with at least one copy of G and C. (i) The percentage of cells infected with at least one copy of G and Y. The dashed black lines represent the pointwise 95% prediction confidence bounds.

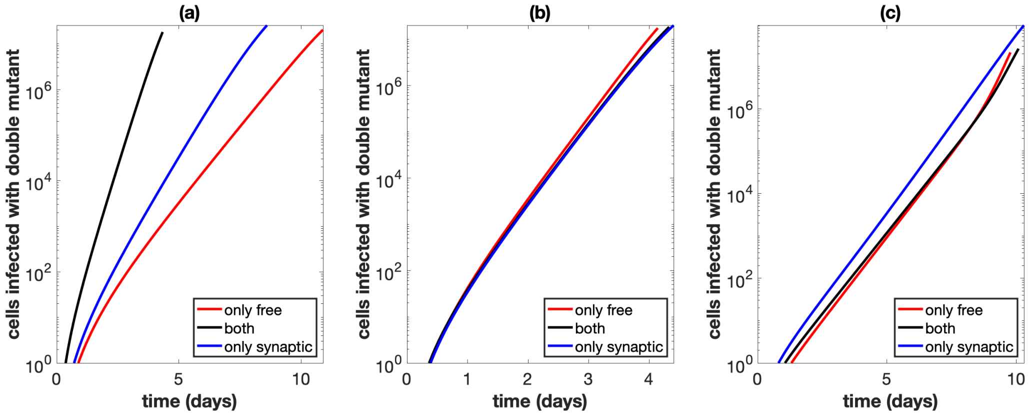


Figure S29: Similar to Figure 4 in the main text, using best fit parameters from experiment A1. Best fit parameters are included in Table S2. Total number of cells infected with at least one active copy of double mutant virus plotted against time. We assume we have $10^9$ initial cells, and run the simulation until the number of infected cells reaches 40% of this initial amount, while also assuming that uninfected cells do not divide. The combination of both transmission pathways is represented by the black lines. For the only free virus transmission case (red lines), synaptic transmission is turned off. For the only synaptic transmission case (blue lines), free virus transmission and reinfection are turned off. (a) Parameters are exactly as in the experiment. (b) The overall growth rate is the same across the three cases. (c) The overall growth rate is the same across the three cases and the simulation starts with only a single infected cell, which is coinfected with both single mutant strains.

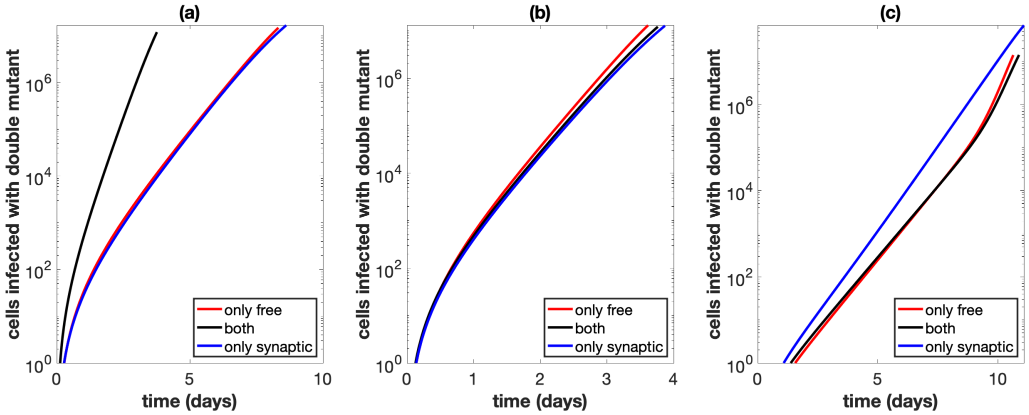


Figure S30: Similar to Figure 4 in the main text, using best fit parameters from experiment A2. Best fit parameters are included in Table S2. Total number of cells infected with at least one active copy of double mutant virus plotted against time. We assume we have $10^9$ initial cells, and run the simulation until the number of infected cells reaches 40% of this initial amount, while also assuming that uninfected cells do not divide. The combination of both transmission pathways is represented by the black lines. For the only free virus transmission case (red lines), synaptic transmission is turned off. For the only synaptic transmission case (blue lines), free virus transmission and reinfection are turned off. (a) Parameters are exactly as in the experiment. (b) The overall growth rate is the same across the three cases. (c) The overall growth rate is the same across the three cases and the simulation starts with only a single infected cell, which is coinfected with both single mutant strains.

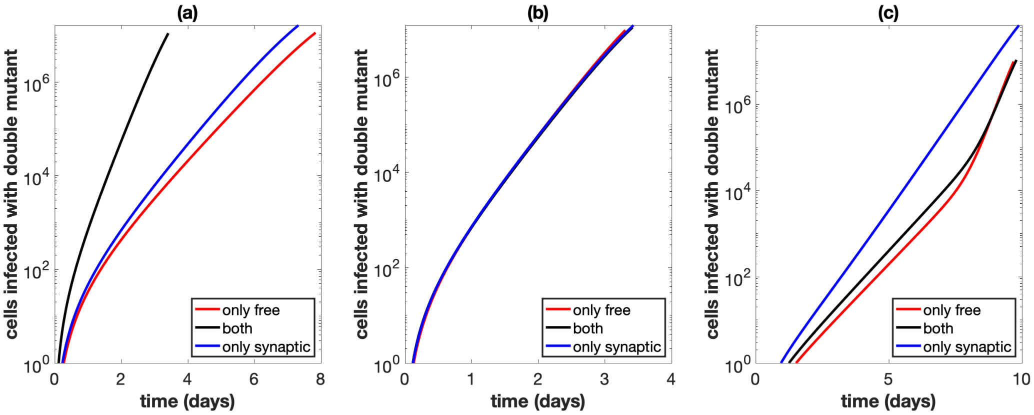


Figure S31: Similar to Figure 4 in the main text, using best fit parameters from experiment A3. Best fit parameters are included in Table S2. Total number of cells infected with at least one active copy of double mutant virus plotted against time. We assume we have $10^9$ initial cells, and run the simulation until the number of infected cells reaches 40% of this initial amount, while also assuming that uninfected cells do not divide. The combination of both transmission pathways is represented by the black lines. For the only free virus transmission case (red lines), synaptic transmission is turned off. For the only synaptic transmission case (blue lines), free virus transmission and reinfection are turned off. (a) Parameters are exactly as in the experiment. (b) The overall growth rate is the same across the three cases. (c) The overall growth rate is the same across the three cases and the simulation starts with only a single infected cell, which is coinfected with both single mutant strains.

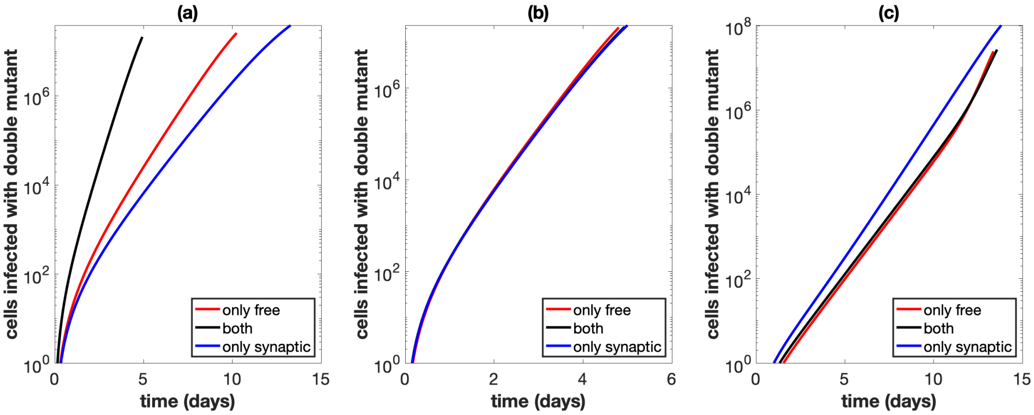


Figure S32: Similar to Figure 4 in the main text, using best fit parameters from experiment B1. Best fit parameters are included in Table S2. Total number of cells infected with at least one active copy of double mutant virus plotted against time. We assume we have $10^9$ initial cells, and run the simulation until the number of infected cells reaches 40% of this initial amount, while also assuming that uninfected cells do not divide. The combination of both transmission pathways is represented by the black lines. For the only free virus transmission case (red lines), synaptic transmission is turned off. For the only synaptic transmission case (blue lines), free virus transmission and reinfection are turned off. (a) Parameters are exactly as in the experiment. (b) The overall growth rate is the same across the three cases. (c) The overall growth rate is the same across the three cases and the simulation starts with only a single infected cell, which is coinfected with both single mutant strains.

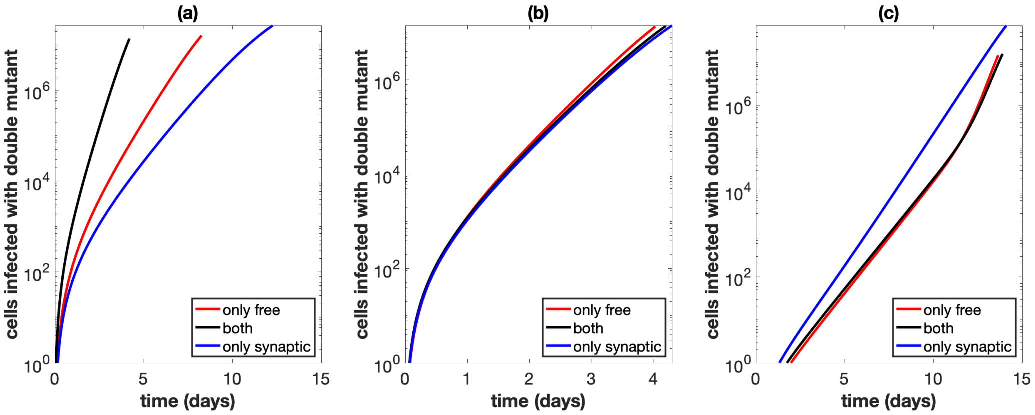


Figure S33: Similar to Figure 4 in the main text, using best fit parameters from experiment B2. Best fit parameters are included in Table S2. Total number of cells infected with at least one active copy of double mutant virus plotted against time. We assume we have $10^9$ initial cells, and run the simulation until the number of infected cells reaches 40% of this initial amount, while also assuming that uninfected cells do not divide. The combination of both transmission pathways is represented by the black lines. For the only free virus transmission case (red lines), synaptic transmission is turned off. For the only synaptic transmission case (blue lines), free virus transmission and reinfection are turned off. (a) Parameters are exactly as in the experiment. (b) The overall growth rate is the same across the three cases. (c) The overall growth rate is the same across the three cases and the simulation starts with only a single infected cell, which is coinfected with both single mutant strains.

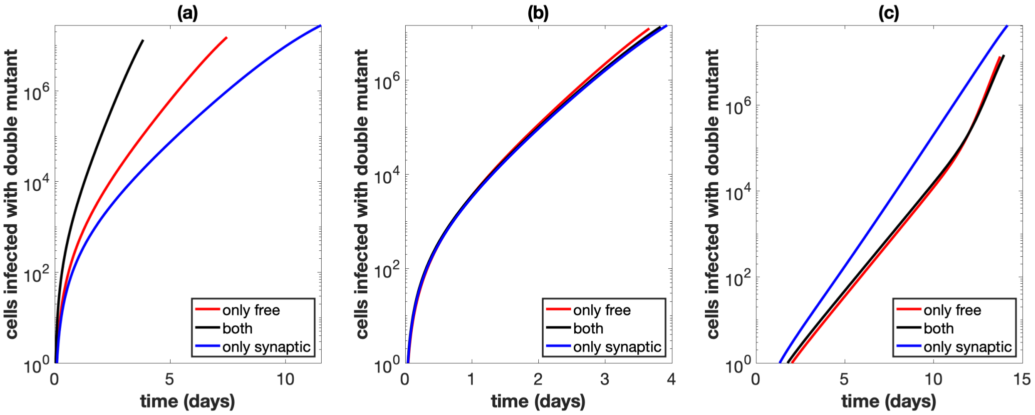


Figure S34: Similar to Figure 4 in the main text, using best fit parameters from experiment B3. Best fit parameters are included in Table S2. Total number of cells infected with at least one active copy of double mutant virus plotted against time. We assume we have $10^9$ initial cells, and run the simulation until the number of infected cells reaches 40% of this initial amount, while also assuming that uninfected cells do not divide. The combination of both transmission pathways is represented by the black lines. For the only free virus transmission case (red lines), synaptic transmission is turned off. For the only synaptic transmission case (blue lines), free virus transmission and reinfection are turned off. (a) Parameters are exactly as in the experiment. (b) The overall growth rate is the same across the three cases. (c) The overall growth rate is the same across the three cases and the simulation starts with only a single infected cell, which is coinfected with both single mutant strains.

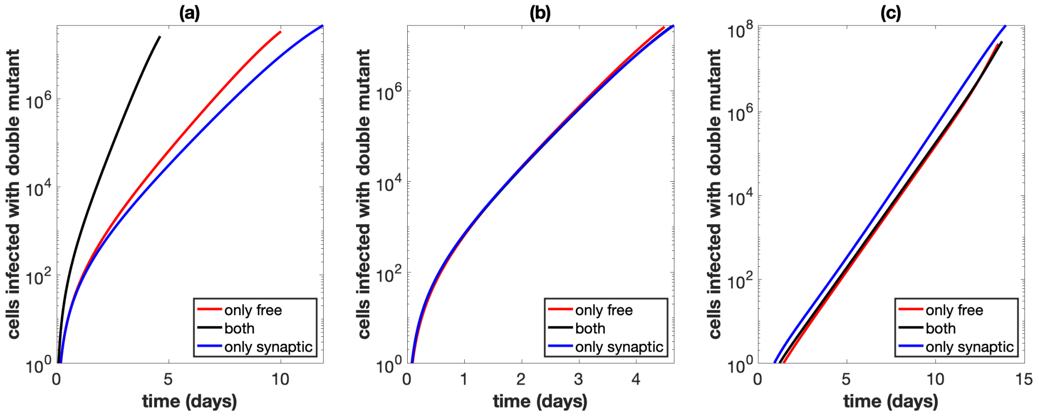


Figure S35: Similar to Figure 4 in the main text, using best fit parameters from experiment C1. Best fit parameters are included in Table S2. Total number of cells infected with at least one active copy of double mutant virus plotted against time. We assume we have $10^9$ initial cells, and run the simulation until the number of infected cells reaches 40% of this initial amount, while also assuming that uninfected cells do not divide. The combination of both transmission pathways is represented by the black lines. For the only free virus transmission case (red lines), synaptic transmission is turned off. For the only synaptic transmission case (blue lines), free virus transmission and reinfection are turned off. (a) Parameters are exactly as in the experiment. (b) The overall growth rate is the same across the three cases. (c) The overall growth rate is the same across the three cases and the simulation starts with only a single infected cell, which is coinfected with both single mutant strains.

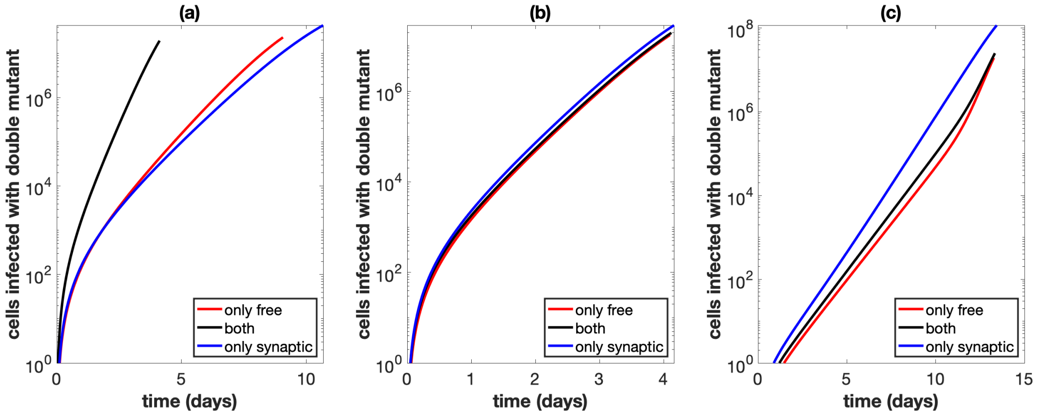


Figure S36: Similar to Figure 4 in the main text, using best fit parameters from experiment C2. Best fit parameters are included in Table S2. Total number of cells infected with at least one active copy of double mutant virus plotted against time. We assume we have $10^9$ initial cells, and run the simulation until the number of infected cells reaches 40% of this initial amount, while also assuming that uninfected cells do not divide. The combination of both transmission pathways is represented by the black lines. For the only free virus transmission case (red lines), synaptic transmission is turned off. For the only synaptic transmission case (blue lines), free virus transmission and reinfection are turned off. (a) Parameters are exactly as in the experiment. (b) The overall growth rate is the same across the three cases. (c) The overall growth rate is the same across the three cases and the simulation starts with only a single infected cell, which is coinfected with both single mutant strains.

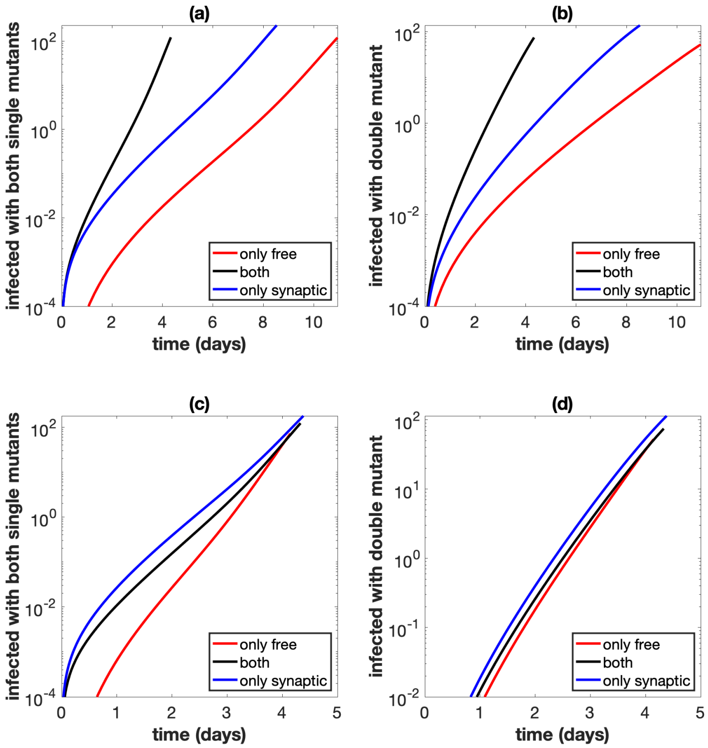


Figure S37: Similar to Figure 5 in the main text, using best fit parameters from experiment A1. Best fit parameters are included in Table S2. Simulations start with a small equal amount of cells singly infected with the wild type, and mutations are included. We assume we have $10^9$ initial cells, and run the simulation until the number of infected cells reaches 40% of this initial amount, while also assuming that uninfected cells do not divide. The combination of both transmission pathways is represented by the black lines. For the only free virus transmission case (red lines), synaptic transmission is turned off. For the only synaptic transmission case (blue lines), free virus transmission and reinfection are turned off. (a) Total number of cells infected with at least one copy of both single mutant strains. Parameters are exactly as in the experiment. (b) Total number of cells infected with at least one active copy of double mutant virus. Parameters are exactly as in the experiment. (c) Same as panel (a), but the overall growth rate is the same across the three cases. (d) Same as panel (b), but the overall growth rate is the same across the three cases.

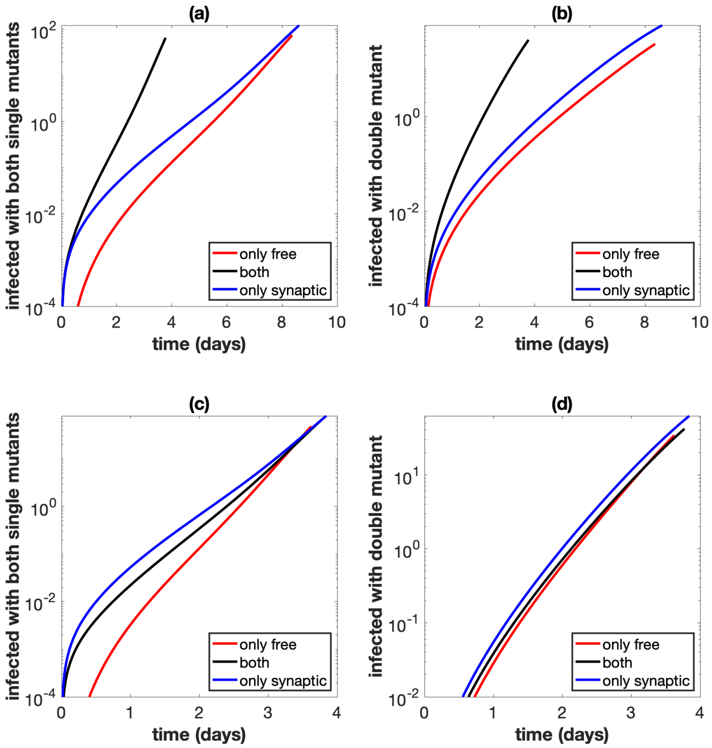


Figure S38: Similar to Figure 5 in the main text, using best fit parameters from experiment A2. Best fit parameters are included in Table S2. Simulations start with a small equal amount of cells singly infected with the wild type, and mutations are included. We assume we have $10^9$ initial cells, and run the simulation until the number of infected cells reaches 40% of this initial amount, while also assuming that uninfected cells do not divide. The combination of both transmission pathways is represented by the black lines. For the only free virus transmission case (red lines), synaptic transmission is turned off. For the only synaptic transmission case (blue lines), free virus transmission and reinfection are turned off. (a) Total number of cells infected with at least one copy of both single mutant strains. Parameters are exactly as in the experiment. (b) Total number of cells infected with at least one active copy of double mutant virus. Parameters are exactly as in the experiment. (c) Same as panel (a), but the overall growth rate is the same across the three cases. (d) Same as panel (b), but the overall growth rate is the same across the three cases.

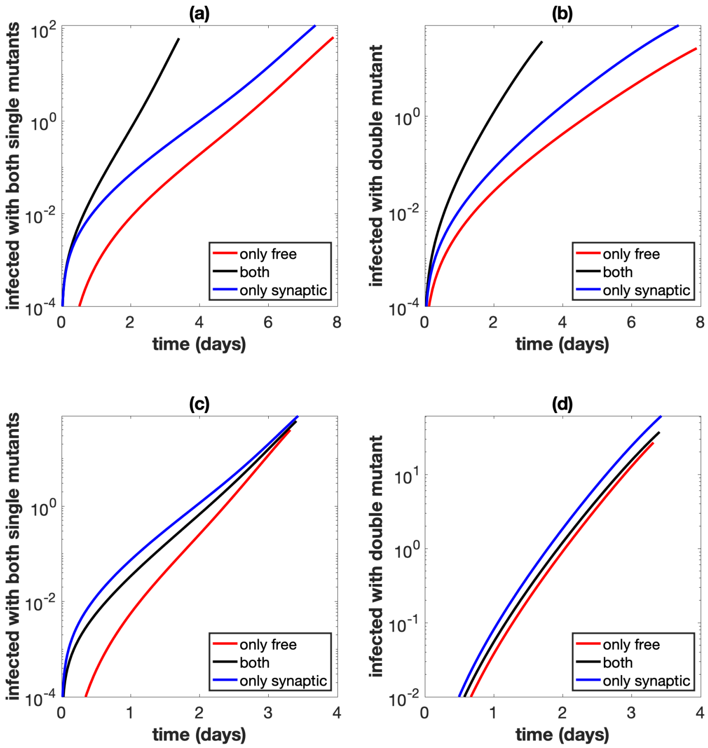


Figure S39: Similar to Figure 5 in the main text, using best fit parameters from experiment A3. Best fit parameters are included in Table S2. Simulations start with a small equal amount of cells singly infected with the wild type, and mutations are included. We assume we have $10^9$ initial cells, and run the simulation until the number of infected cells reaches 40% of this initial amount, while also assuming that uninfected cells do not divide. The combination of both transmission pathways is represented by the black lines. For the only free virus transmission case (red lines), synaptic transmission is turned off. For the only synaptic transmission case (blue lines), free virus transmission and reinfection are turned off. (a) Total number of cells infected with at least one copy of both single mutant strains. Parameters are exactly as in the experiment. (b) Total number of cells infected with at least one active copy of double mutant virus. Parameters are exactly as in the experiment. (c) Same as panel (a), but the overall growth rate is the same across the three cases. (d) Same as panel (b), but the overall growth rate is the same across the three cases.

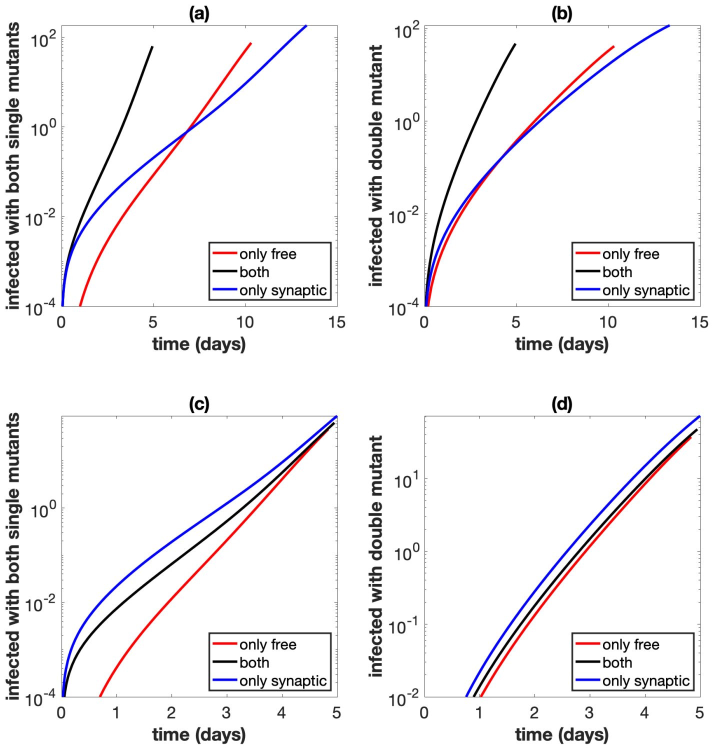


Figure S40: Similar to Figure 5 in the main text, using best fit parameters from experiment B1. Best fit parameters are included in Table S2. Simulations start with a small equal amount of cells singly infected with the wild type, and mutations are included. We assume we have $10^9$ initial cells, and run the simulation until the number of infected cells reaches 40% of this initial amount, while also assuming that uninfected cells do not divide. The combination of both transmission pathways is represented by the black lines. For the only free virus transmission case (red lines), synaptic transmission is turned off. For the only synaptic transmission case (blue lines), free virus transmission and reinfection are turned off. (a) Total number of cells infected with at least one copy of both single mutant strains. Parameters are exactly as in the experiment. (b) Total number of cells infected with at least one active copy of double mutant virus. Parameters are exactly as in the experiment. (c) Same as panel (a), but the overall growth rate is the same across the three cases. (d) Same as panel (b), but the overall growth rate is the same across the three cases.

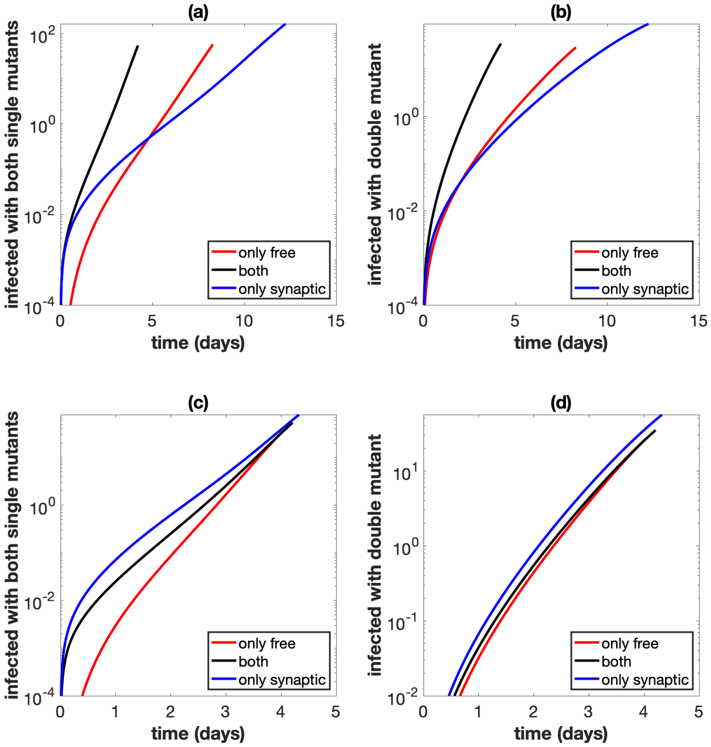


Figure S41: Similar to Figure 5 in the main text, using best fit parameters from experiment B2. Best fit parameters are included in Table S2. Simulations start with a small equal amount of cells singly infected with the wild type, and mutations are included. We assume we have $10^9$ initial cells, and run the simulation until the number of infected cells reaches 40% of this initial amount, while also assuming that uninfected cells do not divide. The combination of both transmission pathways is represented by the black lines. For the only free virus transmission case (red lines), synaptic transmission is turned off. For the only synaptic transmission case (blue lines), free virus transmission and reinfection are turned off. (a) Total number of cells infected with at least one copy of both single mutant strains. Parameters are exactly as in the experiment. (b) Total number of cells infected with at least one active copy of double mutant virus. Parameters are exactly as in the experiment. (c) Same as panel (a), but the overall growth rate is the same across the three cases. (d) Same as panel (b), but the overall growth rate is the same across the three cases.

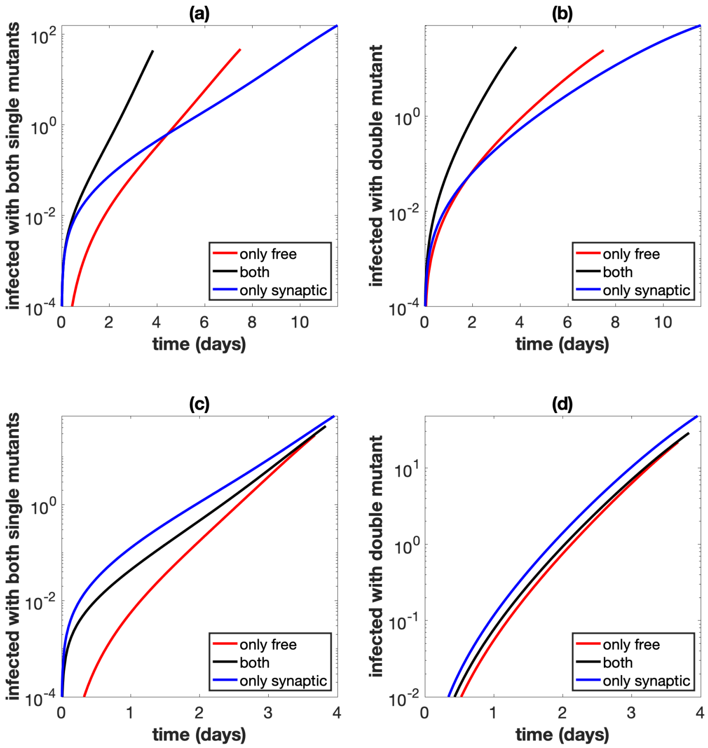


Figure S42: Similar to Figure 5 in the main text, using best fit parameters from experiment B3. Best fit parameters are included in Table S2. Simulations start with a small equal amount of cells singly infected with the wild type, and mutations are included. We assume we have $10^9$ initial cells, and run the simulation until the number of infected cells reaches 40% of this initial amount, while also assuming that uninfected cells do not divide. The combination of both transmission pathways is represented by the black lines. For the only free virus transmission case (red lines), synaptic transmission is turned off. For the only synaptic transmission case (blue lines), free virus transmission and reinfection are turned off. (a) Total number of cells infected with at least one copy of both single mutant strains. Parameters are exactly as in the experiment. (b) Total number of cells infected with at least one active copy of double mutant virus. Parameters are exactly as in the experiment. (c) Same as panel (a), but the overall growth rate is the same across the three cases. (d) Same as panel (b), but the overall growth rate is the same across the three cases.

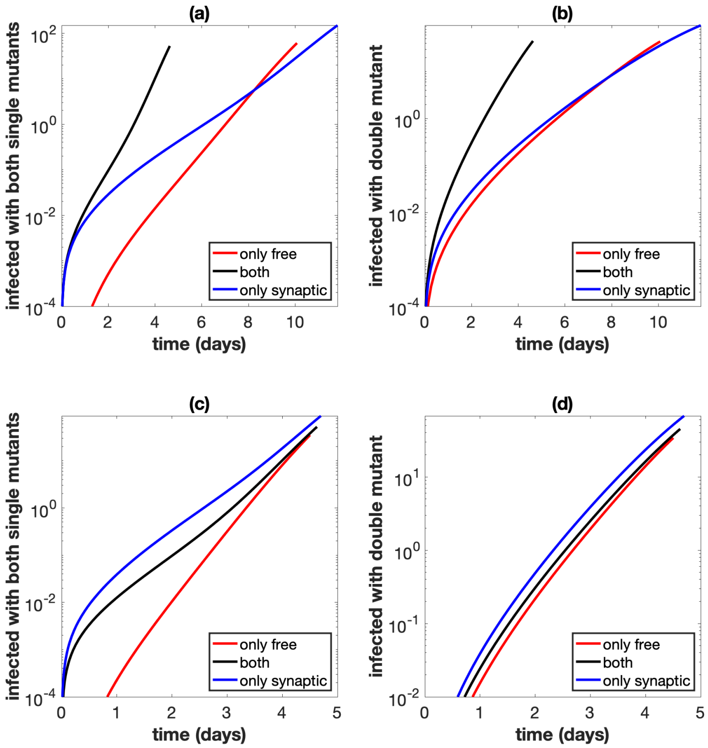


Figure S43: Similar to Figure 5 in the main text, using best fit parameters from experiment C1. Best fit parameters are included in Table S2. Simulations start with a small equal amount of cells singly infected with the wild type, and mutations are included. We assume we have $10^9$ initial cells, and run the simulation until the number of infected cells reaches 40% of this initial amount, while also assuming that uninfected cells do not divide. The combination of both transmission pathways is represented by the black lines. For the only free virus transmission case (red lines), synaptic transmission is turned off. For the only synaptic transmission case (blue lines), free virus transmission and reinfection are turned off. (a) Total number of cells infected with at least one copy of both single mutant strains. Parameters are exactly as in the experiment. (b) Total number of cells infected with at least one active copy of double mutant virus. Parameters are exactly as in the experiment. (c) Same as panel (a), but the overall growth rate is the same across the three cases. (d) Same as panel (b), but the overall growth rate is the same across the three cases.

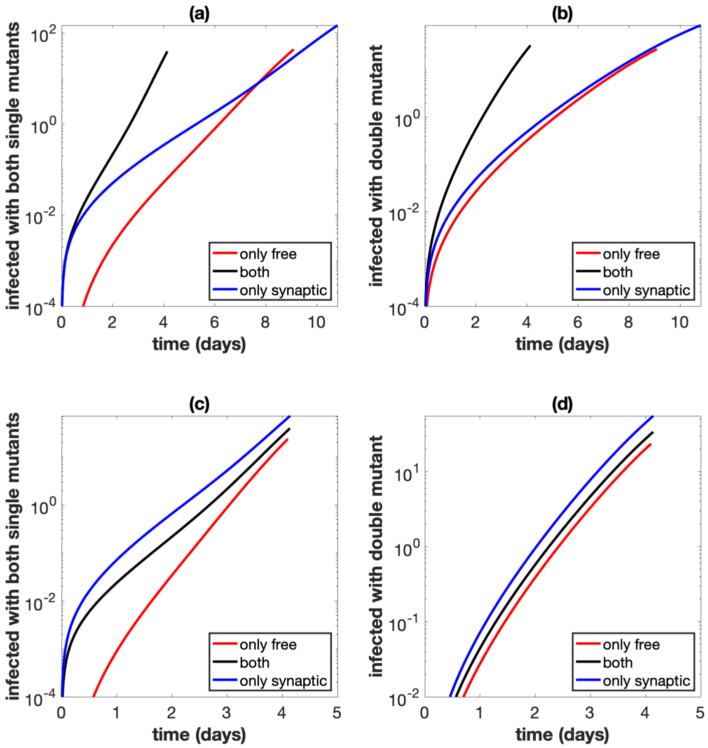


Figure S44: Similar to Figure 5 in the main text, using best fit parameters from experiment C2. Best fit parameters are included in Table S2. Simulations start with a small equal amount of cells singly infected with the wild type, and mutations are included. We assume we have $10^9$ initial cells, and run the simulation until the number of infected cells reaches 40% of this initial amount, while also assuming that uninfected cells do not divide. The combination of both transmission pathways is represented by the black lines. For the only free virus transmission case (red lines), synaptic transmission is turned off. For the only synaptic transmission case (blue lines), free virus transmission and reinfection are turned off. (a) Total number of cells infected with at least one copy of both single mutant strains. Parameters are exactly as in the experiment. (b) Total number of cells infected with at least one active copy of double mutant virus. Parameters are exactly as in the experiment. (c) Same as panel (a), but the overall growth rate is the same across the three cases. (d) Same as panel (b), but the overall growth rate is the same across the three cases.
